# Hyoid barbels as a key innovation triggering goatfish adaptive radiation

**DOI:** 10.64898/2026.09.08.750153

**Authors:** Laurent Mittelheiser, Josefin Stiller, Charlotte Koenig, Denis Baurain, Bruno Frédérich

**Author notes:** Biodiversity Genomics lab, Illinois Natural History Survey, Prairie Research Institute, University of Illinois at Urbana-Champaign, United States (Laurent Mittelheiser).

## Abstract

Key innovations can unlock ecological opportunities and redirect the evolutionary trajectory of entire lineages, yet few traits have been shown to trigger both hallmarks of a genuine adaptive radiation, rapid speciation and pronounced ecomorphological divergence, within a single lineage. This is especially true among coral reef fishes, where trophic novelties are routinely proposed as drivers of diversity but rarely tested against both criteria. Here, we show that goatfishes (Mullidae), a reef fish family having chemosensory hyoid barbels, meet this standard. First, macroevolutionary analyses of speciation dynamics reveal that goatfishes diversified at a markedly elevated rate relative to other Syngnathiformes. Using the most comprehensive phylogeny of the family to date, combining ultraconserved elements and mitochondrial data for 76 of the 103 recognized species, alongside the largest phenotypic dataset assembled for the group, we find that this elevated rate has been sustained since the family’s origin roughly 25 million years ago, rather than declining as ecological opportunity became filled. Head shape, the trait most closely tied to the foraging strategy enabled by hyoid barbels, diverged rapidly among genera early in the family’s history and has remained conserved ever since, while body shape and barbel microstructure evolved conservatively and pigmentation diversified later through repeated convergence. By opening access to previously untapped benthic prey resources and triggering a lasting burst of diversification, hyoid barbels emerge as a key innovation that reshaped the evolutionary trajectory of an entire reef fish lineage.

## Introduction

The lack of physical barriers and dispersive larval stages in marine environments likely sustain less allopatric speciation than in terrestrial or freshwater systems (Rocha & Bowen, 2008). Yet, marine ecosystems exhibit astonishing species richness (Mora et al, 2011; Costello and Chaudhary, 2017), and a prominent example is the reef-associated teleost fish assemblage, which comprises over 6,000 species (Bellwood et al., 2005, Brandl et al., 2018).

Adaptive radiation driven by key innovations probably underpinned reef fish diversification in sympatric conditions. Trophic innovations exemplify this phenotypic expansion: beak-like jaws for coral scraping in parrotfishes (Bellwood and Choat, 1990; Evans et al., 2023), protrusible jaws and hypertrophied lips for catching highly elusive prey in wrasses (Huertas and Bellwood, 2017; Wainwright et al., 2004), intramandibular joints for substrate grazing in surgeonfishes and angelfishes (Konow et al., 2008; Gibb et al., 2015), and small jaws for delicate polyp picking in butterflyfishes (Copus and Gibb, 2013). However, these adaptations have not been formally validated as key innovations satisfying two major criteria of an adaptive radiation scenario in a monophyletic group: (1) an early burst of speciation, and (2) substantial ecomorphological diversification among subclades (Schluter, 2000; Losos, 2010).

To date, clownfishes provide the only textbook example of adaptive radiation in tropical reef fishes, where obligate mutualism with sea anemones triggered both elevated speciation rates and ecomorphological diversification in host specificity and swimming abilities (Litsios et al., 2012; Mercader et al., 2025). Other clades show partial signatures of an adaptive radiation scenario. For example, the evolution of wrasses (Labridae) features pulses of species diversification linked to the rise of various trophic novelties (Brownstein et al., 2025), but these events occurred without concurrent ecomorphological variation within subclades. Conversely, the evolution of balistiform locomotion produced extensive ecomorphological diversification in triggerfishes (Balistidae), yet without elevated speciation rates (Dornburg et al., 2011). Detecting adaptive radiation is inherently challenging, as extinction can erase early diversification signals (Rabosky et al., 2018) and convergence can mask ecomorphological divergence (Frédérich et al., 2013).

Phylogenetically nested within Syngnathiformes, alongside seahorses and pipefishes, goatfishes (Mullidae) are singular among reef fish families despite being considered a typical component of reef assemblages (Bellwood, 1996). The family comprises 103 demersal species across six monophyletic genera: *Mulloidichthys*, *Mullus*, *Parupeneus*, *Pseudupeneus*, *Upeneichthys*, and *Upeneus* (Uiblein et al., 2024). Goatfishes differ from typical reef families through (1) distinctive hyoid barbels bearing sensory cells for detecting substrate-dwelling invertebrates (Gosline, 1984, Figure 1A), (2) large occupation of reef habitats and adjacent areas (Siqueira et al., 2023), and (3) recent origin (∼20 Ma; Nash et al., 2022) compared to most reef families (50-60 Ma; Bellwood et al., 2017).

**Figure 1.**
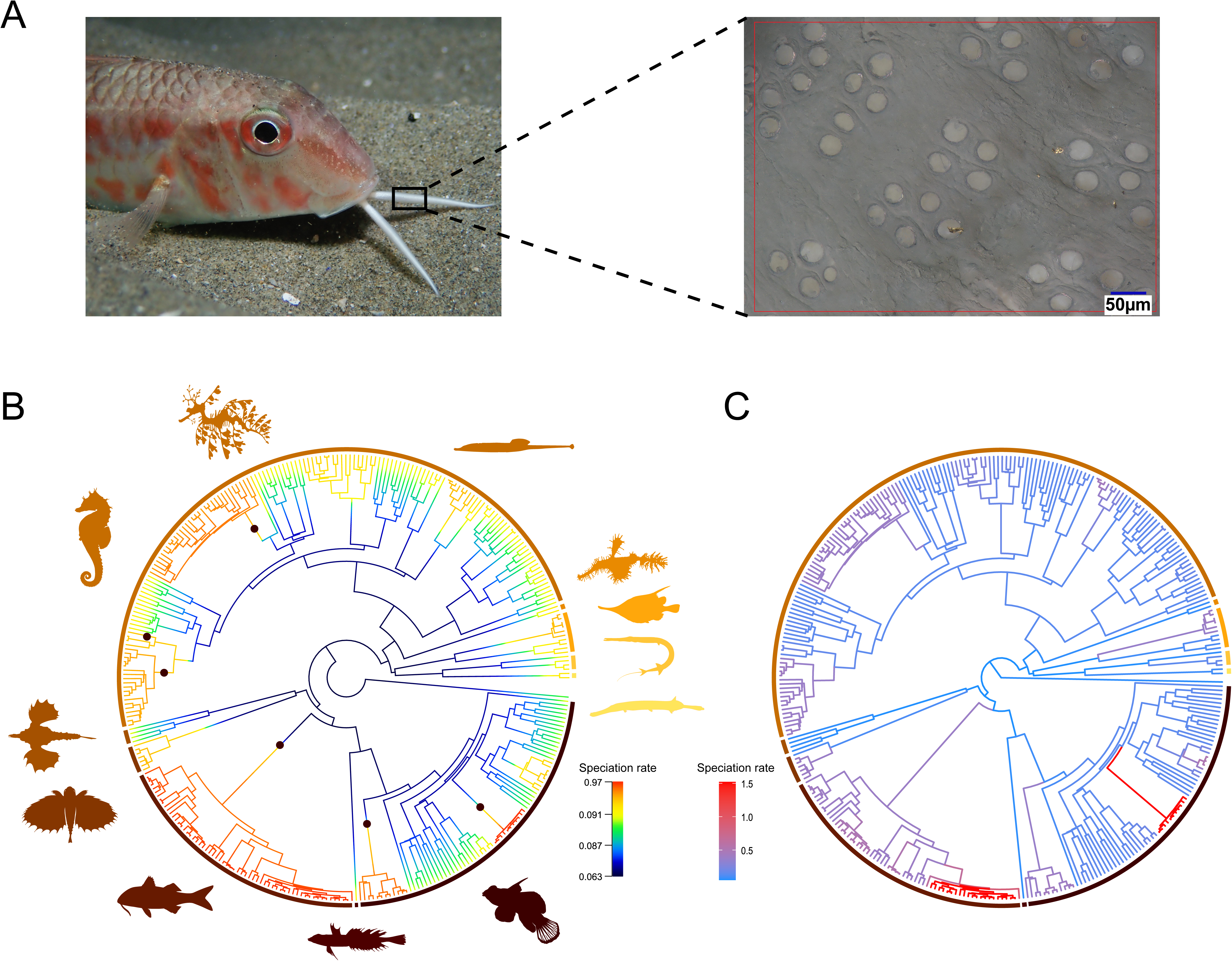
Goatfish’s hyoid barbels and diversification dynamics across Syngnathiformes. (A) Photograph of *Mullus surmuletus* illustrating the paired hyoid barbels, with a magnified optical microscopy view of taste buds on the barbel surface. (B) Time-calibrated phylogeny of Syngnathiformes (Santaquiteria et al., 2025) with branches colored according to speciation rate estimated using BAMM; black dots indicate the locations of significant rate shifts identified by the model. (C) The same phylogeny with branches colored according to speciation rate estimated using MiSSE. The Mullidae clade is at the bottom-left of both trees.

Given their relatively recent origin and distinctive hyoid barbels, we tested whether the diversification of Mullidae was shaped by an adaptive radiation (AR) with barbels acting as a key innovation by enabling access to untapped ecological niches and promoting diversification. To address this, we quantified diversification rates across Syngnathiformes, detecting a distinct acceleration in the Mullidae lineage. Building on this result, we reconstructed comprehensive phylogenomic hypotheses and assembled the largest phenotypic dataset for this family. Analyses of body morphology, head shape, barbels, and pigmentation patterns revealed early head shape diversification coinciding with trophic strategy variation. In contrast, body morphology and barbel structure evolved conservatively, and pigmentation patterns diversified intensively with a high degree of convergence. This complete signature of adaptive radiation, including rapid early lineage diversification, coupled with ecomorphological divergence functionally linked to trophic ecology, has not been previously documented in reef fishes. This suggests that key innovations represent an underappreciated driver of the exceptional diversity observed in coastal ecosystems.

## Results

### Highest rates of lineage diversification in Mullidae across Syngnathiformes

The first criterion of an adaptive radiation scenario requires that speciation be rapid relative to contemporaneous lineages (Schluter, 2000). We therefore employed two complementary approaches to test whether Mullidae exhibit elevated lineage diversification rates relative to other Syngnathiformes. We first applied Bayesian Analysis of Macroevolutionary Mixtures (BAMM; Rabosky, 2014) on the most recent and comprehensive phylogeny of Syngnathiformes (Santaquiteria et al., 2025) to detect diversification rate shifts. BAMM identified between three and nine plausible rate-shift configurations (posterior probabilities > 5%, ranging from 5.4% to 21%), with the six-shift configuration presenting the highest posterior probability. Crucially, all configurations revealed that the goatfish family evolved at a distinct tempo of diversification (Figure 1B), consistently showing some of the highest speciation rates. Then we used the Missing State Speciation and Extinction (MiSSE; Vasconcelos et al., 2022) framework to estimate branch-specific speciation rates and characterize diversification rate heterogeneity across the phylogeny, without any a priori assumption regarding the underlying character states (Figure 1C). MiSSE corroborated BAMM’s findings and recovered support for four alternative numbers of rate regimes (4, 6, 7 and 8 hidden states), each accounting for more than 5% of the relative Akaike weight (Table 1). Because no single model consistently emerged as the best fit based on AIC, we averaged diversification rates across models, and these results revealed that Mullidae indeed have one of the highest speciation rates across the order (up to 1.5). Together, these convergent results obtained through independent modelling frameworks satisfy the first criterion of an adaptive radiation scenario, demonstrating that goatfish speciation was markedly elevated relative to other Syngnathiformes lineages.

**Table 1.** Macroevolutionary diversification dynamics of Syngnathiformes and Mullidae. Posterior probabilities for the number of diversification shifts (BAMM) and AIC-based statistics for hidden state models (MiSSE). Models included in the final model-averaging set (AICw > 0.05) and the best-supported BAMM configurations are indicated in bold and marked with an sasterisk (*).

|  | BAMM |  | MiSSE |  |  |
| --- | --- | --- | --- | --- | --- |
| | Shifts | Posterior probability | Hidden states | ΔAIC | AIC weight |
| Syngnathiformes | 0 | 0.001 |  |  |  |
| | 1 | 0.02 | 1 | 69.74 | $3.18 \times 10^{-16}$ |
| | 2 | 0.05 | 2 | 12.56 | $8.29 \times 10^{-04}$ |
| | 3 | 0.11 | 3 | 13.65 | $4.81 \times 10^{-04}$ |
|  | 4 | 0.17 | **4** | **1.60** | **0.19\*** |
|  | **5** | **0.21\*** | 5 | 9.55 | 0.004 |
|  | 6 | 0.18 | **6** | **0** | **0.44\*** |
|  | 7 | 0.13 | **7** | **1.75** | **0.19\*** |
|  | 8 | 0.07 | **8** | **2.61** | **0.12\*** |
|  | 9 | 0.04 | 9 | 5.33 | 0.03 |
|  | 10 | 0.02 | 10 | 6.30 | 0.02 |
| Mullidae | **0** | **0.92\*** | 1 | **0.69** | **0.29\*** |
|  | 1 | 0.07 | 2 | **0** | **0.41\*** |
|  | 2 | 0.01 | 3 | **1.45** | **0.20\*** |
|  | 3 | 0.001 | 4 | **3.28** | **0.08\*** |
|  | 4 | 0.0001 | 5 | 5.63 | 0.02 |

### Phylogenomic inference of Mullidae and sustained high lineage diversification within the family

To further investigate diversification dynamics within the family, we first assembled the most comprehensive and robust phylogeny of Mullidae to date. Two genetic datasets were compiled from both *de novo* sequencing and mining of sequences from NCBI (Table S1). The first dataset was made of ultraconserved elements (UCEs; Faircloth et al., 2012). We produced UCEs for 56 species from 87 newly sequenced specimens to which we added the UCEs from 46 species collected from three published studies (Longo et al., 2017; Santaquiteria et al., 2021; Nash et al., 2022). This resulted in a UCE dataset for 69 species, including multiple individuals per species, with a total of 753 loci with 703,208 sites (Table S1). The secondary dataset was based on mitochondrial data, which consisted of 94 complete mitogenomes (including 61 de novo sequenced ones) across 54 species and enriched with 3,221 individual sequences of mitochondrial genes for 70 species retrieved from NCBI GenBank. This led to a mitochondrial dataset including 72 species (15 loci with 12,663 sites). The merging of these two datasets into a hybrid supermatrix led to a final taxonomic sampling of 76 species, representing all six of the valid genera and nearly 74% of the 103 currently recognized species in the family (Uiblein et al., 2024, Figure 2). The hybrid matrix had 715,871 sites, of which 109,016 were parsimony-informative and with 7.57% missing or ambiguous character states.

**Figure 2.**
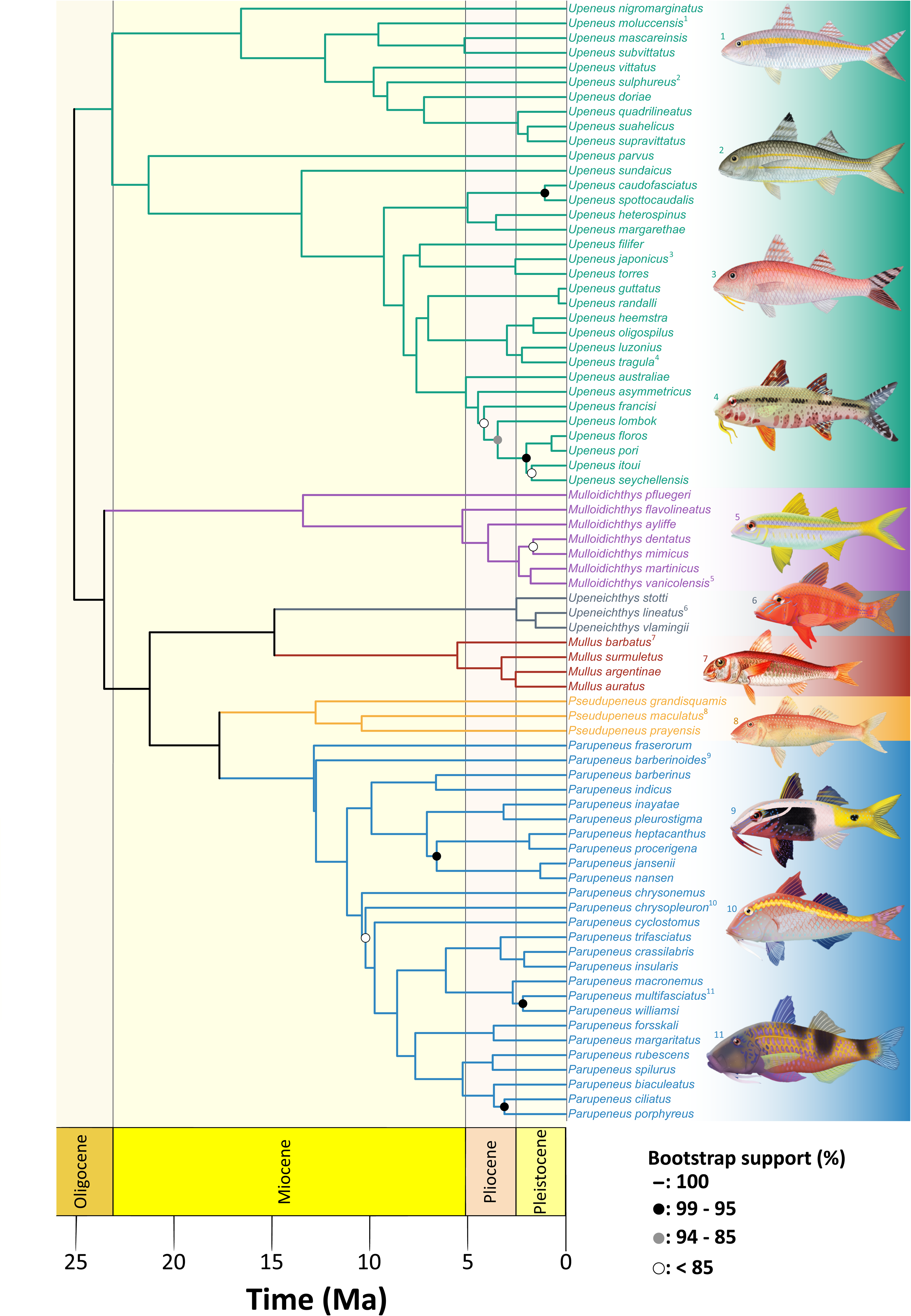
Dated phylogeny of Mullidae. Time-calibrated phylogeny of the 76 sampled Mullidae species inferred from combined UCE and mitochondrial data. Branches are colored by genus (*Upeneus*, teal; *Mulloidichthys*, purple; *Upeneichthys*, grey; *Mullus*, red; *Pseudupeneus*, orange; *Parupeneus*, blue). Node support values are indicated following the bootstrap support scale (filled circles, 100–95%; gray circles, 94–85%; open circles, <85%). Numbers next to species names link them to the corresponding representative illustrations shown at right. Fish illustrations courtesy of Kim In-young (fishillust.com) and MarineWise (Image © MarineWise https://marinewise.com.au), used with permission.

As subsequent diversification analyses required a time-calibrated phylogeny, we calibrated the hybrid supermatrix tree using a Bayesian approach in PhyloBayes (Lartillot et al., 2009). Two temporal constraints were applied to calibrate the autocorrelated relaxed molecular clock. The divergence between *Mullus* and *Upeneichthys* was constrained at 13 Ma using a fossil specimen from the Volhynian (early Sarmatian, Middle Miocene) of the Psheka River (western North Caucasus), representing the oldest known Mullidae with a complete skeleton (Carnevale et al. 2006). Assignment of this fossil to Mullidae relies on a combination of characters including small mouth, 24 vertebrae, 13 branched caudal-fin rays, two well-separated dorsal fins, ctenoid scales, and caudal skeleton morphology. Its referral to Mullus is further supported by skull bone composition, absence of teeth on the upper jaw, and six anal rays, while the single anal spine places it within the eastern Atlantic-Mediterranean clade of the genus (Carnevale et al. 2006). Divergence time estimation, calibrated using a prior centered on the estimate of Nash et al. (2022) (mean = 21.9 Ma), placed the origin of Mullidae at approximately 25 Ma. Alternative calibrations led to comparable results (see Methods).

Systematic relationships were greatly congruent between methodologies (Maximum Likelihood vs Bayesian Inference, Figure S1E) and among the three supermatrices (UCE, mitochondrial and hybrid, Figures S1A-C). When studied separately, the mitochondrial tree showed lower overall node support (Figure S1A and C). Hybrid and UCE trees were identical while some discrepancies were noted when compared to the mitochondrial tree (Figure S1B), indicating that the hybrid topology was predominantly driven by the UCE data. The topology connecting the six genera was totally congruent with the previous Mullidae phylogeny (Nash et al., 2022, Figure S1F) confirming *Mulloidichthys* as sister clade to *Upeneus,* followed by a split between clades of “*Mullus* + *Upeneichthys*” and “*Pseudupeneus* + *Parupeneus*”.

Beyond the elevated tempo of diversification detected at the scale of Syngnathiformes, adaptive radiation theory predicts that speciation rates within a radiating lineage may decline through time as ecological opportunity becomes progressively filled by accumulating diversity (Schluter, 2000; Rabosky and Lovette, 2008; Etienne et al., 2012). We tested this expectation by comparing the fit of five lineage diversification models to the Mullidae phylogeny, including constant-rate (Yule, birth-death), time-variable (exponentially varying), and diversity-dependent (linear and exponential) formulations (Table 2). The constant-rate, extinction-free Yule model received the strongest support (AICc weight = 0.58), followed by the constant-rate birth-death model, which also inferred a near-zero extinction rate (BDcst; AICc weight = 0.20, Table 2). Diversity-dependent models (DDL, DDX) and the time-variable model received comparatively little support (ΔAICc ≥ 3.24; Table 2). Notably, DDX model converged toward an unbounded carrying capacity (K, Table 2).

**Table 2.** Results of lineage accumulation models fitted with the DDD R package. df: degrees of freedom; Lambda: speciation rate; Mu: extinction rate; cst: constant through time; var_exp_: varying exponentially through time; DDL/DDX: diversity-dependent models with speciation and/or extinction rates varying linearly/exponentially with standing species richness, K: carrying capacity. Models are ranked by increasing AICc values. Bold values indicate models within ΔAICc ≤ 4 of the best-supported models.

| Models | df | Likelihood | AICc | $\Delta AICc$ | AICc weight | Lambda | Lambda slope | Mu | Mu slope | K |
| --- | --- | --- | --- | --- | --- | --- | --- | --- | --- | --- |
| **Yule** | **1** | **28.578** | **-55.10** | **0** | **0.58** | **0.15** | **0** | **0** | **0** | **NA** |
| **BDcst** | **2** | **28.577** | **-52.99** | **2.11** | **0.20** | **0.15** | **0** | **0** | **0** | **NA** |
| **DDL** | **3** | **29.100** |  | **3.24** | **0.12** | **0.21** | **NA** | **0.02** | **NA** | **236.97** |
| **DDX** | **3** | **28.692** | **-51.05** | **4.05** | **0.08** | **0.23** | **NA** | **0.01** | **NA** | **Inf** |
| BDvar<sub>exp</sub> | 4 | 28.583 | -48.60 | 6.50 | 0.02 | 0.39 | 0.04 | 0.38 | 0.11 | NA |

In the same way as we did for Syngnathiformes, we also performed BAMM and MiSSE to evaluate the presence of more complex dynamics of speciation and extinction in Mullidae (Figure 3). BAMM strongly supported a homogeneous diversification regime, with two credible configurations: zero shifts (posterior probability = 92%) or a single shift (posterior probability = 7.2%) (Table 1, Figure 3A). MiSSE similarly supported four plausible numbers of hidden states (1, 2, 3 and 4) with speciation rate ranging from 0.1 to 0.175, indicating an overall homogeneous distribution of rates but detected lower speciation in species with long branches such as *Upeneus parvus*, *U. nigromarginatus*, and members of the *Pseudupeneus* genus (Table 1, Figure 3B).

**Figure 3.**
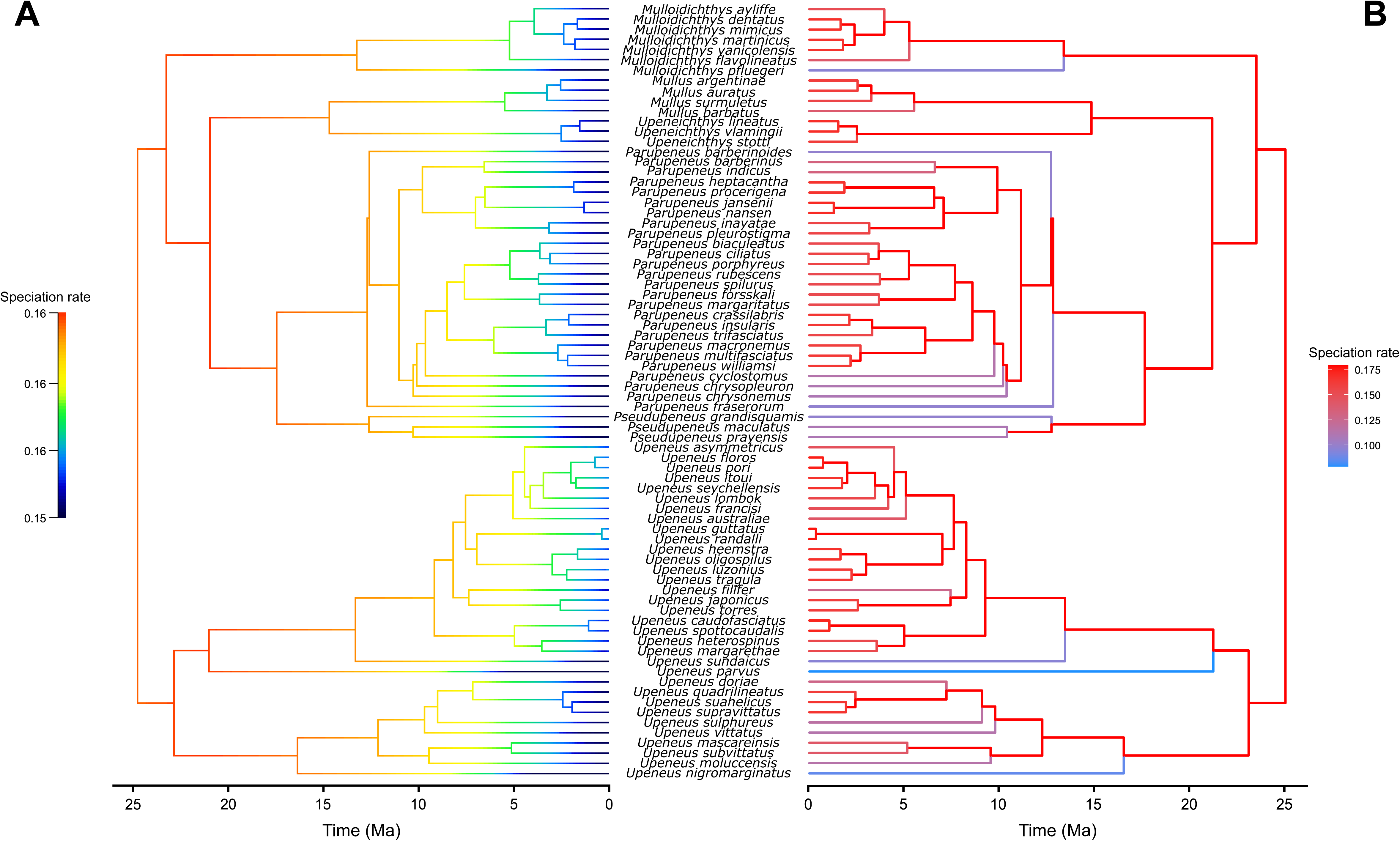
Diversification rates within Mullidae. Time-calibrated phylogeny of Mullidae with branches colored according to per-branch speciation rates estimated using (A) BAMM and (B) MiSSE.

### Contrasting patterns of phenotypic divergence among genera

The second criterion for an AR scenario is the diversification of subclades from a common ancestor into new, diverging adaptive zones (Schluter 2000). To test whether this criterion applies to the evolution of goatfishes, we examined the evolutionary patterns of four phenotypic trait sets: general body morphology, head shape, barbels, and pigmentation. These traits have deep ecological implications for locomotion, prey capture, and predator avoidance, and they have diverged within the family over time (Webb, 1984; Claverie and Wainwright, 2014; Brandl et al., 2015; Hench et al., 2022; Mittelheiser et al., 2025).

Visual exploration of phylomorphospaces, corroborated by Procrustes ANOVA, revealed contrasting patterns of morphospace occupation (Figure 4). Body morphology was characterized across 74 species using traditional morphometrics, built on 13 linear measurements and 11 related ratios (Figure 4A). The six genera were clearly differentiated in the morphospace defined by the first and the third principal components (PC), an among-genera variation supported by the Procrustes ANOVA (Figure 4A; R² = 0.27, F = 31.38, p < 0.001). Pairwise comparisons confirmed major variation in body morphology among most genera, but no significant differences were detected between the sister lineages *Parupeneus* and *Pseudupeneus* (p = 0.247), nor between *Mulloidichthys* and *Mullus* (p = 0.054) (Table S2A). Both PC1 and PC3 primarily captured variation in cephalic features. PC1 was driven by variations in head depth, snout length and body depth, while PC3 was associated with upper jaw length, head length and barbel width. Accordingly, *Upeneus* and *Mullus* species, showing positive PC1 scores, had slender head and body, and shorter snout compared to *Parupeneus* and *Pseudupeneus*. *Mulloidichthys* displayed intermediate body morphologies, whereas *Upeneichthys* had the deepest head and body, and the longest snout.

**Figure 4.**
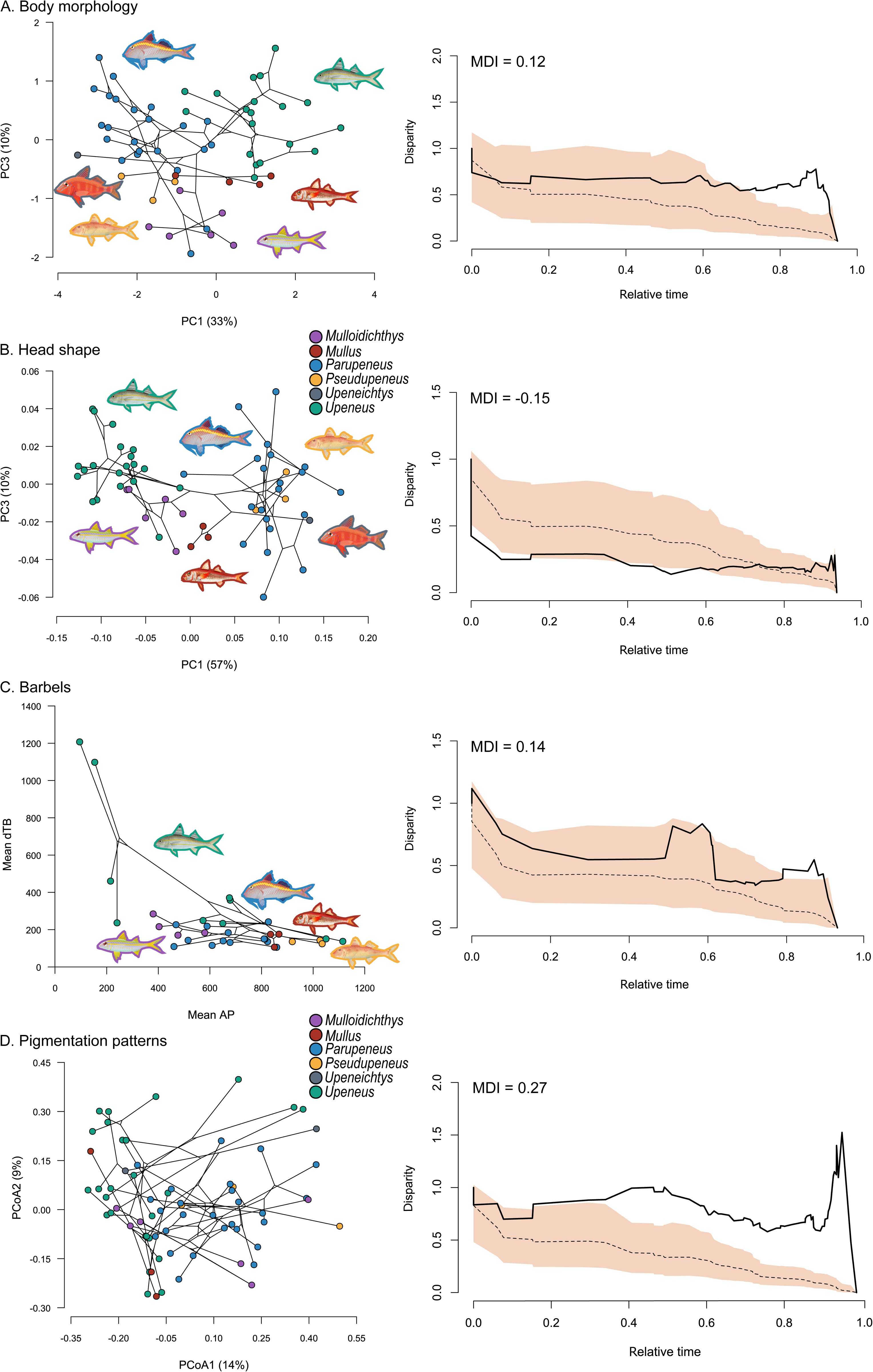
Morphological and pigmentation disparity through time in Mullidae. Left, phylomorphospace projections for (A) body morphology, (B) head shape, (C) taste bud density and pore area, and (D) pigmentation patterns, colored by genus. Right, disparity-through-time (DTT) plots for each trait, showing observed subclade disparity (solid line) relative to the expectation under a Brownian motion model of evolution (dashed line, with 95% confidence envelope). The morphological disparity index (MDI) is indicated for each trait.

Head shape variation in 69 species was quantified by landmark-based geometric morphometric methods. All the genera, except *Pseudupeneus* which was not distinguishable from *Parupeneus* in head shape, occupied different subspaces of the shape space defined by PC1 to PC3 (Figure 4B; among-genera Procrustes ANOVA: R² = 0.62, F = 122.01, p < 0.001; Table S2B). Notably, the factor *genus* explained a substantially larger proportion of variance for head shape (R² = 0.62) than for the overall body morphology (R² = 0.27). Similarly to the analysis of body morphology, PC2 had no power for the discrimination of genera but it explained variation within genera. *Upeneus* species, with the most negative PC1 scores, had larger and more anteriorly positioned eyes, slender elongated heads with shorter snouts, and more anteriorly positioned opercle and fin insertions (Figure S2). In contrast, *Upeneichthys* exhibited the smallest and most posteriorly positioned eyes, the deepest and shortest heads with the longest snouts, and the most anteriorly positioned opercle and fins (Figure S2). *Mullus* species had a central position in the shape space by showing intermediate head shapes (Figure 4B). On the one hand, *Mulloidichthys* species were relatively similar to the *Upeneus* group along PC1 (Figure 4B). On the other hand, *Mulloidichthys* differed from the other genera along PC3 by having more anterior eye and fin insertions, a smaller opercle, and a longer head with a shorter snout (Figure 4B, S2).

Barbel structure was assessed across 35 species capturing density and size of taste buds along with barbel length. More subtle but significant distinctions emerged in the morphospace of hyoid barbels (Figure 4C, among-genera Procrustes ANOVA: R² = 0.25, F = 2.5, p < 0.001; Table S2C). Mean density of taste buds (mean dTB) separated three *Upeneus* species (*U. moluccensis*, *U. suahelicus* and *U. sulphureus*) from other *Upeneus* and all remaining genera, with these species exhibiting markedly higher mean dTB values (Figure 4C). The mean area of pores (mean AP) further distinguished these three species, along with *U. vittatus* and *U. oligospilus*, which showed lower mean AP values compared to other *Upeneus* and representatives of other genera (Figure 4C). Along the mean AP axis, *Mulloidichthys*, *Mullus* and *Pseudupeneus* were distributed sequentially from lower to higher values (Figure 4C). Barbel length provided additional discrimination, with *Mullus* and *Pseudupeneus* species displaying longer barbels than *Upeneus* species (Figure S3).

Pigmentation patterns were scored across 82 species based on discrete traits capturing motif type, position, and distribution. Species distribution in the pigmentation pattern space revealed distinct but overlapping genus-level clusters (Figure 4D; among-genera Procrustes ANOVA: R² = 0.21, F = 23.95, p < 0.001, Table S2D). The first axis (PCoA1) separated a cluster comprising *Upeneus* and *Mullus* species (negative values) from another cluster including *Parupeneus* and *Pseudupeneus* species (positive values). Trait contribution vectors (Figure S4) clarified the drivers of this pattern with *Upeneus* and *Mullus* species in the negative PCoA1 region characterized by horizontal stripes and polychromatic body coloration, whereas *Parupeneus* and *Pseudupeneus* species occupying positive PCoA1 values typically displaying more uniform, monochromatic patterns with reduced ornamentation (Figure 4D, S4). The remaining genera, *Mulloidichthys* and *Upeneichthys,* were broadly distributed across the pigmentation pattern space, exhibiting larger diversity than the other genera. Neither PCoA2 nor PCoA3 revealed distinct patterns of space occupation (Figure S5).

### Phenotypic diversification of Mullidae

First, we inferred the association between the diversity of traits and the phylogenetic structure of goatfishes by testing phylogenetic signals using the multivariate extension of Blomberg’s K-statistic (Blomberg et al., 2003; Adams, 2014). All tests of phylogenetic signal were significant (body morphology, head shape and barbels: p < 0.0001; pigmentation patterns: p = 0.047) but the K-values varied across the phenotypic traits. Head shape and barbels exhibited the highest K values (1.14 and 1.10, respectively), indicating a pronounced phylogenetic structure. Body shape showed a moderate signal (K = 0.50), while pigmentation patterns displayed the weakest phylogenetic signal (K = 0.23).

Under the AR hypothesis, morphological disparity should be partitioned among major subclades, with each clade exhibiting strong ecomorphological differentiation (Schluter 2000). This prediction was tested by using disparity-through-time (DTT) plots combined with the morphological disparity index (MDI) calculated over the first two-thirds of relative time (Harmon et al., 2003). Head shape exhibited an early and sharp decline in subclade disparity, remaining consistently below the lower bound of the 95% confidence envelope (Figure 4B). This pattern was supported by a significant negative MDI (−0.15, p = 0.034) agreeing with the AR scenario. In contrast, pigmentation patterns showed an opposite trend with an empirical curve lying above the confidence envelope throughout the interval (Figure 4D), reflected by a positive MDI (0.27, p = 1). High phenotypic disparity maintained throughout the evolutionary history of goatfishes would be consistent with convergence across clades. Body morphology and barbel traits did not depart from the null Brownian motion model and remained within the confidence envelope (Figure 4A and C), with MDI values of 0.12 (p = 0.94) and 0.14 (p = 0.94), respectively. However, DTT plot interestingly revealed a transient increase in barbel disparity approximately midway through mullids’ evolutionary history, followed by sustained diversification levels (Figure 4C).

Finally, we aimed to test whether phenotypic traits evolved independently or not. We thus performed phylogenetically informed two-block PLS analyses, which revealed independent evolution among traits (Table 3). Only body morphology and hyoid barbels were slightly evolutionary integrated (r-PLS = 0.41, p = 0.045).

**Table 3.** Results from phylogenetically-corrected correlation test between pairwise combinations of the four datasets. Significant correlations are highlighted in bold.

|  | Head shape |  | Barbels |  | Pigmentation patterns (3PC) |  | Pigmentation patterns (32PCs) |  |
| --- | --- | --- | --- | --- | --- | --- | --- | --- |
|  | r-pls | p | r-pls | p | r-pls | p | r-pls | p |
| Body morphology | 0.28 | 0.18 | **0.41** | **0.045** | 0.36 | 0.14 | 0.58 | 0.07 |
| Head shape |  |  | 0.26 | 0.77 | 0.33 | 0.20 | 0.60 | 0.06 |
| Barbels |  |  |  |  | 0.28 | 0.67 | 0.49 | 0.90 |

## Discussion

In this study, we investigated the tempo and the mode of diversification in goatfishes, a singular reef fish family within Syngnathiformes. Despite evolving within a long-standing (Bellwood et al., 2015) and highly diverse (Fisher et al., 2015) marine ecosystem, goatfishes represent a relatively young radiation characterized by distinctive morphology and foraging ecology. By integrating complementary phenotypic datasets with new phylogenomic data, and using various comparative analyses including models of diversification, we show that the evolutionary history of goatfishes bears hallmark features of adaptive radiation, triggered by the emergence of a key innovation.

Our results support a scenario in which the emergence of one trait, the hyoid barbels, acted as a key innovation, enabling access to untapped niches and triggering a burst of diversification. This initial phase of the radiation was characterized by pronounced divergence in head morphology among genera, particularly along the snout elongation axis, a trait known to be tightly linked to trophic strategies (Wainwright and Bellwood, 2002). In parallel, this elevated tempo of lineage diversification, sustained relative to other Syngnathiformes since the origin of the family, indicates that goatfishes have maintained an unusually rapid pace of speciation throughout their evolutionary history. Together, these findings outline a rare but theoretically well-established evolutionary scenario in which a phenotypic innovation profoundly structures the pattern of diversification within a lineage.

### Ecomorphological diversification of goatfishes

Among all phenotypic datasets examined, head shape emerged as the primary axis of early diversification in goatfishes. Significant divergence in head morphology occurred near the root of the phylogeny, resulting in clearly distinct head shapes among genera (Figure 4B). This early differentiation is reflected by a negative MDI value and a strong phylogenetic signal, indicating that head shape divergence was both rapid and subsequently conserved through time.

Head shape variation is a major axis of morphological diversification in teleost fishes (Claverie and Wainwright, 2014), largely due to its tight association with feeding strategies (Costa and Cataudella, 2007; Aguilar-Medrano et al., 2011; Evans et al., 2019; Mittelheiser et al., 2022). Here, both analyses of head shape and body morphology revealed that one of the main axes of morphological variation across goatfishes relates to traits associated with head morphology: head length, head depth, snout length, eye size and its relative position, and barbel length and size. Various morpho-functional and ecological analyses support the hypothesis that head shape variation observed in goatfishes is associated with different feeding habits (Gosline, 1984; Mittelheiser et al., 2022). As suggested by a comprehensive trophic analysis including stable isotopes, species from the genera *Mulloidichthys* and *Upeneus*, which have short snout and large, anteriorly positioned eyes, forage for preys in the upper layers of the substrate (Mittelheiser et al., 2022). This ocular configuration where a large eye is positioned closer to the jaws has already been associated with foraging in complex habitats (Lisney et al., 2020). In contrast, *Parupeneus* species are thought to either search for prey in less complex habitats or probe deeper into the substrate (Mittelheiser et al., 2022, 2025). Their smaller and more posteriorly positioned eyes would facilitate the detection of prey from greater distances or allow them to probe deeper in the substrate while keeping their eyes above. Head shape variation in the genera *Mullus*, *Upeneichthys*, and *Pseudupeneus* also suggests divergent ecological strategies. *Upeneichthys* and *Pseudupeneus*, which occupy the same region of the phylomorphospace as *Parupeneus*, likely exhibit similar trophic habits. Conversely, the intermediate head shape observed in *Mullus* suggests a more generalist foraging ecology.

Beyond these differences in foraging strategies associated with head shape variation, trophic niche differentiation in goatfishes is further structured by spatial partitioning of foraging areas. In the coral reef systems of Toliara (SW Madagascar), clear cross-reef segregation has been demonstrated: *Upeneus* species are restricted to the outer reef shelf, *Mulloidichthys* species forage in the lagoon, and *Parupeneus* species occupy both habitats (Mittelheiser et al., 2025). Similar bathymetric and ecological partitioning occurs in the Mediterranean Sea, where *Mullus barbatus* is primarily found on deep muddy bottoms, whereas *M. surmuletus* inhabits shallower, coarser substrates (Golani, 1994; Lombarte et al., 2000). This fine-scale trophic and spatial segregation likely minimizes competition among goatfish species, which may further account for the absence of pronounced divergence in macrohabitat-related morphological traits (Mittelheiser et al., 2022, 2025). The limited morphological variation observed in the body plan of Mullidae (Nash et al., 2022) probably reflects the evolutionarily conserved demersal lifestyle and the lack of major adaptive shifts in locomotion strategies.

Barbels are central to goatfish biology, playing critical roles in trophic ecology and behavior (Gosline, 1984; Uiblein, 2007). Given that head shape, which serves as the morphological foundation for barbel attachment, exhibits the strongest interspecific variation and is tightly linked to trophic ecology, we anticipated corresponding divergence in finer scale barbel anatomy. Previous work in goatfishes from Madagascar revealed striking differences among species: *Upeneus* species, which exclusively inhabit outer reef shelves, possess densely distributed small taste buds, contrasting with the less dense, larger taste buds of *Mulloidichthys* and *Parupeneus* species, independent of their trophic habits (Mittelheiser et al., 2025). Conversely to our expectations, the present analysis revealed a relatively low variation in barbel microscopic characteristics at the family level, but this disparity showed a clear phylogenetic structure. This evolutionary pattern coincides with the emergence of a distinct *Upeneus* subclade including *Upeneus moluccensis, U. oligospilus, U. suahelicus, U. sulphureus* and *U. vittatus,* characterized by short barbels densely covered with small taste buds (Figure 4C and S3), the morphology previously documented in Madagascar (Mittelheiser et al., 2025). While this signal does not alter the broader evolutionary pattern, it suggests an additional axis of diversification related to sensory structures having occurred within an *Upeneus* lineage, warranting further investigation into the ecological specialization associated with this morphological divergence.

Pigmentation pattern evolution contrasted sharply with other morphological traits. Despite significant among-genus differences, pigmentation explained considerably less phenotypic variance (R² = 0.14) than head shape or body morphology. This low explanatory power, combined with weak phylogenetic signal and persistently high disparity relative to the null Brownian motion model (Figure 4D), suggests pronounced evolutionary lability and extensive phenotypic convergence across genera. This trend of pigmentation pattern evolution seems consistent across different reef fish families (Frédérich et al., 2026). The evolutionary flexibility and pervasive convergence in pigmentation patterns point to natural selection acting through ecological factors. Documented examples include adaptive dynamic coloration for camouflage during resting and foraging (Tosetto et al., 2021), and protective mimicry where *Mulloidichthys martinicus* resembles co-occurring Haemulidae (Krajewski et al., 2004) or Lutjanidae (Sikkel and Hardison, 1992) during mixed-species schooling. Even in the absence of documented sexual dimorphism, sexual selection could still operate through mechanisms such as mutual mate choice or species recognition, which do not require sex-specific coloration to shape ornamentation.

Interestingly, the divergence between *Parupeneus* and *Pseudupeneus* illustrates how phenotype can remain stable despite long-term biogeographic isolation. As sister clades, these genera diverged around 18 Ma (Figure 2) and show no geographic overlap across their entire distributions (Froese and Pauly, 2025), with *Pseudupeneus* occurring in the Atlantic and Eastern Tropical Pacific and *Parupeneus* in the coral-rich Indian and Central-West Pacific (Bellwood et al., 2015). This divergence time coincides with the closure of the Tethys Seaway (12-18 Ma, Steininger and Rögl, 1979; Adams et al., 1983; Rögl, 1998), consistent with a vicariant split isolating an Atlantic-associated lineage from its Indo-Pacific sister, as proposed for other Syngnathiformes lineages by Santaquiteria et al. (2021). Despite this prolonged isolation across ecologically distinct ocean basins, the two genera show complete phenotypic equivalence across all four trait axes examined (Figure 4). Given the early establishment and strong conservatism of head shape disparity documented above, this resemblance most likely reflects retention of the ancestral ecomorphology inherited from their common ancestor rather than independent evolution following the split. This pattern illustrates that an ecological niche established early in the radiation can persist over long timescales and across major biogeographic barriers, underscoring the strength of ecological niche conservatism in shaping goatfish phenotypes (Wiens and Graham, 2005). This interpretation is further supported by the strong phylogenetic signal detected in head shape, consistent with the general relationship between phylogenetic conservatism and niche stability (Losos, 2008).

### Hyoid barbels as a key innovation

Schluter (2000) delineated two defining criteria for adaptive radiation (AR). First, speciation should be rapid relative to contemporaneous lineages, reflecting the exploitation of novel ecological opportunity. Our BAMM and MiSSE analyses conducted at the scale of Syngnathiformes robustly and independently identified goatfishes as evolving at a markedly elevated tempo of diversification compared to their relatives (Figure 1), satisfying this first criterion. Within Mullidae itself, this elevated rate has been sustained since the origin of the family rather than following a pattern of decline. Our model comparison consistently favored constant-rate diversification, with the extinction-free Yule model receiving the strongest support and diversity-dependent models receiving little support (Table 2), while BAMM and MiSSE similarly supported a homogeneous diversification regime across the family (Figure 3). The absence of support for declining or diversity-dependent speciation, including the tendency of the exponential diversity-dependent model to converge toward an unbounded carrying capacity, indicates that density-dependence is not detectable in the accumulation of goatfish lineages. Despite representing the theoretically expected signature of niche filling during an adaptive radiation, this pattern may reflect the relatively recent origin of Mullidae (∼25 Ma) compared to other reef fish lineages (50–60 Ma; Bellwood et al., 2017), suggesting that multiple diversification mechanisms are still at work.

The second criterion of Schluter (2000) requires substantial phenotypic divergence in ecologically relevant traits across clades, driven by ecological opportunity through novel habitat access, competitive release, or key innovation (Losos, 2010). The observed early divergence in head shape satisfies this requirement, demonstrating pronounced among-genus differentiation in traits functionally linked to trophic ecology. Collectively, our results meet the two criteria of Schluter (2000), providing robust evidence that the evolutionary history of Mullidae was shaped by an adaptive radiation likely driven by the emergence of hyoid barbels.

Recently, Miller et al. (2023) proposed a conceptual refinement of key innovation terminology. They advocate restricting key innovation to its original definition, which means a trait enabling occupation of previously inaccessible ecological states, while introducing the concept of diversifying traits for innovations that additionally catalyze lineage diversification. This framework decouples ecological opportunity through acquisition of an innovation from its macroevolutionary consequences, recognizing these processes as potentially independent. The hyoid barbels of goatfishes satisfy both criteria. Substrate disturbance coupled with chemosensory prey detection through dense taste buds coverage certainly permits the exploitation of trophic niches in a unique way among reef fishes. To our knowledge, no other reef-associated fish taxa, nor indeed any fish, employ a functionally equivalent foraging strategy or possess barbels of comparable sensory complexity (Gosline, 1984). Additionally, the large head shape variation observed across subclades of Mullidae and associated eco-functional diversity even support the hypothesis that the emergence of barbels acted as a major trait novelty driving their adaptive radiation. Hyoid barbels therefore constitute a key innovation according to the definition of Miller et al. (2023).

Our analyses conducted at the scale of Syngnathiformes further revealed a burst of lineage diversification at the origin of Mullidae, robustly supported by both BAMM and MiSSE methods (Figure 1). The temporal coincidence between barbel emergence and elevated speciation rates indicates this trait functioned not only as an ecological enabler but also acted as a diversifying trait *sensu* Miller et al. (2023). Mullidae barbels therefore exemplify both classical key innovation frameworks emphasizing coupled ecological divergence and diversification (Hodges and Arnold, 1995; Hunter, 1998; Schluter, 2000) and the refined diversifying trait concept (Miller et al., 2023).

Unlike other reef fish lineages, which mainly diversify within structurally complex coral structures, goatfishes probably took advantage of their highly specialized barbels to transition into peripheral soft-bottom habitats. This adaptive shift indicates that barbel-mediated sensory innovation allowed goatfishes to occupy underutilized ecological spaces, such as sand and seagrass beds, effectively bypassing the intense competition found within the primary reef matrix.

### Temporal persistence of signatures attesting for an adaptive radiation

Several reef fish radiations have been documented and explored, including wrasses (Burress and Wainwright, 2019; Westneat et al., 2005), butterflyfishes (Bellwood et al., 2010; Cowman and Bellwood, 2011), surgeonfishes (Sorenson et al., 2013; Friedman et al. 2016), damselfishes (Frédérich et al., 2013, McCord et al., 2021), and hamlets (Hench et al., 2022) to cite a few. However, to the best of our knowledge, only clownfishes (Litsios et al., 2012; Mercader et al., 2025) and goatfishes exhibit the hallmarks of an adaptive radiation *sensu* Schluter (2000): an early and rapid lineage diversification with pronounced ecomorphological divergences among subclades. In both families, it is strongly supported that the ecological opportunity permitting this burst of diversification is the development of a key innovation: the physiological adaptation for living with giant stinging sea anemones for clownfishes and the hyoid barbels allowing new feeding habits in goatfishes.

Both families share relatively recent origins (clownfishes 25–30 Ma; goatfishes ∼25 Ma), potentially explaining why elevated diversification rates remain detectable in time-calibrated molecular phylogenies without yet showing signs of decline. Younger lineages may not yet have reached the ecological limits that would trigger a slowdown in diversification, whereas such limits, and the erosion of early diversification signatures by subsequent extinction, are more likely to have been reached in older radiations (Rabosky et al., 2018).

Beyond the effect of clade age discussed above, the sustained and elevated speciation rate observed within Mullidae may also reflect additional, non-exclusive mechanisms. First, speciation in Mullidae may be driven, at least in part, by allopatric or vicariant processes independent of ecological opportunity, as illustrated by the divergence between *Parupeneus* and *Pseudupeneus* following the closure of the Tethys Seaway. Second, the sustained rate of lineage accumulation could partly reflect speciation mechanisms linked to communication and species recognition, such as the labile and convergent evolution of pigmentation patterns documented across the family, which are not directly constrained by ecological opportunity. Together with clade youth, these complementary hypotheses suggest that the persistence of high speciation rates in Mullidae likely results from the combined action of ongoing ecological opportunity, historical biogeographic events and behaviorally driven diversification processes.

Time is probably not the sole factor obscuring past diversification patterns. Rapid and repeated evolutionary shifts across a limited set of ecological optima—such as those documented in damselfishes (Cooper & Westneat 2009; Frédérich et al., 2013)—when viewed over extended timescales (*i.e.*, a family’s entire evolutionary history), may appear as constant background diversification, effectively masking episodic bursts. When distributed across tens of millions of years these episodic bursts can be averaged into apparently stable rates. Robust detection of adaptive radiation in ancient lineages therefore necessitates total-evidence phylogenies integrating paleontological data with trait-based diversification models, as exemplified by recent analyses of cetaceans (Lloyd and Slater, 2021), Hymenoptera (Ronquist et al., 2012), and penguins (Gavryushkina et al., 2016).

### Hyoid barbels as a key innovation driving adaptive radiation

Together, our results establish goatfishes as a rare example of adaptive radiation among reef fishes, joining clownfishes as one of only two documented cases meeting both defining criteria of Schluter (2000) in the hyperdiverse coral reef ecosystem. The emergence of hyoid barbels, a chemosensory innovation enabling exploitation of previously inaccessible benthic prey resources, functioned as both a classical key innovation, opening access to new ecological opportunity, and a diversifying trait, additionally promoting an increase in speciation rate, triggering an early burst of speciation accompanied by pronounced and lasting ecomorphological divergence. By jointly meeting both hallmarks of adaptive radiation and satisfying the criteria of a key innovation, hyoid barbels constitute a textbook example of this evolutionary mechanism, illustrating how the acquisition of a single novel structure can reshape the diversification trajectory of an entire lineage.

## Methods

All analyses subsequent to phylogenomic reconstruction were conducted in R v4.4.0 (R Core Team, 2024).

### Taxon sampling and curation of morphological data

Tissue samples and specimens for molecular and morphological analyses were obtained through a combination of fieldwork, museum collections, and collaborative partnerships. Specimens were collected during fieldwork in Toliara and Nosy Be, Madagascar (May 2022), and examined during visits to the fish collections of the Muséum National d’Histoire Naturelle (MNHN, Paris, France; November 2021) and the National Museum of Natural History (NMNH, Washington, USA; January 2023). Across these three sources, barbel tissue, photographs, and morphometric data were collected following a consistent protocol. Further tissue samples were obtained on loan from natural history museum collections (see Table S3 for full specimen and voucher information). Additional ultraconserved element (UCE) and mitogenomic data were generated by co-author J.S. from whole-genome assemblies. These efforts were combined with published UCE and mitochondrial datasets (Longo et al., 2017; Santaquiteria et al., 2021; Nash et al., 2022) and sequences retrieved from NCBI GenBank (Clark et al., 2015), yielding a final dataset of 76 species representing all six genera and nearly 74% of the family’s recognized diversity.

### Phylogenomic reconstruction

#### Genetic data acquisition

We assembled a set of ultraconserved elements (UCEs) from three complementary sources to maximize taxonomic coverage and data quality. First, we obtained tissue samples representing 24 specimens across 15 species from museum collections (Table S1). Total genomic DNA was extracted using the DNeasy Blood & Tissue Kit (Qiagen) following the manufacturer’s protocol. Extracted DNA was sent to Arbor Biosciences for UCE library preparation, target enrichment using the Acanthomorphs 1Kv1 probe set (Faircloth et al., 2013), and Illumina sequencing. Second, UCE data for 49 species were extracted *in silico* from whole-genome draft assemblies (Stiller et al. in prep). To this end, DNA was extracted using the Thermo Scientific KingFisher instrument following the manufacturer’s protocol. Library preparation for short-read sequencing was carried out with Illumina PCR-free library preparation kits with Tagmentation and sequenced on an Illumina NovaSeq (150 PE) at the GeoGenetics Sequencing Core, University of Copenhagen. Sequencing data was trimmed from adapters and filtered for low-quality reads using fastp v.0.23.2 (Chen et al. 2023). Draft genome assemblies were generated using MaSuRCA v.4.0.9 (Zimin et al. 2013), which were then used to extract UCEs *in silico*, using the Acanthomorphs 1Kv1 probe set (obtained from https://www.ultraconserved.org/). Third, we incorporated published UCE datasets for 56 species from prior phylogenomic studies (Longo et al., 2017; Santaquiteria et al., 2021; Nash et al., 2022). Combined, these efforts yielded UCE data for 69 species, represented by 150 individuals.

Mitochondrial genomic data were compiled to expand taxonomic representation and provide independent corroboration of phylogenetic hypotheses. Complete mitochondrial genomes from 61 individuals representing 49 species were extracted *in silico* from raw reads of whole-genome sequencing data by first isolating potential mitochondrial reads from the read pool using mirabait (mira v.4.9.6) (Chevreux et al., 1999), which were then assembled and annotated using MitoFinder v.1.4.2 (Allio et al., 2020). If the reads extracted with mirabait did not yield complete mitochondrial genomes, the assembly was re-attempted directly from the raw read pool using only MitoFinder v.1.4.2. Each extraction yielded up to 15 mitochondrial genes per specimen (13 protein-coding genes: ATP6, ATP8, COX1, COX2, COX3, CYTB, ND1, ND2, ND3, ND4, ND4L, ND5, ND6; 2 ribosomal RNA genes: rrnL, rrnS), totaling 901 gene sequences. We further augmented sampling by systematically mining complete mitochondrial genomes and individual gene sequences from NCBI GenBank (Clark et al., 2015), identifying 35 complete mitogenomes (15,686–17,984 bp) and 4,405 individual gene sequences (111–1,294 bp). The dataset was carefully curated to exclude entries with uncertain taxonomy (e.g., *Mulloidichthys* sp., *Mulloidichthys* cf. species), environmental DNA sequences, non-accepted taxa (e.g., *Parupeneus bifasciatus*, *Mullus auriflama*, *Upeneichthys porosus*), and two genomes flagged as “UNVERIFIED” (GenBank accessions OK554512.1, OK554511.1) that were already represented in other datasets. After curation, we retained 33 complete mitogenomes and 4,164 validated individual sequences, of which 3,221 corresponded to actual mitochondrial genes (the remaining 943 primarily representing D-loop fragments from population genetic studies). Individual gene sequences represented eight genes: ATP6, ATP8, COX1, COX2, CYTB, ND2, rrnL, and rrnS. The final mitochondrial dataset encompassed 72 species.

#### UCE data processing and quality control

All UCE data processing was performed using the PHYLUCE pipeline v1.7.3 (Faircloth, 2016). Raw sequencing reads from Arbor Biosciences and NCBI-downloaded datasets (Santaquiteria et al., 2021; Nash et al., 2022) were cleaned to remove adapter contamination and low-quality bases using Illumiprocessor v2.10 (Faircloth, 2013), a wrapper for Trimmomatic (Bolger et al., 2014), with default settings. Cleaned reads were assembled into contigs using SPAdes v3.14.1 (Prjibelski et al., 2020). UCE loci were identified and extracted from assembled contigs using the Acanthomorphs 1Kv1 probe set. Sequences were aligned using MAFFT v7.475 (Katoh and Standley, 2013) through the *phyluce_align_seqcap_align* function with the --incomplete-matrix argument to accommodate loci absent in some specimens. Data from Longo et al. (2017) entered the pipeline at this stage as available reads were already assembled. Following recommendations for clades with recent divergence times (<30–50 Ma; Faircloth, 2016), only edge trimming was performed. This approach is appropriate for Mullidae, which originated ∼20 Ma (Nash et al., 2022). During trimming, UCE loci represented by fewer than three specimens were automatically discarded, resulting in exclusion of 63 loci. We evaluated UCE matrix completeness at multiple taxon-occupancy thresholds to balance phylogenetic information content against missing data. Completeness matrices were generated, testing thresholds of 70% and 90% taxon occupancy. The 70% completeness matrix contained 969 loci encompassing 889,190 bp (4.1% missing data), while the 90% matrix included 753 loci encompassing 703,208 bp (4.0% missing data). Both supermatrices were concatenated and phylogenomic trees were inferred for both matrices to assess potential topological differences arising from threshold selection, using IQ-TREE v1.6.12 (Nguyen et al., 2015) with 1,000 ultrafast bootstrap replicates (Hoang et al., 2018) and automatic model selection via ModelFinder (Kalyaanamoorthy et al., 2017). Tree comparison revealed minimal topological differences (data not shown), leading to selection of the 90% completeness threshold for subsequent analyses in order to optimize data quality while maintaining broad taxonomic representation.

Positional congruence within alignments was evaluated to identify and remove poorly aligned or highly variable sites that could introduce phylogenetic noise. We used Spruceup v2024.7.22 (Borowiec, 2019), which evaluates positional congruence by computing distances among sequences and discards portions exceeding distance cutoffs, replacing overly variable bases with gaps. We tested distance cutoffs ranging from 0.70 to 0.95 (0.70, 0.73, 0.75, 0.77, 0.80, 0.83, 0.85, 0.87, 0.90, 0.95) based on Weibull-min distribution criteria, with distances computed using Jukes-Cantor correction. Spruceup parameters included a window size of 20 bp with 10 bp overlap, allowing the window to slide by 10 bp at each step. The 90% UCE matrix tree, provided as a cladogram rooted on *Upeneus*, served as a guide tree. Distance distribution plots for each sequence revealed smooth exponential decay patterns, with most windows exhibiting extremely low distances among taxa (<0.01), indicating both limited outliers and high topological consistency across the dataset. Most distributions tailed off below 0.20, so we applied a manual cutoff of 0.20 to flag and remove windows containing highly variable positions, thereby preserving the overall informativeness of the alignment. Quality control assessment revealed that Spruceup removed 1,000 to 14,030 positions per specimen. Gaps introduced by Spruceup represent windows of uncertain nucleotide-level homology rather than true biological gaps (insertions/deletions). To avoid subsequent analytical artifacts, we converted Spruceup-introduced gaps (“-”) to missing character states (“?”) using the custom script *missify-ali.pl* (D. Baurain), which compared pre- and post-Spruceup matrices to identify replaced positions included in removed windows.

Species-level and genus-level monophyly were systematically verified using check-mono.pl (D. Baurain, custom script). Non-monophyletic sequences were evaluated using a semi-automated decision tree based on: (1) phylogenetic placement consistency, (2) sequence quality metrics (length, gap frequency, nucleotide ambiguity), (3) taxonomic reliability of voucher specimens, and (4) potential contamination or misidentification. This process identified nine specimens across seven species for exclusion.

For species represented by multiple sequences, we implemented a selection strategy to retain one representative sequence per species while maximizing data completeness. The 90% completeness tree was rooted on *Upeneus* following established phylogenetic hypotheses (Longo et al., 2017; Santaquiteria et al., 2021; Nash et al., 2022) using format-tree.pl (Bio::MUST::Core v0.250380, D. Baurain; https://metacpan.org/dist/Bio-MUST-Core). We tested four parameter combinations using *scaphoid.pl* (D. Baurain, custom script), a modern re-implementation of SCaFoS (Roure et al., 2007): (1) discard based on sequence length, where only the longest (*i.e.*, most complete) sequence remained, (2) discard based on distance-to-root, where only the sequence inducing the shortest distance to the root in the trees computed in the previous step remained, (3) chimerization based on sequence length, where available duplicates were chimerized with duplicates being ranked according to sequence length and (4) chimerization based on distance-to-root, where available duplicates were chimerized with duplicates being ranked according to descending distance to root in the trees computed in the previous step. Results showed no topological differences among the four strategies, but the distance-to-root method (phylogenetic path from leaf to root) provided superior branch support values, and chimerization significantly increased completeness compared to simple sequence discard. Therefore, chimerization ranked by phylogenetic distance-to-root was retained, creating chimeric consensus sequences by ranking conspecific sequences by phylogenetic proximity to root, then merging to fill missing character states and maximize completeness while preserving phylogenetic accuracy.

#### Mitochondrial data processing and quality control

Mitochondrial gene sequences from whole-genome extractions and NCBI downloads were organized into 15 gene-specific matrices, each containing sequences from all specimens for the corresponding gene. Protein-coding genes (ATP6, ATP8, COX1, COX2, COX3, CYTB, ND1, ND2, ND3, ND4, ND4L, ND5, ND6) were aligned individually using MACSE v2.07 (Ranwez et al., 2018) with codon-aware alignment based on amino acid translation. The MACSE alignSequences function was run with parameter -gc_def 2, specifying vertebrate mitochondrial genetic code (Ranwez et al., 2021). Ribosomal RNA genes (rrnL, rrnS) were aligned using MAFFT v7.475 with --auto parameter for automatic algorithm selection. Individual mitochondrial gene sequences from NCBI (3,221 sequences after initial curation) were aligned to corresponding gene alignments from complete mitogenomes using TwoScalp v0.243240 (D. Baurain; https://metacpan.org/dist/Bio-MUST-Apps-TwoScalp) with --fragments and --keep-length options enabled. This approach integrates fragmentary sequences into existing alignments while maintaining existing alignment length and structure.

Alignment completeness was systematically improved by filtering sites and sequences based on site-wise gap frequency using ali2phylip.pl (Bio-MUST-Core v0.250380, D. Baurain; https://metacpan.org/dist/Bio-MUST-Core). We tested filtering thresholds ranging from 0.1 to 0.9 in 0.1 increments for both site and sequence screening. The site threshold (−-max parameter) specified the minimum proportion of non-gapped (and non-missing) characters required at a site relative to total sequences. The sequence threshold (−-min parameter) specified minimum relative sequence length compared to the longest sequence (excluding gaps). Based on completeness statistics, we selected --max=0.4 (removing sites with >60% gaps) and --min=0.6 (removing sequences <60% of maximum length) to balance data retention and quality. For protein-coding genes, parameters --keep-codons and --codon-max=0.1 ensured that site-wise masks respected codon structure, preventing the introduction of frameshifts. Before applying --keep-codons, each alignment was verified to begin with a start codon. Preliminary phylogenetic trees were constructed for each of the 15 mitochondrial genes using IQ-TREE v1.6.12 with automatic model selection via ModelFinder (−m MFP argument) and 1,000 ultrafast bootstrap replicates. These gene trees served to identify non-monophyletic sequences and evaluate alignment quality, serving as additional topology controls. Species-level and genus-level monophyly were verified using check-mono.pl (Bio::MUST::Core). Non-monophyletic sequences were evaluated using the same decision tree framework as for UCE data. We excluded 119 sequences from 29 species (117 COX1, 1 ATP8, 1 ND6) from a total of 3,221 sequences across 15 genes. For species and/or specimens with multiple mitochondrial sequences, we implemented a hierarchical two-step selection strategy using *scaphoid.pl* on individual gene trees, re-estimating these trees from the sequences retained at each step to provide accurate distance-to-root rankings. First, at the specimen level, we tested four parameter combinations: (1) discard based on sequence length, (2) discard based on distance-to-root from trees computed during the previous step, (3) chimerization based on sequence length, and (4) chimerization based on distance-to-root from trees computed during the previous step. Results showed no major differences in sequence completeness between discard and chimerization methods, nor between sequence-length and distance-to-root ranking. We selected the discard/sequence-length method to retain only the longest (i.e., most complete) sequence per individual specimen or complete mitogenome, maximizing sequence length while avoiding incorporation of potentially low-quality nucleotides from shorter sequences. Second, at the species level, we applied the same four parameter combinations to per-gene trees rebuilt from the specimen-level-retained sequences. Selection criteria prioritized sequence completeness and phylogenetic resolution (topology quality, branch support values). The chimerization/distance-to-root method was retained as it produced the most complete sequences while maintaining a strong phylogenetic signal. This approach created chimeric consensus sequences by ranking conspecific sequences by phylogenetic proximity to root (shortest path from leaf to root in the resulting rooted tree), then merging sequences to fill missing character states and maximize data completeness while preserving phylogenetic accuracy. Resulting per-gene matrices were concatenated into candidate supermatrices for each parameter combination, and phylogenetic trees were rebuilt from each to confirm topological consistency before final matrix selection.

The final concatenated mitochondrial supermatrix contained all 15 genes with one sequence per species, totaling 12,663 bp across 72 species. A preliminary maximum likelihood tree was constructed using IQ-TREE with the same parameters as single-gene trees to validate topology and data quality before partitioning optimization.

#### Partitioning schemes and model selection

We used the sliding-window approach and entropy site characteristic (SWSC-EN), a partitioning scheme proposed for UCEs, to determine the best fit scheme for the flanks and cores across each locus in the 90%. Optimal substitution models were determined for each partition using IQ-TREE’s extended model set (MFP+MERGE). ModelFinder assigned TVM+F+R3 to both left and right flank partitions and TVME+R3 to the core partition. Given UCE structural characteristics, prior studies (Nash et al., 2022) merged left and right flank partitions into a single flank partition under the assumption that both flanks share similar evolutionary dynamics (symmetric variability gradients). To assess the influence of this merging strategy, we compared tree topologies based on the three-partition scheme (core, left flank, right flank) versus a two-partition scheme (core, merged flanks). Since IQ-TREE assigned the same model (TVM+F+R3) to both flank partitions, this model was also applied to the merged flank partition. Phylogenomic trees were generated for both partitioning schemes using IQ-TREE with 1,000 ultrafast bootstrap replicates and 1,000 approximate likelihood ratio tests (aLRT). No differences were detected in topology or branch support. To formally compare partitioning schemes, we calculated Akaike Information Criterion (AIC; Akaike, 1973), corrected AIC (AICc; Burnham and Anderson, 2002), and Bayesian Information Criterion (BIC; Schwarz, 1978); full model selection statistics, including the number of free parameters, log-likelihood, and sample size for each scheme, are provided in Table S4. Across all three criteria, the 3-partition scheme was decisively preferred over the 2-partition scheme (ΔAICc ≈ 258; Table S4) and was therefore retained for all subsequent analyses.

The initial mitochondrial partitioning scheme comprised 41 partitions: 13 protein-coding genes subdivided by codon position (first, second, third) plus 2 rRNA genes. The partitioning scheme was generated using *scaphoid.pl*, which defined gene positions as charsets and subdivided each protein-coding gene into three codon-position partitions. This partitioning scheme was implemented in IQ-TREE to determine optimal evolutionary models per partition using the extended model set (MF+MERGE) under the fast relaxed clustering search algorithm (−rclusterf; Lanfear et al., 2016). This analysis retained 15 final partitions, each with an independent substitution model (Table S5).

The UCE supermatrix (69 species, 703,208 bp, 753 loci, 3 partitions) and mitochondrial supermatrix (72 species, 12,663 bp, 15 partitions) were merged into a hybrid supermatrix (76 species, 715,871 bp, 18 partitions) using *scaphoid.pl* (−-merge-schemes), which combined datasets and partitioning schemes while maintaining partition-specific models, representing 73.8% of the 103 currently recognized extant goatfish species (Uiblein et al., 2024).

#### Phylogenetic inference

Maximum likelihood (ML) phylogenies were inferred for UCE, mitochondrial, and hybrid supermatrices using IQ-TREE with partition-specific substitution models as described above. Branch support was assessed using 100 standard bootstrap replicates (−b 100) and 1,000 approximate likelihood ratio test replicates (−alrt 1000; Anisimova and Gascuel, 2006).

To assess topology robustness toward model complexity and inference method, we performed Bayesian inference on the hybrid supermatrix considered as a single partition using PhyloBayes MPI v1.8c (Lartillot et al., 2013). We compared two site-heterogeneous models: CAT-G and CAT-GTR-G, by running four independent MCMC chains per model for 10,000 generations and saving a point every 10th generation (−x 10 1000). Separate consensus trees were generated for each model using bpcomp (PhyloBayes) with a burn-in of 200 points, sampling every 8th tree up to the last (1,000th) point (−x 200 8, hence 100 trees), and posterior probability cutoff of 0.25 (−c 0.25).

All ML and Bayesian trees were rooted on *Upeneus* following previous phylogenetic hypotheses (Longo et al., 2017; Santaquiteria et al., 2021; Nash et al., 2022), which placed the Mullidae root on the branch leading to the *Upeneus* clade based on broader Syngnathiformes phylogenies including outgroups.

#### Time calibration

We dated both the ML hybrid tree and the CAT-GTR-G Bayesian tree using a Bayesian relaxed molecular clock approach in PhyloBayes v4.1e (Lartillot et al., 2009). Two calibration points were applied: (1) the *Mullus* fossil, an articulated specimen dated to 13 Ma (Carnevale et al., 2006), and (2) a root age prior for Mullidae.

We tested four calibration strategies varying root priors and *Mullus* constraints:

1. No root constraint + *Mullus* with soft maximum (48.6 Ma) and hard minimum (13 Ma)
2. Root calibrated as interval (27.7–17 Ma) + *Mullus* with hard minimum (13 Ma), no maximum (−1)
3. Root as gamma distribution (−rp argument: mean = 21.9, sd = 2.7) + *Mullus* with soft maximum (48.6 Ma) and hard minimum (13 Ma)
4. Root as gamma distribution (−rp argument: mean = 21.9, sd = 2.7) + *Mullus* with hard minimum (13 Ma), no maximum (−1)

Root calibration values (crown age 21.9 Ma, 95% CI: 17–27.7 Ma; stem age 48.6 Ma) were derived from total-evidence dating of Syngnathiformes incorporating fossil calibrations across the order (Nash et al., 2022).

For each strategy, analyses were conducted with: (1) constant sites removed (−dc), (2) site-heterogeneous CAT model with equilibrium frequency profiles inferred non-parametrically using Dirichlet process prior (−cat), (3) GTR time-reversible matrix (−gtr), (4) discrete gamma distribution with 4 categories (−dgam 4), (5) birth-death prior on divergence times (−bd), and (6) log-normal autocorrelated relaxed clock model (−ln). Four independent MCMC chains were run per strategy, saving every 10th generation until 1,000 saved points (−x 10 1000). Divergence times were extracted using the readdiv function from PhyloBayes after a burn-in of 200 points, sampling every 8th point (−x 200 8, hence 100 data points).

Given the minor differences between the ML and Bayesian-inferred topologies and the absence of a priori criteria to favor one over the other, we based all subsequent analyses on the phylogeny derived from the ML topology. Among the four calibration strategies tested, we selected the fourth configuration, which combines a root prior distribution (mean = 21.9, sd = 2.7) with a lower-only fossil constraint for *Mullus* (hard minimum of 13 Ma without an upper maximum bound), as our definitive dating framework.

#### Lineage diversification rates

The fit of five lineage accumulation models was tested on our newly generated phylogeny, including: (1) Yule, constant speciation rate with no extinction (i.e., equivalent to a pure birth model), (2) BDcst, constant speciation and extinction rates (equivalent to a birth-death model), (3) BDvar_exp_, speciation and extinction rates both varying exponentially through time, (4) DDL, speciation rate declining linearly with increasing lineage number, and (5) DDX, speciation rate declining exponentially with increasing lineage number. The fit of each model was assessed using the bd_ML and dd_ML functions from the DDD R-package (ver. 5.2.2, Etienne et al., 2012). Following Etienne et al. (2012), likelihoods were computed from branching times only (btorph=0) and conditioned on the survival of the two daughter lineages descending from the root (cond=1). Missing species were accounted for by setting missnumspec to 27 (103 recognized species minus 76 species included in the analysis), corresponding to the currently recognized Mullidae species absent from our phylogeny. For DDL and DDX, diversity-dependence followed a linear (ddmodel=1) or exponential (ddmodel=2) decline in speciation rate with increasing species richness, respectively. Starting parameter values for each model were obtained from the maximum likelihood estimates of the next simpler nested model, with Yule seeding BDcst, and BDcst in turn seeding BDvar_exp_, DDL and DDX. Models were compared using AICc scores, applying a small sample size correction given the ratio between the number of speciation events and the number of estimated parameters.

We estimated lineage diversification rates and assessed potential shifts in regimes across the Syngnathiformes and the Mullidae using the MiSSE framework implemented in the hisse R-package (v.2.1.11) and BAMM (Bayesian Analysis of Macroevolutionary Mixtures) software (ver. 2.5.0; Rabosky, 2014), respectively. For MiSSE, we compared the fit of models with 1– 10 rate classes, setting a global sampling fraction of 25% for the Syngnathiformes and 75% for the Mullidae. Following recommended practices (Caetano et al., 2018), we model-averaged the rates inferred from models with >5% of the AICc weight, where the contribution of each model towards the mean was proportional to its Akaike weight. We plotted model-averaged rates onto the branches of the tree using the gghisse R-package (v.0.1.1; https://github.com/discindo/gghisse). For BAMM, prior distributions were estimated using the *setBAMMpriors* function from the BAMMtools R-package (ver. 2.1.11, Rabosky et al., 2014b). MCMC chains were run using the BAMM software for 10 million generations, with sampling every 1,000 generations. Then, convergence and effective sample size (>200) were assessed using the coda R-package (v0.19-4.1, Plummer et al., 2006). The first 10% of samples were discarded as burn-in. All BAMM analyses assumed the same sampling thresholds as the MiSSE analyses. A visual representation of the speciation rate through time over our phylogenomic tree was produced by the *plot.bammdata* function from the BAMMtools R-package. Speciation rate through time plot including 95% confidence interval was generated using the *plotRateThroughTime* function from the BAMMtools R-package.

### Phenotypic diversification

#### Quantification of general body morphology

Body morphology was quantified by 13 linear measurements taken to the nearest millimeter with digital caliper on each specimen (standard length - SL, head length - HL, head depth - HD, snout length - SnL, barbel length - BL, barbel width - BW, gill raker length - GRL, upper jaw length - UJL, body depth - BD, caudal peduncle length - CPL, depth - CPD and width - CPW) (Figure S6A). Each measurement represents the mean of three independent replicates. Eleven size-corrected ratios were then computed to account for allometric scaling (HL/SL, HD/HL, SnL/HL, BL/HL, BW/HL, GRL/HL, UJL/HL, BD/SL, CPL/SL, CPD/SL, CPW/SL). Morphological variation among species was summarized using a principal component analysis (PCA) performed on a matrix combining the standard length (body size) and the 11 ratios (body morphology), with data scaling enabled to account for differing units (function prcomp, R-package stats v4.4.0).

#### Quantification of head shape

Cephalic shape variation was quantified using landmark-based geometric morphometrics. To capture comprehensive head shape, the x- and y-coordinates of 17 fixed landmarks combined with 20 semi-landmarks tracing the head profile were digitized from the left side of each specimen using tpsDig (v2.31; Rohlf, 2015; Figure S6B). All configurations were superimposed via Generalized Procrustes Analysis (GPA), and the resulting Procrustes tangent coordinates were used as shape descriptors. A PCA was performed on these shape variables using the *gm.prcomp* function from the R-package geomorph (v4.0.8; Baken et al., 2021; Adams et al., 2024), retaining the first four axes (≥80% of total variance). Shape deformations along each PC axis were visualized using thin-plate spline grids (tpsRelw v1.75; Rohlf, 2015).

#### Hyoid barbel morphology and taste bud quantification

Barbel proportions were expressed as the ratio of barbel length to head length (BL/HL). For ultrastructural analyses, barbels were dehydrated in absolute ethanol, dried by critical point (Anderson, 1951), sputter-coated with gold, and imaged by scanning electron microscopy (Keyence VHX-7000). Both fresh-tissue and formalin-fixed barbels were processed following the same protocol. Up to six images were captured at x500 magnification along the whole barbel length depending on tissue quality, from which taste bud pore density (dTB, TB/mm²) and pore area (Ap, µm²; excluding the surrounding epithelial ring) were measured in ImageJ (v1.54; Abràmoff et al., 2004).

#### Quantification of pigmentation pattern

Pigmentation patterns were scored for 82 species (∼80% of extant Mullidae diversity) from illustrated taxonomic works and FishBase (fishbase.org), at the adult stage, following the trait-scoring protocol described in Frédérich et al. (2026). These data were originally compiled by L.M. and are shared with, and previously reported for a broader set of reef fish families in, Frédérich et al. (2026). Patterns were annotated independently across three body regions (head, trunk, tail) then merged. Fourteen binary traits encoding five pattern classes were scored (presence/absence of: single or multiple colors, three types of color separation, horizontal [H-sep], vertical [V-sep] and oblique [O-sep], three stripe orientations [H-, V- and O-stripes], regular blotches, saddle blotches, dots, eye stripe and dorsal fin markings). Pairwise dissimilarities among species were computed using Gower’s metric and ordinated via a Principal Coordinates Analysis (PCoA, Mouillot et al., 2014), retaining axes accounting for ≥80% of variance (32 dimensions). A reduced three-axis matrix (30% variance) was also retained to assess sensitivity to overparameterization. Traits significantly driving variation along the first three PCoA axes were identified using the *envfit* function from the R package vegan (v2.7-2; Oksanen et al. 2025).

#### Phylogenetic comparative analyses

All comparative analyses were performed on species mean values. Four phenotypic datasets were analyzed independently: body morphology, head shape, barbels and pigmentation patterns.

#### Phylomorphospace analyses

Phylogenetic relationships were projected into multivariate space constituted of regular PCA axes using phylomorphospace (phytools R-package v2.3-0, Revell, 2024), plotting pairwise combinations of the first three PCA/PCoA axes. Subclade partitioning of phylomorphospaces was tested statistically using Procrustes ANOVA with *procD.lm* from the geomorph R-package and further investigation was conducted through pairwise comparisons of genera based on the three first axes of the PCA/PCoA, using the *pairwise* function from the geomorph R-package.

#### Phylogenetic signal

The degree to which phenotypic similarity reflected phylogenetic relatedness was assessed using the multivariate extension of Blomberg’s *K* (*K*_mult_), calculated with *physignal* (geomorph; 10,000 permutations). *K*_mult_ ≈ 0 indicates phylogenetic independence, *K*_mult_ ≈ 1 is consistent with Brownian motion (BM), and *K*_mult_ > 1 indicates stronger-than-expected conservatism among relatives. Analyses used the complete raw trait matrices for traditional morphometrics and barbel characteristics, the first four PC axes for head shape, and both the 3- and 32-axis pigmentation pattern matrices.

#### Disparity-through-time analyses

The tempo of morphological diversification was examined using disparity-through-time (DTT) analyses through the *dtt* function from the R-package geiger (v2.0.11; Pennell et al., 2014), which track average subclade disparity at successive phylogenetic nodes relative to a null BM expectation (10,000 simulations; 95% confidence envelope). Disparity was measured as average squared Euclidean distance for PCA-based datasets and as average Manhattan distance for datasets combining traits into heterogeneous units. The morphological disparity index (MDI), quantifying the signed deviation observed from expected subclade disparity, was computed over the first two-third of the phylogeny’s timespan to avoid tip-sampling artefacts (Harmon et al. 2003).

#### Evolutionary independence of phenotypic datasets

Pairwise correlations among phenotypic datasets were estimated using two-block phylogenetic partial least squares with the *phylo.integration* function from the geomorph R-package (10,000 permutations) to test whether overall morphology, cephalic, barbel and pigmentation patterns variation evolved independently.

## Acknowledgments

We thank A. Dettaï and Z. Gabsi at the Muséum National d’Histoire Naturelle (MNHN), Paris, and L. Parenti and D. Pitassy at the National Museum of Natural History (NMNH), Washington, for facilitating access to their fish collections and for assistance during specimen examination; D. Pitassy also provided tissue samples on loan (see Table S3). We also thank M. Berumen and A. Kattan at King Abdullah University of Science and Technology (KAUST), R. Freitas at The Atlantic Technical University (UTA), M. Nakae at the National Museum of Nature and Science, Ibaraki, E. Post at Florida Fish and Wildlife Conservation Commission, Fish and Wildlife Research Institute (FSBC), H. L. Prestridge at Texas A&M University, S. Rothman and T. Gurevich at The Steinhardt Museum of Natural History, A. Graham at the Australian National Fish Collection, CSIRO, Hobart, M. Banse (personal collection), S.-Y. V. Liu at the National Museum of Marine Biology & Aquarium (NMMBA), Taiwan, S. Kimura at Mie University (MIE), P.R. Møller at the Zoological Museum, University of Copenhagen (ZMUC), N. Lujan and M. Zur at the Royal Ontario Museum (ROM), R. Bills and N. Mazungula at the South African Institute for Aquatic Biodiversity (SAIAB), S. South at the South Australian Museum (SAM), M. Sabbaj at the Academy of Natural Sciences of Drexel University (ANSP), Amanda Hay at the Australian Museum (AMS), Sydney, M. Sonnewald and T. Alpermann at the Senckenberg Forschungsinstitut und Naturmuseum Frankfurt (SMF), and D. Boyd at the Louisiana State University Museum of Natural Science (LSUMZ) for providing tissue samples on loan (see Table S3).

We thank the Consortium des Équipements de Calcul Intensif (CÉCI), and in particular the nic5 cluster at the University of Liège, funded by the Fonds de la Recherche Scientifique de Belgique (F.R.S.-FNRS), as well as Computerome2, made available through resources from the Department of Biology at the University of Copenhagen, for providing the computational resources used in this study.

We are grateful to G. Todinanaharay, director of the Institut Halieutique et des Sciences Marines (IH.SM), Toliara, for facilitating fieldwork in Madagascar, and to H. Mahavory, J.-L. Randrianariso, and Patrick for their assistance in the field.

## Data availability

Raw sequencing reads are available at the NCBI Sequence Read Archive (SRA) under BioProject accession [PRJNA1523961]. Pigmentation data were previously compiled by L.M. and are available at Figshare (https://doi.org/10.6084/m9.figshare.31169398; Frédérich et al., 2026). All other data supporting the findings of this study, including sequence alignments, the dated phylogeny, body shape, head shape and barbel microstructure measurements, and diversification analysis output files, are available at Zenodo (https://doi.org/10.5281/zenodo.22262806).

## Code availability

All custom scripts used to perform the analyses described in this study are available at Zenodo (https://doi.org/10.5281/zenodo.22262878), archived from the GitHub repository at [lien GitHub].

## Figure captions

Figure S1. Cophylo-plot assessing robustness and topological consistency of Mullidae phylogenomic reconstructions. (A) Comparison between nuclear (UCE) and mitochondrial DNA signals. (B) Comparison between hybrid (UCE and mitochondrial) dataset and UCE. (C) Comparison between hybrid (UCE and mitochondrial) dataset and mitochondrial. (D) Comparison of site-heterogeneous substitution models between CAT-G and CAT GTR-G models. (E) Comparison between Maximum Likelihood (ML) and Bayesian CAT-GTR-G inference methods. (F) Comparison of the current ML topology with the previous reference framework from Nash et al. (2022). In all panels, support values (bootstrap/SH-aLRT for ML or posterior probabilities for Bayesian) are only shown for nodes below 95/95 or 0.95 to highlight areas of uncertainty, and the number of unambiguous nucleotides in each concatenated sequence is indicated after the @ symbol.

Figure S2. Head shape deformation grids. Thin-plate spline deformation grids illustrating shape variation at the negative and positive extremes of (A) PC1, (B) PC2, and (C) PC3 of head shape variation in Mullidae.

Figure S3. Barbel trait relationships with head shape. Phylomorphospace projections relating the ratio of barbel length to head length (BL/HL) to (A) mean taste bud pore area (AP) and (B) mean taste bud density (dTB) across Mullidae, colored by genus.

Figure S4. Trait loadings on the pigmentation PCoA. Biplot of trait vectors associated with the first two axes of the pigmentation Principal Coordinates Analysis

Figure S5. Pigmentation phylomorphospace (PCoA2 vs PCoA3). Phylomorphospace projection of pigmentation pattern space along PCoA2 and PCoA3, with the phylogeny superimposed and tips colored by genus.

Figure S6. Morphological measurements and landmarks used in this study. (A) The 13 linear measurements taken on each specimen to quantify body morphology (standard length, head length, head depth, snout length, barbel length, barbel width, gill raker length, upper jaw length, body depth, caudal peduncle length, depth, and width; numbered 1–13), with a schematic inset illustrating the gill raker measurement. (B) The 17 fixed landmarks (red) and 20 semi-landmarks (green) digitized on the head profile for geometric morphometric analysis of head shape.

## Table captions

Table S1. Summary of morphological and genomic data sampling from all goatfish species. – indicates data not available for the species, x indicates presence data, numbers in morphology columns represent the number of individuals.

Table S2. Multivariate pairwise comparisons among genera for each dataset. Significant differences are highlighted in bold.

Table S3. Specimen and voucher information for tissue samples used in this study. For each sample, the species identifier, source institution or collection, sample ID, and associated voucher number (where available) are provided.

Table S4. Model selection statistics for the two partitioning schemes evaluated in this study. For each scheme, the number of partitions, number of free parameters (k), log-likelihood (lnL), and number of sites (n) are reported alongside the Akaike Information Criterion (AIC), corrected AIC (AICc), and Bayesian Information Criterion (BIC).

Table S5. Best-fit partitioning scheme and substitution models for the concatenated UCE and mitochondrial supermatrix, as identified by ModelFinder in IQ-TREE (Nexus format).

