## Supplementary material for "Hyoid barbels as a key innovation triggering goatfish adaptive radiation": Figure S1

Figure S1. Cophylo-plot assessing robustness and topological consistency of Mullidae phylogenomic reconstructions. (A) Comparison between nuclear (UCE) and mitochondrial DNA signal. (B) Comparison between hybrid (UCE and mitochondrial) dataset and UCE. (C) Comparison between hybrid (UCE and mitochondrial) dataset and mitochondrial. (D) Comparison of site-heterogeneous substitution models between CAT-G and CAT-GTR-G models. (E) Comparison between Maximum Likelihood (ML) and Bayesian CAT-GTR-G inference methods. (F) Comparison of the current ML topology with the previous reference framework from Nash et al. (2022). In all panels, support values (bootstrap/SH-aLRT for ML or posterior probabilities for Bayesian) are only shown for nodes below 95/95 or 0.95 to highlight areas of uncertainty, and the number of unambiguous nucleotides in each concatenated sequence is indicated after the @ symbol.

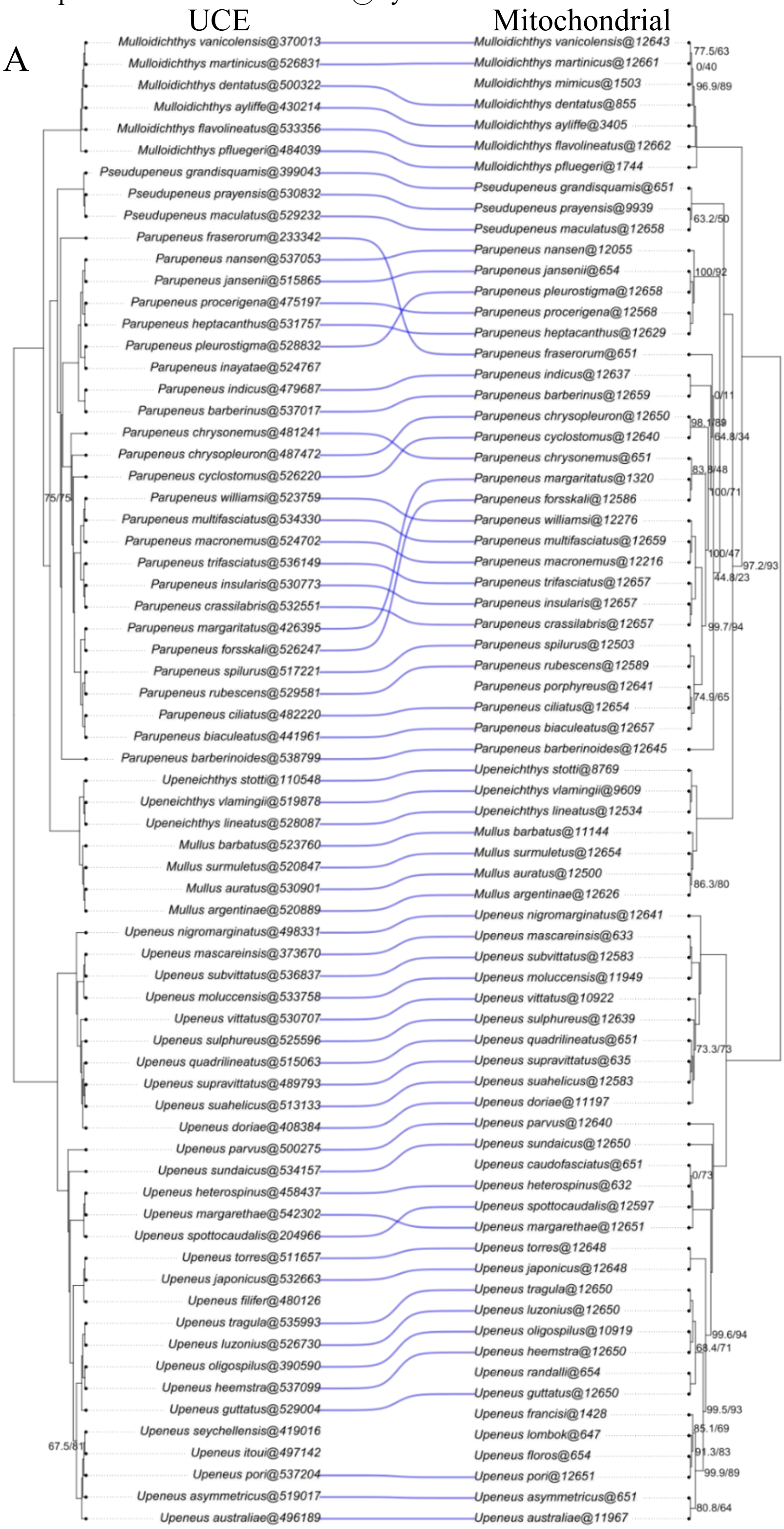

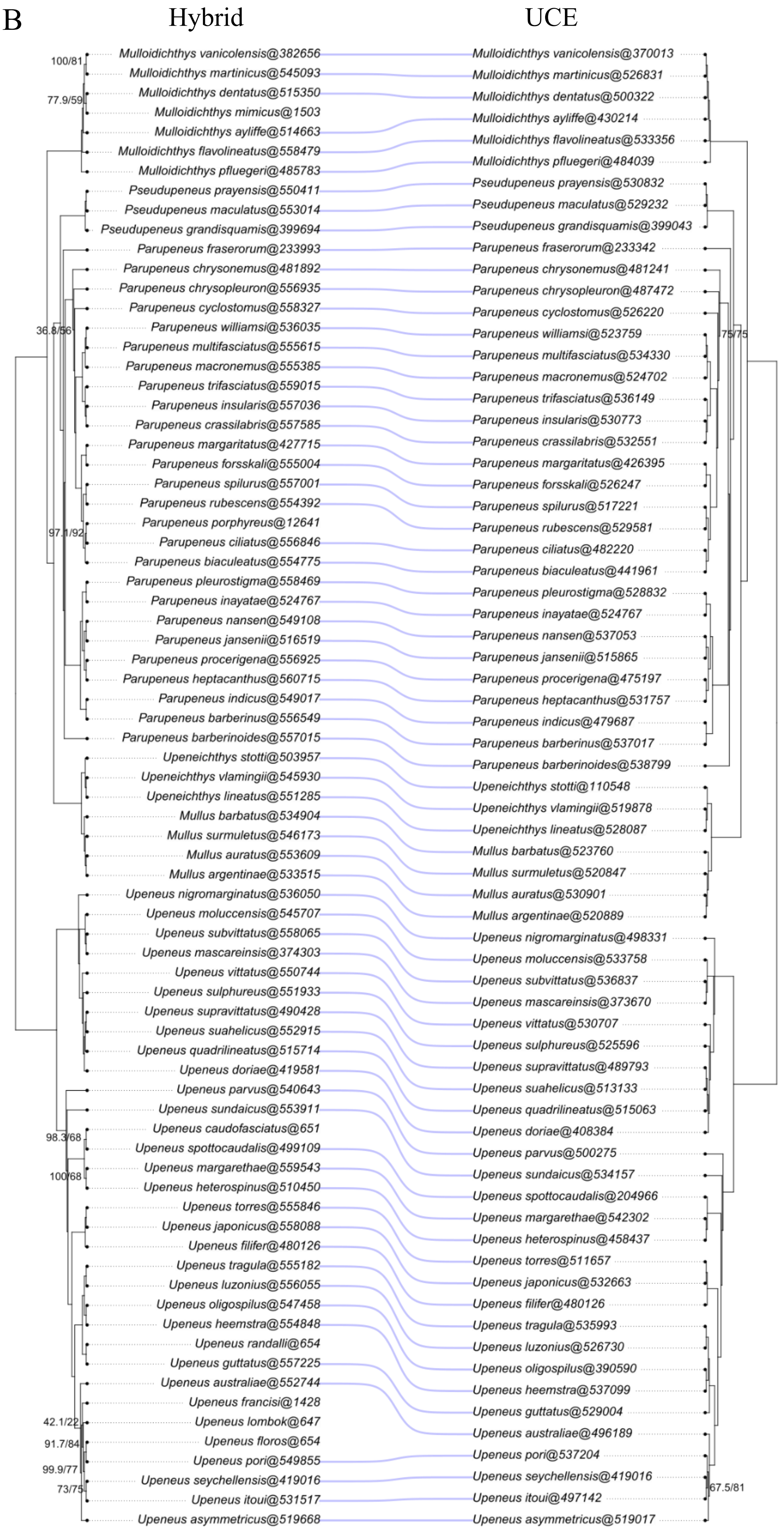
B
Hybrid
UCE
100/81
77.9/59
36.8/56
97.1/92
98.3/68
100/68
42.1/22
91.7/84
99.9/77
73/75
75/75
67.5/81
Mulloidichthys vanicolensis@382656
Mulloidichthys martinicus@545093
Mulloidichthys dentatus@515350
Mulloidichthys mimicus@1503
Mulloidichthys ayliffe@514663
Mulloidichthys flavolineatus@558479
Mulloidichthys pfluegeri@485783
Pseudupeneus prayensis@550411
Pseudupeneus maculatus@553014
Pseudupeneus grandisquamis@399694
Parupeneus fraserorum@233993
Parupeneus chrysonemus@481892
Parupeneus chrysopleuron@556935
Parupeneus cyclostomus@558327
Parupeneus williamsi@536035
Parupeneus multifasciatus@555615
Parupeneus macronemus@555385
Parupeneus trifasciatus@559015
Parupeneus insularis@557036
Parupeneus crassilabris@557585
Parupeneus margaritatus@427715
Parupeneus forsskali@555004
Parupeneus spilurus@557001
Parupeneus rubescens@554392
Parupeneus porphyreus@12641
Parupeneus ciliatus@556846
Parupeneus biaculeatus@554775
Parupeneus pleurostigma@558469
Parupeneus inayatae@524767
Parupeneus nansen@549108
Parupeneus jansenii@516519
Parupeneus procerigena@556925
Parupeneus heptacanthus@560715
Parupeneus indicus@549017
Parupeneus barberinus@556549
Parupeneus barberinoides@557015
Upeneichthys stotti@503957
Upeneichthys vlamingii@545930
Upeneichthys lineatus@551285
Mullus barbatus@534904
Mullus surmuletus@546173
Mullus auratus@553609
Mullus argentinae@533515
Upeneus nigromarginatus@536050
Upeneus moluccensis@545707
Upeneus subvittatus@558065
Upeneus mascareinsis@374303
Upeneus vittatus@550744
Upeneus sulphureus@551933
Upeneus supravittatus@490428
Upeneus suahelicus@552915
Upeneus quadrilineatus@515714
Upeneus doriae@419581
Upeneus parvus@540643
Upeneus sundaicus@553911
Upeneus caudofasciatus@651
Upeneus spottocaudalis@499109
Upeneus margarethae@559543
Upeneus heterospinus@510450
Upeneus torres@555846
Upeneus japonicus@558088
Upeneus filifer@480126
Upeneus tragula@555182
Upeneus luzonius@556055
Upeneus oligospilus@547458
Upeneus heemstra@554848
Upeneus randalli@654
Upeneus guttatus@557225
Upeneus australiae@552744
Upeneus francisi@1428
Upeneus lombok@647
Upeneus floros@654
Upeneus pori@549855
Upeneus seychellensis@419016
Upeneus itoui@531517
Upeneus asymmetricus@519668
Mulloidichthys vanicolensis@370013
Mulloidichthys martinicus@526831
Mulloidichthys dentatus@500322
Mulloidichthys ayliffe@430214
Mulloidichthys flavolineatus@533356
Mulloidichthys pfluegeri@484039
Pseudupeneus prayensis@530832
Pseudupeneus maculatus@529232
Pseudupeneus grandisquamis@399043
Parupeneus fraserorum@233342
Parupeneus chrysonemus@481241
Parupeneus chrysopleuron@487472
Parupeneus cyclostomus@526220
Parupeneus williamsi@523759
Parupeneus multifasciatus@534330
Parupeneus macronemus@524702
Parupeneus trifasciatus@536149
Parupeneus insularis@530773
Parupeneus crassilabris@532551
Parupeneus margaritatus@426395
Parupeneus forsskali@526247
Parupeneus spilurus@517221
Parupeneus rubescens@529581
Parupeneus ciliatus@482220
Parupeneus biaculeatus@441961
Parupeneus pleurostigma@528832
Parupeneus inayatae@524767
Parupeneus nansen@537053
Parupeneus jansenii@515865
Parupeneus procerigena@475197
Parupeneus heptacanthus@531757
Parupeneus indicus@479687
Parupeneus barberinus@537017
Parupeneus barberinoides@538799
Upeneichthys stotti@110548
Upeneichthys vlamingii@519878
Upeneichthys lineatus@528087
Mullus barbatus@523760
Mullus surmuletus@520847
Mullus auratus@530901
Mullus argentinae@520889
Upeneus nigromarginatus@498331
Upeneus moluccensis@533758
Upeneus subvittatus@536837
Upeneus mascareinsis@373670
Upeneus vittatus@530707
Upeneus sulphureus@525596
Upeneus supravittatus@489793
Upeneus suahelicus@513133
Upeneus quadrilineatus@515063
Upeneus doriae@408384
Upeneus parvus@500275
Upeneus sundaicus@534157
Upeneus spottocaudalis@204966
Upeneus margarethae@542302
Upeneus heterospinus@458437
Upeneus torres@511657
Upeneus japonicus@532663
Upeneus filifer@480126
Upeneus tragula@535993
Upeneus luzonius@526730
Upeneus oligospilus@390590
Upeneus heemstra@537099
Upeneus guttatus@529004
Upeneus australiae@496189
Upeneus pori@537204
Upeneus seychellensis@419016
Upeneus itoui@497142
Upeneus asymmetricus@519017

C

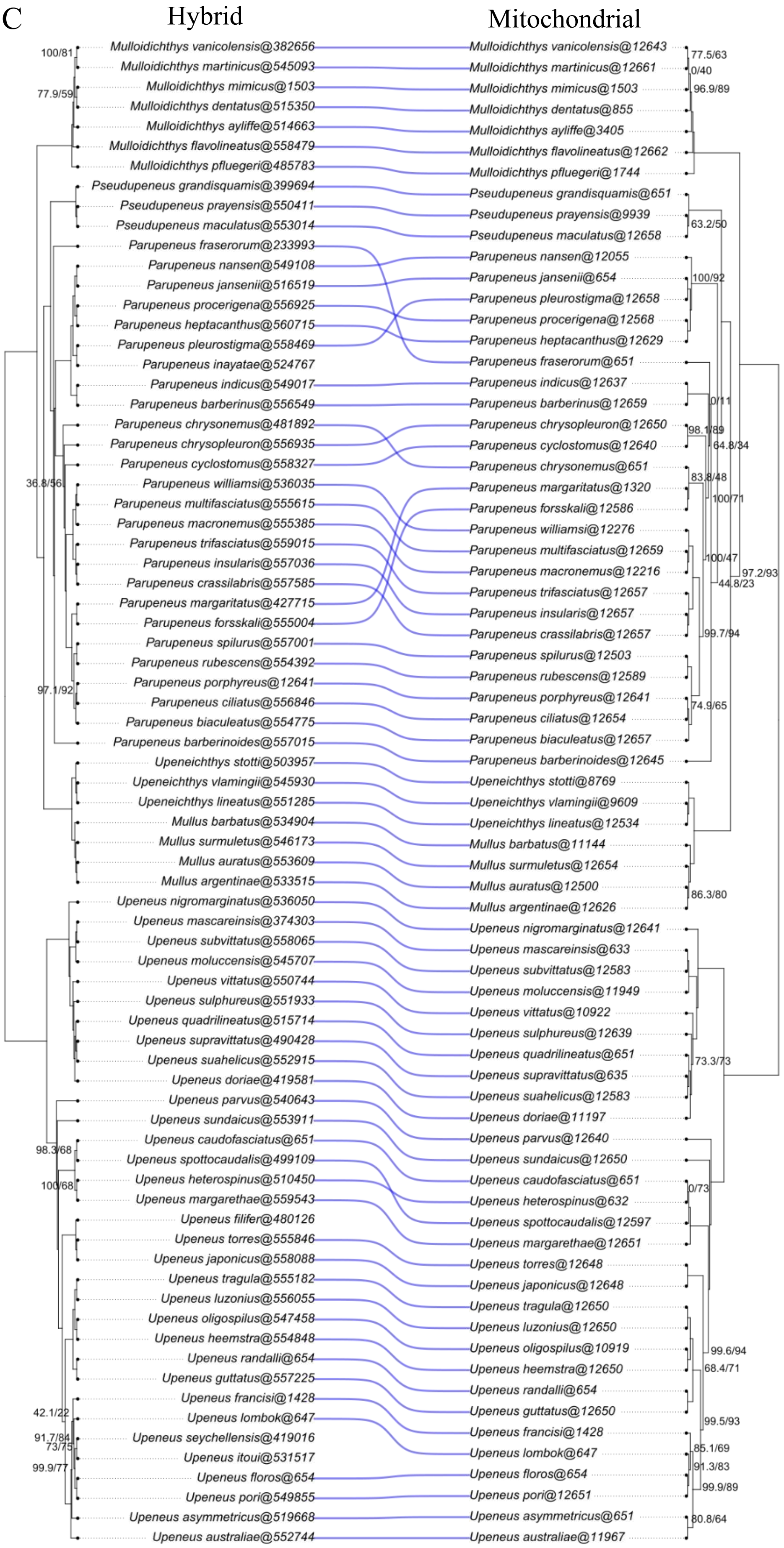
Hybrid
Mitochondrial
Mulloidichthys vanicolensis@382656
Mulloidichthys martinicus@545093
Mulloidichthys mimicus@1503
Mulloidichthys dentatus@515350
Mulloidichthys ayliffe@514663
Mulloidichthys flavolineatus@558479
Mulloidichthys pfluegeri@485783
Pseudupeneus grandisquamis@399694
Pseudupeneus prayensis@550411
Pseudupeneus maculatus@553014
Parupeneus fraserorum@233993
Parupeneus nansen@549108
Parupeneus jansenii@516519
Parupeneus procerigena@556925
Parupeneus heptacanthus@560715
Parupeneus pleurostigma@558469
Parupeneus inayatae@524767
Parupeneus indicus@549017
Parupeneus barberinus@556549
Parupeneus chrysonemus@481892
Parupeneus chrysopleuron@556935
Parupeneus cyclostomus@558327
Parupeneus williamsi@536035
Parupeneus multifasciatus@555615
Parupeneus macronemus@555385
Parupeneus trifasciatus@559015
Parupeneus insularis@557036
Parupeneus crassilabris@557585
Parupeneus margaritatus@427715
Parupeneus forsskali@555004
Parupeneus spilurus@557001
Parupeneus rubescens@554392
Parupeneus porphyreus@12641
Parupeneus ciliatus@556846
Parupeneus biaculeatus@554775
Parupeneus barberinoides@557015
Upeneichthys stotti@503957
Upeneichthys vlamingii@545930
Upeneichthys lineatus@551285
Mullus barbatus@534904
Mullus surmuletus@546173
Mullus auratus@553609
Mullus argentinae@533515
Upeneus nigromarginatus@536050
Upeneus mascareinsis@374303
Upeneus subvittatus@558065
Upeneus moluccensis@545707
Upeneus vittatus@550744
Upeneus sulphureus@551933
Upeneus quadrilineatus@515714
Upeneus supravittatus@490428
Upeneus suahelicus@552915
Upeneus doriae@419581
Upeneus parvus@540643
Upeneus sundaicus@553911
Upeneus caudofasciatus@651
Upeneus spottocaudalis@499109
Upeneus heterospinus@510450
Upeneus margarethae@559543
Upeneus filifer@480126
Upeneus torres@555846
Upeneus japonicus@558088
Upeneus tragula@555182
Upeneus luzonius@556055
Upeneus oligospilus@547458
Upeneus heemstra@554848
Upeneus randalli@654
Upeneus guttatus@557225
Upeneus francisi@1428
Upeneus lombok@647
Upeneus seychellensis@419016
Upeneus itoui@531517
Upeneus floros@654
Upeneus pori@549855
Upeneus asymmetricus@519668
Upeneus australiae@552744
Mulloidichthys vanicolensis@12643
Mulloidichthys martinicus@12661
Mulloidichthys mimicus@1503
Mulloidichthys dentatus@855
Mulloidichthys ayliffe@3405
Mulloidichthys flavolineatus@12662
Mulloidichthys pfluegeri@1744
Pseudupeneus grandisquamis@651
Pseudupeneus prayensis@9939
Pseudupeneus maculatus@12658
Parupeneus nansen@12055
Parupeneus jansenii@654
Parupeneus pleurostigma@12658
Parupeneus procerigena@12568
Parupeneus heptacanthus@12629
Parupeneus fraserorum@651
Parupeneus indicus@12637
Parupeneus barberinus@12659
Parupeneus chrysopleuron@12650
Parupeneus cyclostomus@12640
Parupeneus chrysonemus@651
Parupeneus margaritatus@1320
Parupeneus forsskali@12586
Parupeneus williamsi@12276
Parupeneus multifasciatus@12659
Parupeneus macronemus@12216
Parupeneus trifasciatus@12657
Parupeneus insularis@12657
Parupeneus crassilabris@12657
Parupeneus spilurus@12503
Parupeneus rubescens@12589
Parupeneus porphyreus@12641
Parupeneus ciliatus@12654
Parupeneus biaculeatus@12657
Parupeneus barberinoides@12645
Upeneichthys stotti@8769
Upeneichthys vlamingii@9609
Upeneichthys lineatus@12534
Mullus barbatus@11144
Mullus surmuletus@12654
Mullus auratus@12500
Mullus argentinae@12626
Upeneus nigromarginatus@12641
Upeneus mascareinsis@633
Upeneus subvittatus@12583
Upeneus moluccensis@11949
Upeneus vittatus@10922
Upeneus sulphureus@12639
Upeneus quadrilineatus@651
Upeneus supravittatus@635
Upeneus suahelicus@12583
Upeneus doriae@11197
Upeneus parvus@12640
Upeneus sundaicus@12650
Upeneus caudofasciatus@651
Upeneus heterospinus@632
Upeneus spottocaudalis@12597
Upeneus margarethae@12651
Upeneus torres@12648
Upeneus japonicus@12648
Upeneus tragula@12650
Upeneus luzonius@12650
Upeneus oligospilus@10919
Upeneus heemstra@12650
Upeneus randalli@654
Upeneus guttatus@12650
Upeneus francisi@1428
Upeneus lombok@647
Upeneus floros@654
Upeneus pori@12651
Upeneus asymmetricus@651
Upeneus australiae@11967

D

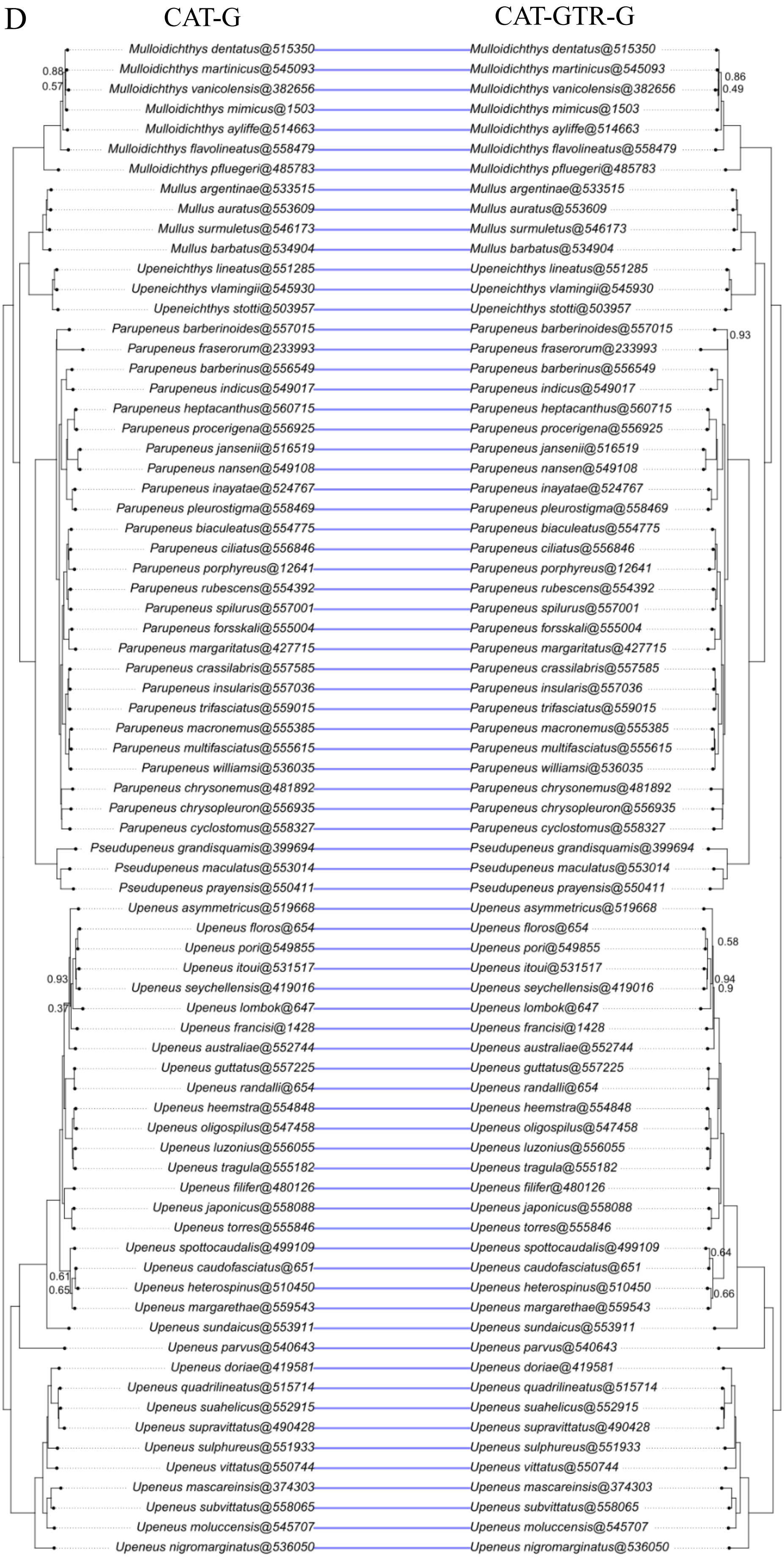

CAT-G
CAT-GTR-G
Mulloidichthys dentatus@515350
Mulloidichthys martinicus@545093
Mulloidichthys vanicolensis@382656
Mulloidichthys mimicus@1503
Mulloidichthys ayliffe@514663
Mulloidichthys flavolineatus@558479
Mulloidichthys pfluegeri@485783
Mullus argentinae@533515
Mullus auratus@553609
Mullus surmuletus@546173
Mullus barbatus@534904
Upeneichthys lineatus@551285
Upeneichthys vlamingii@545930
Upeneichthys stotti@503957
Parupeneus barberinoides@557015
Parupeneus fraserorum@233993
Parupeneus barberinus@556549
Parupeneus indicus@549017
Parupeneus heptacanthus@560715
Parupeneus procerigena@556925
Parupeneus jansenii@516519
Parupeneus nansen@549108
Parupeneus inayatae@524767
Parupeneus pleurostigma@558469
Parupeneus biaculeatus@554775
Parupeneus ciliatus@556846
Parupeneus porphyreus@12641
Parupeneus rubescens@554392
Parupeneus spilurus@557001
Parupeneus forsskali@555004
Parupeneus margaritatus@427715
Parupeneus crassilabris@557585
Parupeneus insularis@557036
Parupeneus trifasciatus@559015
Parupeneus macronemus@555385
Parupeneus multifasciatus@555615
Parupeneus williamsi@536035
Parupeneus chrysonemus@481892
Parupeneus chrysopleuron@556935
Parupeneus cyclostomus@558327
Pseudupeneus grandisquamis@399694
Pseudupeneus maculatus@553014
Pseudupeneus prayensis@550411
Upeneus asymmetricus@519668
Upeneus floros@654
Upeneus pori@549855
Upeneus itoui@531517
Upeneus seychellensis@419016
Upeneus lombok@647
Upeneus francisi@1428
Upeneus australiae@552744
Upeneus guttatus@557225
Upeneus randalli@654
Upeneus heemstra@554848
Upeneus oligospilus@547458
Upeneus luzonius@556055
Upeneus tragula@555182
Upeneus filifer@480126
Upeneus japonicus@558088
Upeneus torres@555846
Upeneus spottocaudalis@499109
Upeneus caudofasciatus@651
Upeneus heterospinus@510450
Upeneus margarethae@559543
Upeneus sundaicus@553911
Upeneus parvus@540643
Upeneus doriae@419581
Upeneus quadrilineatus@515714
Upeneus suahelicus@552915
Upeneus supravittatus@490428
Upeneus sulphureus@551933
Upeneus vittatus@550744
Upeneus mascareinsis@374303
Upeneus subvittatus@558065
Upeneus moluccensis@545707
Upeneus nigromarginatus@536050
0.88
0.57
0.86
0.49
0.93
0.93
0.37
0.58
0.94
0.9
0.61
0.65
0.64
0.66

E

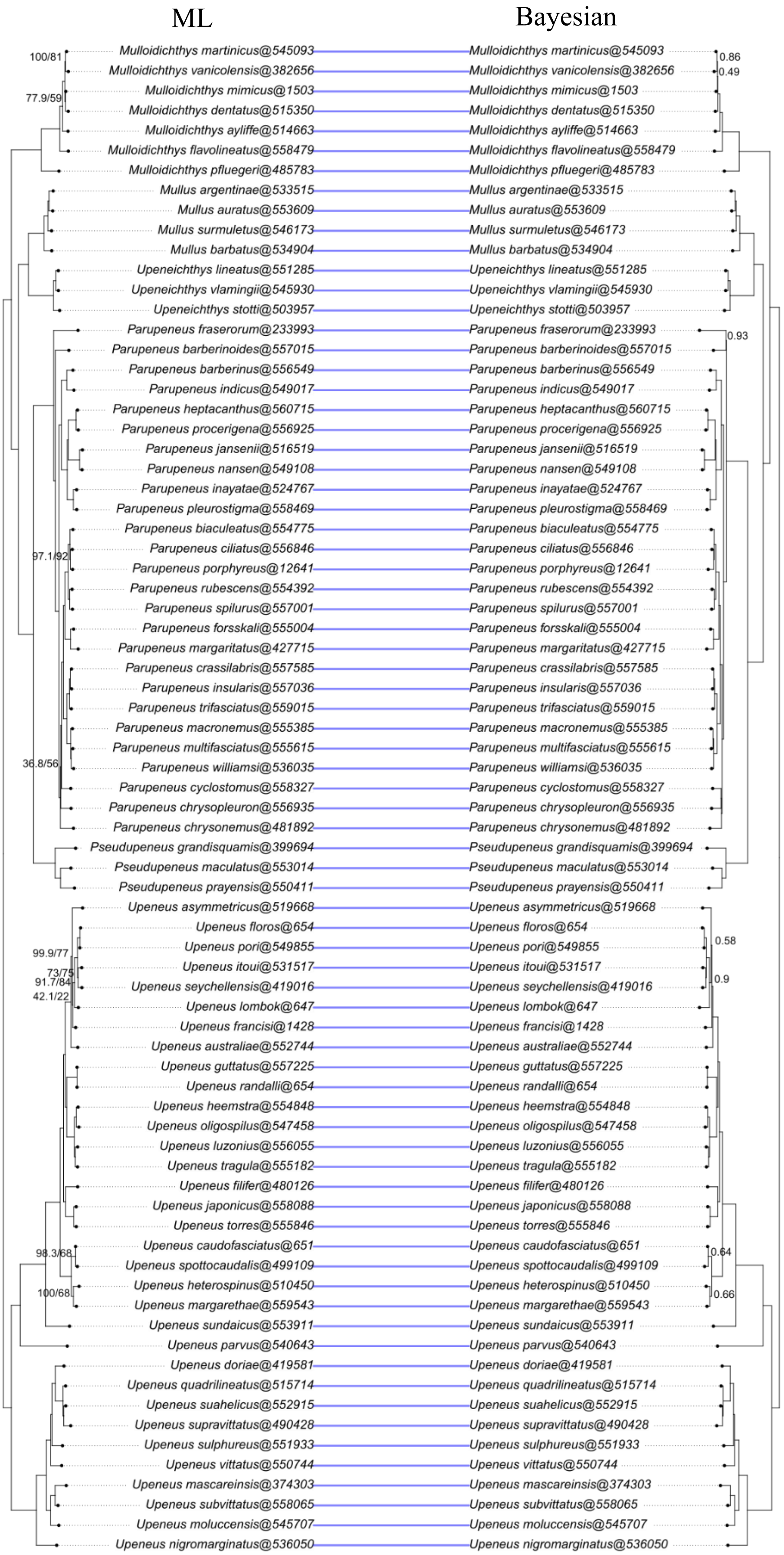
ML
Bayesian
Mulloidichthys martinicus@545093
Mulloidichthys vanicolensis@382656
Mulloidichthys mimicus@1503
Mulloidichthys dentatus@515350
Mulloidichthys ayliffe@514663
Mulloidichthys flavolineatus@558479
Mulloidichthys pfluegeri@485783
Mullus argentinae@533515
Mullus auratus@553609
Mullus surmuletus@546173
Mullus barbatus@534904
Upeneichthys lineatus@551285
Upeneichthys vlamingii@545930
Upeneichthys stotti@503957
Parupeneus fraserorum@233993
Parupeneus barberinoides@557015
Parupeneus barberinus@556549
Parupeneus indicus@549017
Parupeneus heptacanthus@560715
Parupeneus procerigena@556925
Parupeneus jansenii@516519
Parupeneus nansen@549108
Parupeneus inayatae@524767
Parupeneus pleurostigma@558469
Parupeneus biaculeatus@554775
Parupeneus ciliatus@556846
Parupeneus porphyreus@12641
Parupeneus rubescens@554392
Parupeneus spilurus@557001
Parupeneus forsskali@555004
Parupeneus margaritatus@427715
Parupeneus crassilabris@557585
Parupeneus insularis@557036
Parupeneus trifasciatus@559015
Parupeneus macronemus@555385
Parupeneus multifasciatus@555615
Parupeneus williamsi@536035
Parupeneus cyclostomus@558327
Parupeneus chrysopleuron@556935
Parupeneus chrysonemus@481892
Pseudupeneus grandisquamis@399694
Pseudupeneus maculatus@553014
Pseudupeneus prayensis@550411
Upeneus asymmetricus@519668
Upeneus floros@654
Upeneus pori@549855
Upeneus itoui@531517
Upeneus seychellensis@419016
Upeneus lombok@647
Upeneus francisi@1428
Upeneus australiae@552744
Upeneus guttatus@557225
Upeneus randalli@654
Upeneus heemstra@554848
Upeneus oligospilus@547458
Upeneus luzonius@556055
Upeneus tragula@555182
Upeneus filifer@480126
Upeneus japonicus@558088
Upeneus torres@555846
Upeneus caudofasciatus@651
Upeneus spottocaudalis@499109
Upeneus heterospinus@510450
Upeneus margarethae@559543
Upeneus sundaicus@553911
Upeneus parvus@540643
Upeneus doriae@419581
Upeneus quadrilineatus@515714
Upeneus suahelicus@552915
Upeneus supravittatus@490428
Upeneus sulphureus@551933
Upeneus vittatus@550744
Upeneus mascareinsis@374303
Upeneus subvittatus@558065
Upeneus moluccensis@545707
Upeneus nigromarginatus@536050
100/81
77.9/59
97.1/92
36.8/56
99.9/77
73/75
91.7/84
42.1/22
98.3/68
100/68
0.86
0.49
0.93
0.58
0.9
0.64
0.66

F

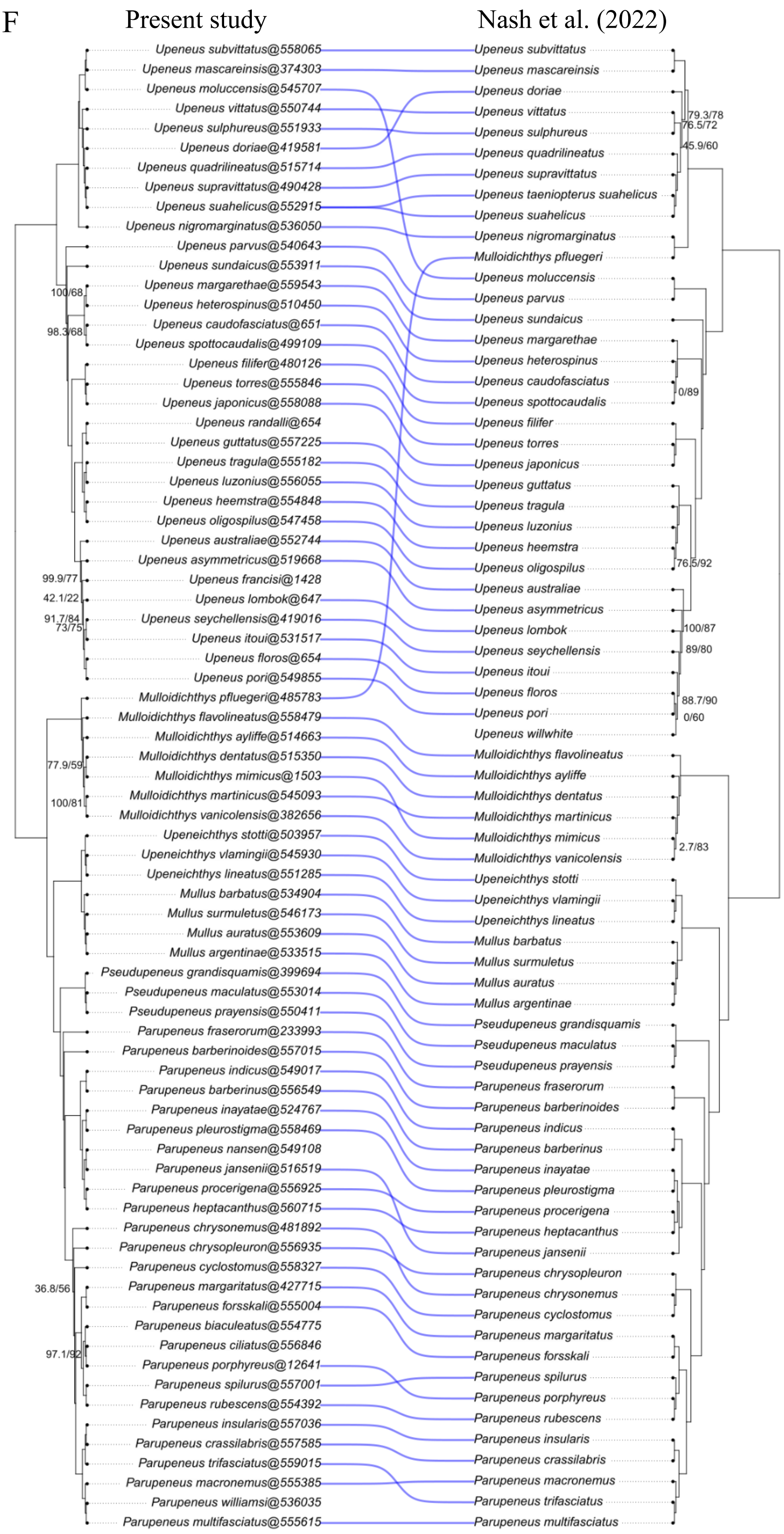
Present study
Nash et al. (2022)
Upeneus subvittatus@558065
Upeneus mascareinsis@374303
Upeneus moluccensis@545707
Upeneus vittatus@550744
Upeneus sulphureus@551933
Upeneus doriae@419581
Upeneus quadrilineatus@515714
Upeneus supravittatus@490428
Upeneus suahelicus@552915
Upeneus nigromarginatus@536050
Upeneus parvus@540643
Upeneus sundaicus@553911
Upeneus margarethae@559543
Upeneus heterospinus@510450
Upeneus caudofasciatus@651
Upeneus spottocaudalis@499109
Upeneus filifer@480126
Upeneus torres@555846
Upeneus japonicus@558088
Upeneus randalli@654
Upeneus guttatus@557225
Upeneus tragula@555182
Upeneus luzonius@556055
Upeneus heemstra@554848
Upeneus oligospilus@547458
Upeneus australiae@552744
Upeneus asymmetricus@519668
Upeneus francisi@1428
Upeneus lombok@647
Upeneus seychellensis@419016
Upeneus itoui@531517
Upeneus floros@654
Upeneus pori@549855
Mulloidichthys pfluegeri@485783
Mulloidichthys flavolineatus@558479
Mulloidichthys ayliffe@514663
Mulloidichthys dentatus@515350
Mulloidichthys mimicus@1503
Mulloidichthys martinicus@545093
Mulloidichthys vanicolensis@382656
Upeneichthys stotti@503957
Upeneichthys vlamingii@545930
Upeneichthys lineatus@551285
Mullus barbatus@534904
Mullus surmuletus@546173
Mullus auratus@553609
Mullus argentinae@533515
Pseudupeneus grandisquamis@399694
Pseudupeneus maculatus@553014
Pseudupeneus prayensis@550411
Parupeneus fraserorum@233993
Parupeneus barberinoides@557015
Parupeneus indicus@549017
Parupeneus barberinus@556549
Parupeneus inayatae@524767
Parupeneus pleurostigma@558469
Parupeneus nansen@549108
Parupeneus jansenii@516519
Parupeneus procerigena@556925
Parupeneus heptacanthus@560715
Parupeneus chrysonemus@481892
Parupeneus chrysopleuron@556935
Parupeneus cyclostomus@558327
Parupeneus margaritatus@427715
Parupeneus forsskali@555004
Parupeneus biaculeatus@554775
Parupeneus ciliatus@556846
Parupeneus porphyreus@12641
Parupeneus spilurus@557001
Parupeneus rubescens@554392
Parupeneus insularis@557036
Parupeneus crassilabris@557585
Parupeneus trifasciatus@559015
Parupeneus macronemus@555385
Parupeneus williamsi@536035
Parupeneus multifasciatus@555615
100/68
98.3/68
99.9/77
42.1/22
91.7/84
73/75
77.9/59
100/81
36.8/56
97.1/92
Upeneus subvittatus
Upeneus mascareinsis
Upeneus doriae
Upeneus vittatus
Upeneus sulphureus
Upeneus quadrilineatus
Upeneus supravittatus
Upeneus taeniopterus suahelicus
Upeneus suahelicus
Upeneus nigromarginatus
Mulloidichthys pfluegeri
Upeneus moluccensis
Upeneus parvus
Upeneus sundaicus
Upeneus margarethae
Upeneus heterospinus
Upeneus caudofasciatus
Upeneus spottocaudalis
Upeneus filifer
Upeneus torres
Upeneus japonicus
Upeneus guttatus
Upeneus tragula
Upeneus luzonius
Upeneus heemstra
Upeneus oligospilus
Upeneus australiae
Upeneus asymmetricus
Upeneus lombok
Upeneus seychellensis
Upeneus itoui
Upeneus floros
Upeneus pori
Upeneus willwhite
Mulloidichthys flavolineatus
Mulloidichthys ayliffe
Mulloidichthys dentatus
Mulloidichthys martinicus
Mulloidichthys mimicus
Mulloidichthys vanicolensis
Upeneichthys stotti
Upeneichthys vlamingii
Upeneichthys lineatus
Mullus barbatus
Mullus surmuletus
Mullus auratus
Mullus argentinae
Pseudupeneus grandisquamis
Pseudupeneus maculatus
Pseudupeneus prayensis
Parupeneus fraserorum
Parupeneus barberinoides
Parupeneus indicus
Parupeneus barberinus
Parupeneus inayatae
Parupeneus pleurostigma
Parupeneus procerigena
Parupeneus heptacanthus
Parupeneus jansenii
Parupeneus chrysopleuron
Parupeneus chrysonemus
Parupeneus cyclostomus
Parupeneus margaritatus
Parupeneus forsskali
Parupeneus spilurus
Parupeneus porphyreus
Parupeneus rubescens
Parupeneus insularis
Parupeneus crassilabris
Parupeneus macronemus
Parupeneus trifasciatus
Parupeneus multifasciatus
79.3/78
76.5/72
45.9/60
0/89
76.5/92
100/87
89/80
88.7/90
0/60
2.7/83
