## Supplementary figures and images for "Hyoid barbels as a key innovation triggering goatfish adaptive radiation"

### Figure S2

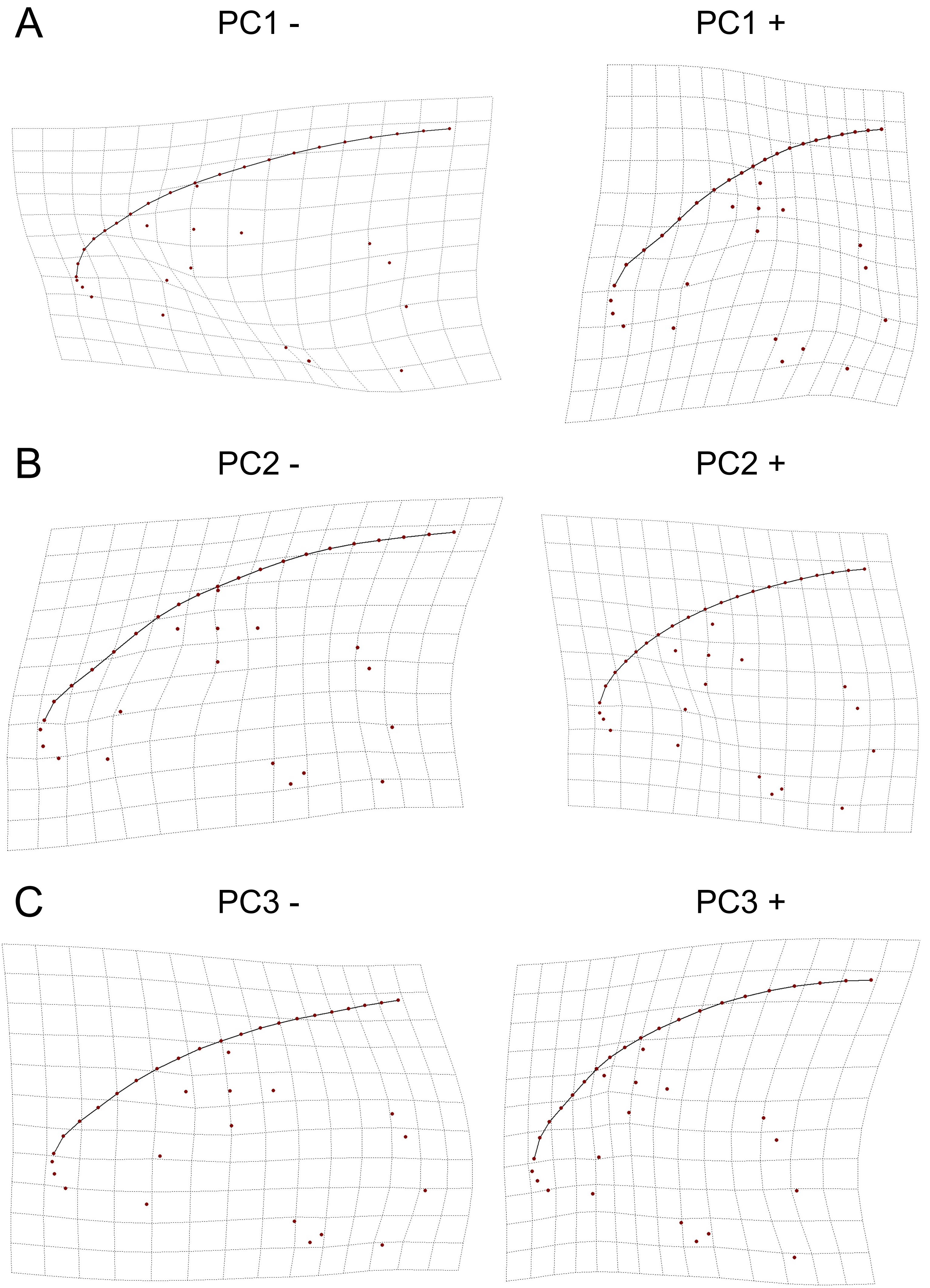

### Figure S3

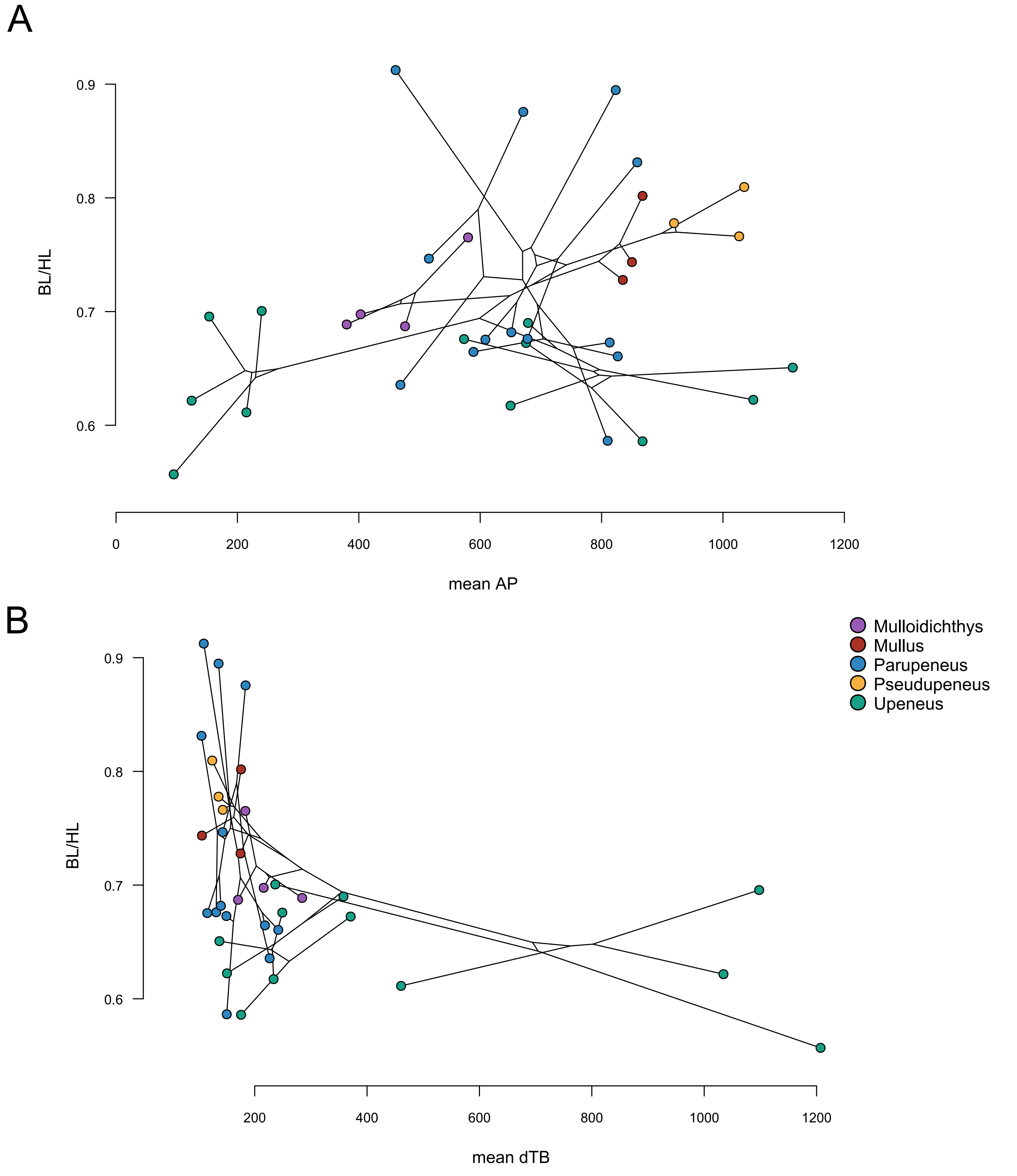

### Figure S4

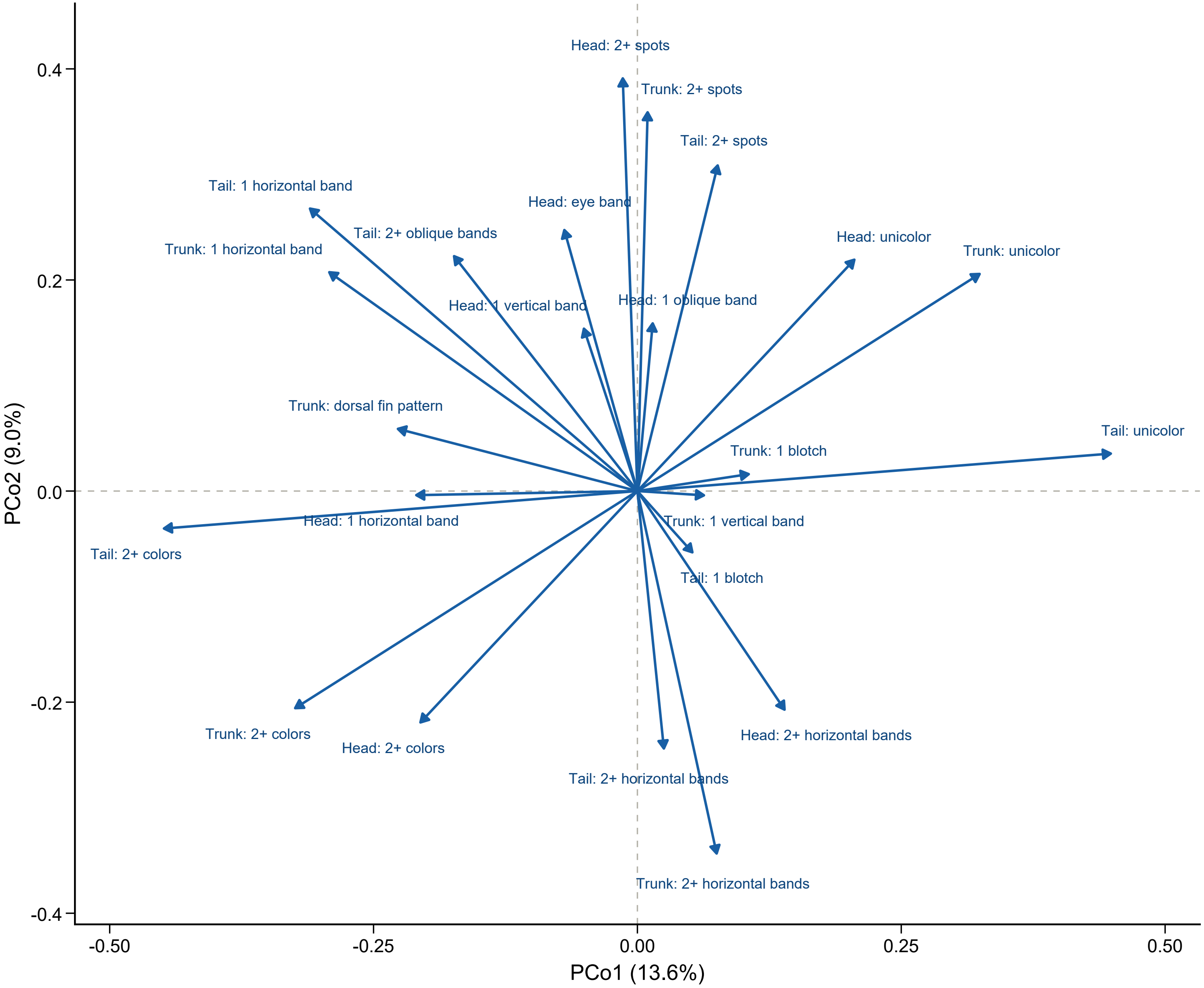

### Figure S5

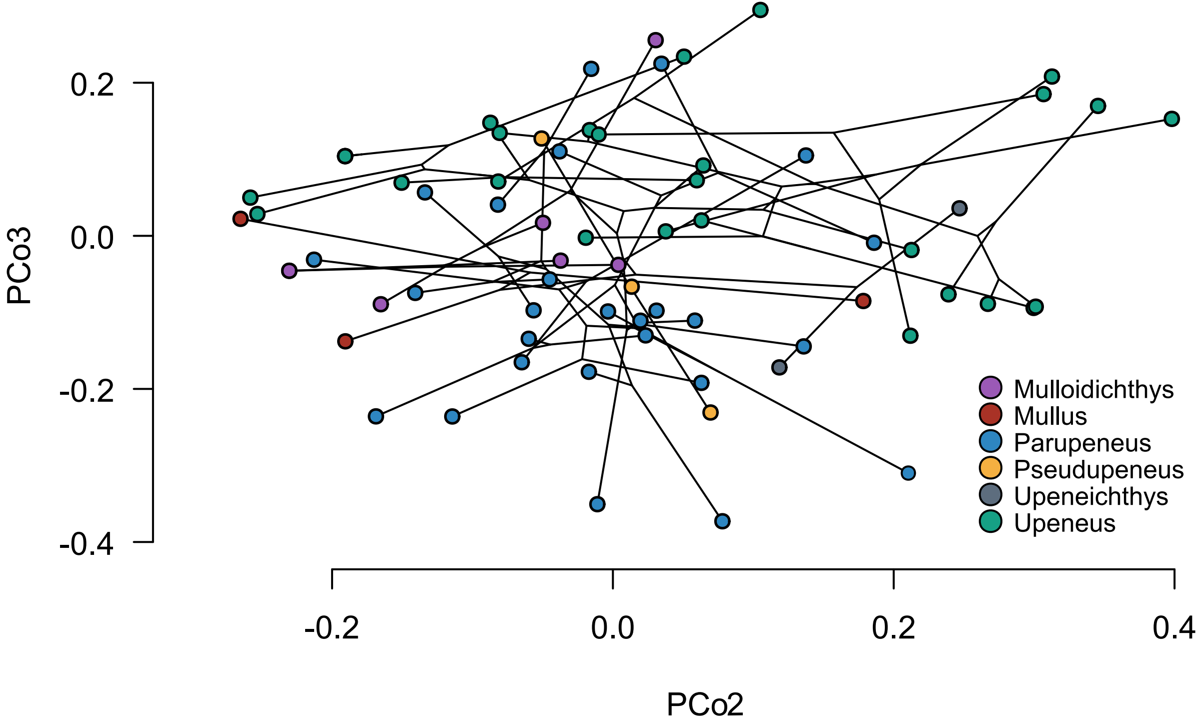

### Figure S6

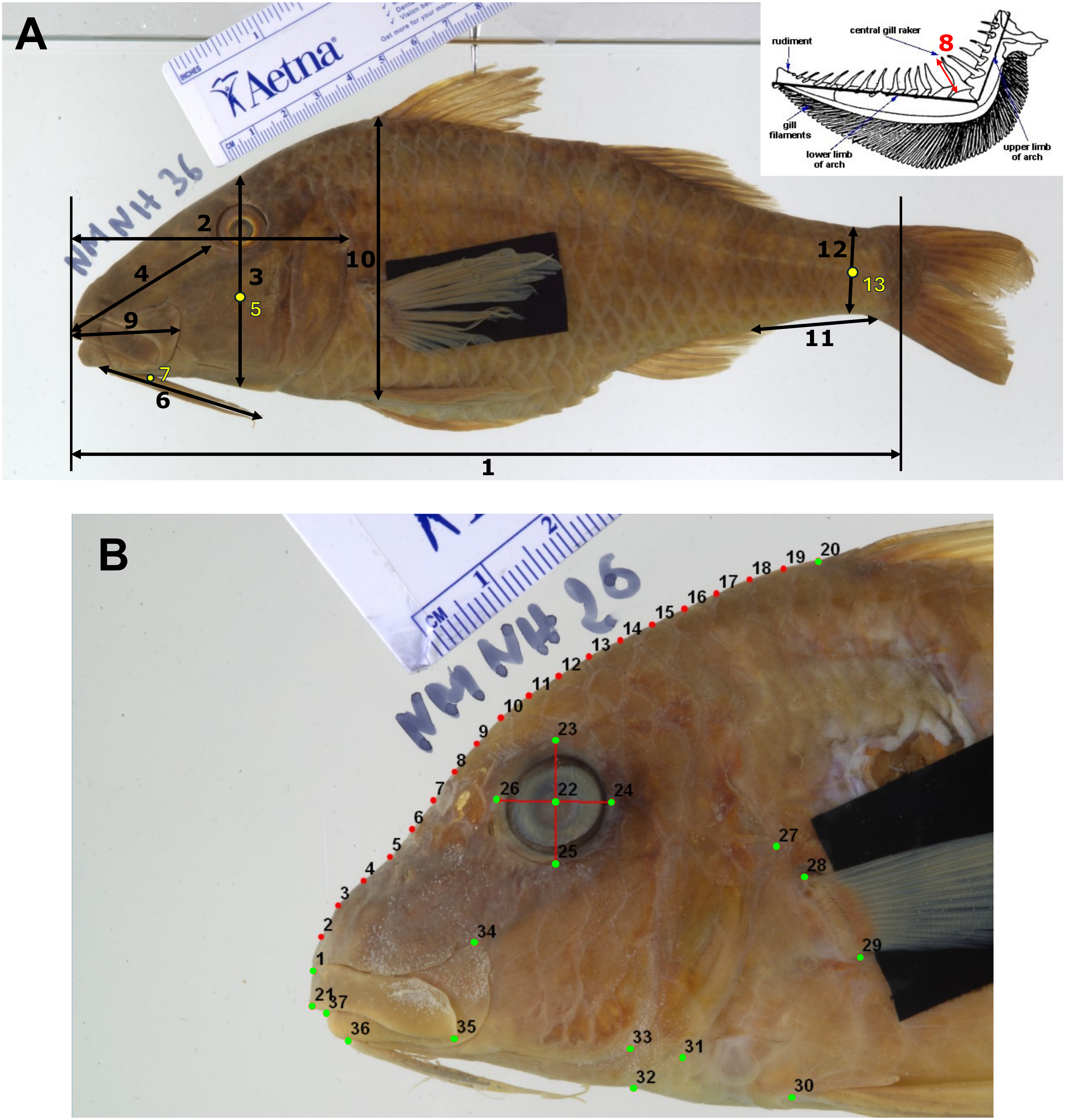
