## Supplementary material for "Hyoid barbels as a key innovation triggering goatfish adaptive radiation": Table S1

Table S1. Summary of morphological and genomic data sampling from all goatfish species. – indicates data not available for the species, x indicates presence data, numbers in morphology columns represent the number of individuals.

|  | Morphology | | | | Phylogeny | |
| --- | --- | --- | --- | --- | --- | --- |
| Species | Body morphology | Head shape | Barbels | Pigmentation patterns | UCEs | Mitogenomes |
| *Parupeneus angulatus* | – | – | – | – | – | – |
| *Parupeneus barberinoides* | 5 | 5 | – | x | x | x |
| *Parupeneus barberinus* | 15 | 15 | 11 | x | x | x |
| *Parupeneus biaculeatus* | 1 | 1 | – | x | x | x |
| *Parupeneus chrysonemus* | 4 | 4 | 3 | x | x | x |
| *Parupeneus chrysopleuron* | 3 | 3 | – | x | x | x |
| *Parupeneus ciliatus* | 3 | 3 | 2 | x | x | x |
| *Parupeneus crassilabris* | 3 | 3 | 4 | x | x | x |
| *Parupeneus cyclostomus* | 15 | 9 | 4 | x | x | x |
| *Parupeneus diagonalis* | 2 | 2 | – | x | – | – |
| *Parupeneus forsskali* | 5 | 9 | 3 | x | x | x |
| *Parupeneus fraserorum* | – | – | – | x | x | x |
| *Parupeneus heptacanthus* | 30 | 26 | 4 | x | x | x |
| *Parupeneus inayatae* | – | – | – | x | x | – |
| *Parupeneus indicus* | 12 | 10 | 11 | x | x | x |
| *Parupeneus insularis* | 4 | 3 | – | x | x | x |
| *Parupeneus jansenii* | 4 | 3 | – | x | x | x |
| *Parupeneus louise* | – | – | – | x | – | – |
| *Parupeneus macronemus* | 15 | 9 | 6 | x | x | x |
| *Parupeneus margaritatus* | 5 | 4 | 1 | x | x | x |
| *Parupeneus minys* | 2 | 2 | – | x | – | – |
| *Parupeneus moffitti* | 1 | – | – | x | – | – |
| *Parupeneus multifasciatus* | 3 | 3 | 2 | x | x | x |
| *Parupeneus nansen* | 1 | – | – | x | x | x |
| *Parupeneus orientalis* | 1 | 1 | – | x | – | – |
| *Parupeneus pleurostigma* | 14 | 14 | 6 | x | x | x |
| *Parupeneus porphyreus* | 5 | 4 | – | x | – | x |
| *Parupeneus posteli* | 6 | 3 | – | x | – | – |
| *Parupeneus procerigena* | 4 | 4 | – | x | x | x |
| *Parupeneus rubescens* | 10 | 10 | 1 | x | x | x |
| *Parupeneus seychellensis* | 1 | 1 | – | x | – | – |
| *Parupeneus sinai* | 1 | 1 | – | x | – | – |
| *Parupeneus spilurus* | 4 | 4 | – | x | x | x |
| *Parupeneus trifasciatus* | 16 | 12 | – | x | x | x |
| *Parupeneus williamsi* | 3 | 2 | – | x | x | x |
| *Upeneus andamanensis* | – | – | – | – | – | – |
| *Upeneus asymmetricus* | 6 | 6 | 3 | x | x | x |
| *Upeneus aurorae* | – | – | – | – | – | – |
| *Upeneus australiae* | – | – | – | x | x | x |
| *Upeneus brevignathus* | – | – | – | – | – | – |
| *Upeneus caudofasciatus* | – | – | – | x | – | x |
| *Upeneus davidaromi* | 4 | 1 | – | x | – | – |
| *Upeneus dimipavlov* | 1 | 1 | – | – | – | – |
| *Upeneus doriae* | – | – | – | x | x | x |
| *Upeneus elongatus* | – | – | – | – | – | – |
| *Upeneus farnis* | – | – | – | x | – | – |
| *Upeneus filifer* | – | – | – | – | x | – |
| *Upeneus floros* | – | – | – | x | – | x |
| *Upeneus francisi* | 2 | 1 | – | x | – | x |
| *Upeneus gubal* | 1 | 1 | – | – | – | – |
| *Upeneus guttatus* | 5 | 5 | 2 | x | x | x |
| *Upeneus heemstra* | – | – | – | x | x | x |
| *Upeneus heterospinus* | 1 | 1 | – | x | x | x |
| *Upeneus huan* | – | – | – | – | – | – |
| *Upeneus indicus* | – | – | – | x | – | – |
| *Upeneus itoui* | 2 | 2 | – | x | x | – |
| *Upeneus japonicus* | 3 | 3 | 3 | x | x | x |
| *Upeneus lombok* | – | – | – | – | – | x |
| *Upeneus luzonius* | 3 | 3 | 3 | x | x | x |
| *Upeneus madras* | – | – | – | – | – | – |
| *Upeneus margarethae* | 6 | 4 | 2 | x | x | x |
| *Upeneus mascareinsis* | 6 | 4 | 1 | – | x | x |
| *Upeneus moluccensis* | 36 | 13 | 4 | x | x | x |
| *Upeneus mouthami* | 3 | 2 | – | – | – | – |
| *Upeneus niebuhri* | 1 | – | – | – | – | – |
| *Upeneus nigromarginatus* | 2 | 2 | – | x | x | x |
| *Upeneus oligospilus* | 3 | 3 | 2 | x | x | x |
| *Upeneus parvus* | 6 | 6 | – | x | x | x |
| *Upeneus pori* | 24 | 1 | – | x | x | x |
| *Upeneus quadrilineatus* | – | – | – | x | x | x |
| *Upeneus randalli* | – | – | – | x | – | x |
| *Upeneus saiab* | – | – | – | – | – | – |
| *Upeneus seychellensis* | – | – | – | – | x | x |
| *Upeneus spottocaudalis* | – | – | – | – | x | x |
| *Upeneus stenopsis* | 2 | 1 | – | – | – | – |
| *Upeneus suahelicus* | 8 | 9 | 6 | x | x | x |
| *Upeneus subvittatus* | 3 | 3 |  | x | x | x |
| *Upeneus sulphureus* | 16 | 16 | 9 | x | x | x |
| *Upeneus sundaicus* | 6 | 6 | 2 | x | x | x |
| *Upeneus supravittatus* | – | – | – | x | x | x |
| *Upeneus taeniopterus* | 4 | 4 | 2 | x | – | – |
| *Upeneus torres* | – | – | – | x | x | x |
| *Upeneus tragula* | 3 | 3 | – | x | x | x |
| *Upeneus vanuatu* | 3 | 2 | – | – | – | – |
| *Upeneus vittatus* | 61 | 28 | 9 | x | x | x |
| *Upeneus willwhite* | – | – | – | x | – | – |
| *Upeneichthys lineatus* | 3 | 4 | – | x | x | x |
| *Upeneichthys stotti* | – | – | – | – | x | x |
| *Upeneichthys vlamingii* | – | – | – | x | x | x |
| *Mullus auratus* | 3 | 3 | 2 | x | x | x |
| *Mullus argentinae* | 2 | – | – | – | x | x |
| *Mullus barbatus* | 24 | 3 | 3 | x | x | x |
| *Mullus surmuletus* | 8 | 5 | 2 | x | x | x |
| *Mulloidichthys ayliffe* | – | – | – | x | x | x |
| *Mulloidichthys dentatus* | 3 | 3 | 2 | x | x | x |
| *Mulloidichthys flavolineatus* | 20 | 19 | 16 | x | x | x |
| *Mulloidichthys martinicus* | 4 | 4 | 1 | x | x | x |
| *Mulloidichthys mimicus* | 3 | 3 | – | x | – | x |
| *Mulloidichthys pfluegeri* | 4 | 3 | – | x | x | x |
| *Mulloidichthys vanicolensis* | 21 | 21 | 11 | x | x | x |
| *Pseudupeneus grandisquamis* | 5 | 5 | 2 | x | x | x |
| *Pseudupeneus maculatus* | 4 | 4 | 5 | x | x | x |
| *Pseudupeneus prayensis* | 3 | 3 | 3 | x | x | x |
| TOTAL SPECIMENS | 526 | 381 | 153 |  |  |  |
| TOTAL SPECIES | 73 | 69 | 36 | 82 | 69 | 72 |
