## Supplementary material for "Hyoid barbels as a key innovation triggering goatfish adaptive radiation": Table S2

Table S2. Multivariate pairwise comparisons among genera for each dataset. Significant differences are highlighted in bold.

|  | *Mullus* | *Parupeneus* | *Pseudupeneus* | *Upeneichthys* | *Upeneus* |
| --- | --- | --- | --- | --- | --- |
| *Mulloidichthys* | 0.0542 | **0.0001** | **0.0025** | **0.0009** | **0.0001** |
| *Mullus* |  | **0.0001** | **0.0002** | **0.0001** | **0.0001** |
| *Parupeneus* |  |  | 0.2472 | 0.1412 | **0.0001** |
| *Pseudupeneus* |  |  |  | 0.2351 | **0.0001** |
| *Upeneichthys* |  |  |  |  | **0.0001** |

A. Body morphology

B. Head shape

|  | *Mullus* | *Parupeneus* | *Pseudupeneus* | *Upeneichthys* | *Upeneus* |
| --- | --- | --- | --- | --- | --- |
| *Mulloidichthys* | **0.002** | **0.0001** | **0.0001** | **0.0001** | **0.0001** |
| *Mullus* |  | **0.0088** | **0.0460** | **0.0175** | **0.0001** |
| *Parupeneus* |  |  | 0.8116 | **0.0286** | **0.0001** |
| *Pseudupeneus* |  |  |  | **0.1260** | **0.0001** |
| *Upeneichthys* |  |  |  |  | **0.0001** |

C. Barbels

|  | *Mullus* | *Parupeneus* | *Pseudupeneus* | *Upeneus* |
| --- | --- | --- | --- | --- |
| *Mulloidichthys* | 0.1365 | 0.2910 | **0.0347** | 0.2522 |
| *Mullus* |  | 0.4756 | 0.6633 | 0.0794 |
| *Parupeneus* |  |  | 0.1517 | **0.0275** |
| *Pseudupeneus* |  |  |  | **0.0190** |

D. Pigmentation patterns

|  | *Mullus* | *Parupeneus* | *Pseudupeneus* | *Upeneichthys* | *Upeneus* |
| --- | --- | --- | --- | --- | --- |
| *Mulloidichthys* | 0.2262 | 0.5312 | 0.4373 | 0.1812 | **0.0095** |
| *Mullus* |  | 0.1376 | **0.0440** | 0.0830 | 0.2110 |
| *Parupeneus* |  |  | 0.4833 | 0.3588 | **0.0001** |
| *Pseudupeneus* |  |  |  | 0.6347 | **0.0099** |
| *Upeneichthys* |  |  |  |  | 0.0967 |
