## Supplementary material for "Hyoid barbels as a key innovation triggering goatfish adaptive radiation": Table S3

| Species ID | Source | ID | Voucher |
|---|---|---|---|
| Mulloidichthys_ayliffe_1_arbor2 | KAUST | RS5951 | |
| Mulloidichthys_ayliffe_2_arbor2 | KAUST | RS5952 | |
| Mulloidichthys_ayliffe_nash | CAS | CAS-ICH237544 | ICH237544 |
| Mulloidichthys_dentatus_longo | SIO | SIO 07-88 | |
| Mulloidichthys_dentatus_nash | CAS | CAS-ICH236117 | ICH236117 |
| Mulloidichthys_flavolineatus_1_stille | USNM | 442395 | 442395 |
| Mulloidichthys_flavolineatus_2_stille | Tel Aviv | P.15669 (RH1633) | |
| Mulloidichthys_flavolineatus_santa | KU | 5644 | KUI 32516 |
| Mulloidichthys_martinicus_longo | PC Wainwright | PCW 3433 | |
| Mulloidichthys_martinicus_stiller | Rui Freitas | | |
| Mulloidichthys_pfluegeri_arbor | NSMT | NSMT-P 131247 | |
| Mulloidichthys_vanicolensis_longo | SJ Longo | SJL046 | |
| Mullus_argentinae_santa | UNMDP | T-0134 | UNMDP 0134 |
| Mullus_auratus_1_stiller | FSBC | 34025 | FSBC:34025 |
| Mullus_auratus_2_stiller | Texas A&M | 17070.01; 17070.06 | |
| Mullus_auratus_longo | KU | KU 1229 | |
| Mullus_barbatus_stiller | Tel Aviv | P.16661 | |
| Mullus_surmuletus_1_stiller | Existing genome | GCA_901007815_1 | |
| Mullus_surmuletus_2_stiller | Existing genome | GCA_903171835_1 | |
| Parupeneus_barberinoides_1_stiller | SAM | ABTC82688 | |
| Parupeneus_barberinoides_2_stiller | CSIRO | GT 9652 | H 6341-22 |
| Parupeneus_barberinoides_longo | PC Wainwright | PCW 1677 | |
| Parupeneus_barberinus_longo | SJ Longo | SJL072 | |
| Parupeneus_barberinus_santa | KU | 4124 | KUI 31836 |
| Parupeneus_barberinus_stiller | Guam/Marine Banse | L4 | |
| Parupeneus_biaculeatus_1_stiller | NSYU | HO-3647 | |
| Parupeneus_biaculeatus_2_stiller | MIE | FRLM56080 | |
| Parupeneus_bifasciatus_longo | PC Wainwright | PCW 2157 | |
| Parupeneus_chrysonemus_nash | BPBM | BPBM-PCMB567 | PCMB567 |
| Parupeneus_chrysopleuron_1_arbor | MIE | FRLM 37506 | |
| Parupeneus_chrysopleuron_1_stiller | USNM | 423612 | USNM:423612 |
| Parupeneus_chrysopleuron_2_arbor | MIE | FRLM 37515 | |
| Parupeneus_chrysopleuron_2_stiller | CSIRO | GT 1916 | H 6381-05 |
| Parupeneus_chrysopleuron_nash | KUNHM | KUNHM-10560 | KUI 4146 |
| Parupeneus_ciliatus_arbor | NSMT | NSMT-P 106172 | |
| Parupeneus_ciliatus_stiller | USNM | 409348 | USNM:409348 |
| Parupeneus_crassilabris_santa | USNM | AG9RE44 | PI-0264 |
| Parupeneus_crassilabris_stiller | USNM | 437884 | USNM:437884 |
| Parupeneus_cyclostomus_1_stiller | Guam/Marine Banse | L5 | |
| Parupeneus_cyclostomus_2_stiller | Tel Aviv | P.15690 (RH1685) | |
| Parupeneus_cyclostomus_3_stiller | Madagascar/L. M. | TUL57Bis | |
| Parupeneus_cyclostomus_longo | SJ Longo | SJL074 | |
| Parupeneus_forsskali_santa | FMNH | RS1304T54 | FMNH 143490 |
| Parupeneus_forsskali_stiller | Tel Aviv | P. 15661 (RH1617) | |
| Parupeneus_fraserorum_nash | SAIAB | SAIAB-AV/2010-026 | 97405 |
| Parupeneus_heptacanthus_1_arbor2 | KAUST | RS7519 | |
| Parupeneus_heptacanthus_1_stiller | Tel Aviv | P.15737 (RH1806) | |
| Parupeneus_heptacanthus_2_arbor2 | KAUST | RS7520 | |
| Parupeneus_heptacanthus_2_stiller | Tel Aviv | (RH2028) | |
| Parupeneus_heptacanthus_nash | FMNH | FMNH-125506 | 125506 |
| Parupeneus_inayatae_santa | CSIRO | IN00502 | CSIRO unreg LM81 |

| | | | |
|---|---|---|---|
| Parupeneus_indicus_santa | USNM | AG9RD89 | PI-0209 |
| Parupeneus_indicus_stiller | USNM | 403245 | USNM:403245 |
| Parupeneus_insularis_santa | USNM | AG5NQ87 | mbio1648 |
| Parupeneus_insularis_stiller | USNM | 424183 | USNM:424183 |
| Parupeneus_jansenii_santa | CSIRO | GT 4602 | CSIRO H 6926-06 |
| Parupeneus_macronemus_nash | CAS | CAS-ICH236336 | ICH236336 |
| Parupeneus_macronemus_stiller | ZMUC | P2397934 | ZMUC:P2397934 |
| Parupeneus_margaritatus_nash | LSUMZ | LSUMZ | 18000 |
| Parupeneus_multifasciatus_longo | SJ Longo | SJL003 | |
| Parupeneus_multifasciatus_stiller | ROM | ROMI-T04016M | ROMI-T04016M |
| Parupeneus_nansen_stiller | SAIAB | TZW-256 | |
| Parupeneus_pleurostigma_santa | KU | 743 | USNM 334645 |
| Parupeneus_pleurostigma_stiller | USNM | 403237 | USNM:403237 |
| Parupeneus_procerigena_1_stiller | SAIAB | ACEP08-1835 | |
| Parupeneus_procerigena_2_stiller | KU | T6817 | |
| Parupeneus_procerigena_santa | KU | 6817 | SAIAB 77857 |
| Parupeneus_rubescens_longo | SIO | SIO 04-51 | |
| Parupeneus_rubescens_stiller | KAUST | RS4172 | |
| Parupeneus_spilurus_Sally_arbor2 | SAM | ABTC82678 | |
| Parupeneus_spilurus_Seishi_arbor2 | MIE | FRLM 56115 | |
| Parupeneus_spilurus_nash | CAS | CAS-PAP02 | 237673 |
| Parupeneus_spilurus_stiller | CSIRO | 355 Par spi 01 | H 4101-04 |
| Parupeneus_trifasciatus_1_arbor2 | KAUST | RS7118 | |
| Parupeneus_trifasciatus_2_arbor2 | KAUST | RS7119 | |
| Parupeneus_trifasciatus_stiller | Madagascar/L. M. | TUL-136 | |
| Parupeneus_williamsi_stiller | USNM | 409089 | USNM:409089 |
| Pseudupeneus_grandisquamis_longo | SIO | SIO 07-193 | |
| Pseudupeneus_maculatus_longo | PC Wainwright | PCW 3687 | |
| Pseudupeneus_maculatus_stiller | ANSP | ANSP199413 | |
| Pseudupeneus_prayensis_santa | USNM | AD9NF17 | CV11-150 |
| Pseudupeneus_prayensis_stiller | USNM | 405151 | USNM:405151 |
| Upeneichthys_lineatus_longo | AM | I.45633-036 | |
| Upeneichthys_lineatus_nash | AMS | AMS-I.45027-021 | I.45027-021 |
| Upeneichthys_lineatus_stiller | AMS | I.42890-023 | AMS:I.42890-023 |
| Upeneichthys_stotti_stiller | CSIRO | GT 5950 | H 6351-07 |
| Upeneichthys_vlamingii_stiller | CSIRO | GT 266 | H 6346-01 |
| Upeneus_asymmetricus_santa | CSIRO | IN01738 | CSIRO H 7417-02 |
| Upeneus_australiae_santa | CSIRO | GT 4135 | CSIRO H 6898-09 |
| Upeneus_australiae_stiller | CSIRO | GT 9595 | H 6927-03 |
| Upeneus_doriae_stiller | KAUST | RS7502 | |
| Upeneus_filifer_santa | CSIRO | CMR003702A | CSIRO unreg |
| Upeneus_guttatus_1_santa | CSIRO | GT 4122 | CSIRO H 6519-18 |
| Upeneus_guttatus_2_santa | CSIRO | IN02796 | CSIRO H 7217-07 |
| Upeneus_guttatus_stiller | SMF | KAU17-104 | not catalogued |
| Upeneus_heemstra_arbor | LSUMZ | LSUMZ 20950 | |
| Upeneus_heemstra_stiller | SMF | KAU12-0626 | SMF:35056 |
| Upeneus_heterospinus_1_arbor | NSYU | NMMB-P28522 | |
| Upeneus_heterospinus_2_arbor | NSYU | NMMB-P28523 | |
| Upeneus_itoui_arbor2 | MIE | FRLM 40465 | |
| Upeneus_itoui_nash | KAUM | KAUM-I.112722 | KAUM-I.112722 |
| Upeneus_japonicus_santa | KU | 10324 | NSMT-P 105304 |
| Upeneus_japonicus_stiller | KU | T10473 | |

| | | | |
|---|---|---|---|
| Upeneus_luzonius_nash | CSIRO | CSIRO-GT4129 | H6790-04 |
| Upeneus_luzonius_stiller | CSIRO | GT 4129 | H 6790-04 |
| Upeneus_margarethae_1_stiller | USNM | 437634 | USNM:437634 |
| Upeneus_margarethae_2_stiller | SMF | KAU12-0584 | SMF:35060 |
| Upeneus_margarethae_nash | CSIRO | CSIRO-GT10581 | H8224-21 |
| Upeneus_mascareinsis_nash | SAIAB | SAIAB-HM07-261 | 82328 |
| Upeneus_moluccensis_santa | KU | 6747 | SAIAB 76409 |
| Upeneus_moluccensis_stiller | Tel Aviv | P.16804 (RH1895) | |
| Upeneus_nigromarginatus_nash | Bos personal coll | BOS | BOS |
| Upeneus_nigromarginatus_stiller | USNM | 403319 | USNM:403319 |
| Upeneus_oligospilus_nash | KAUST | KAUST-RS7497 | RS7497 |
| Upeneus_oligospilus_stiller | KAUST | RS7498 | |
| Upeneus_parvus_1_arbor2 | Texas A&M | 17068.03 | |
| Upeneus_parvus_2_arbor2 | Texas A&M | 17070.15 | |
| Upeneus_parvus_nash | KUNHM | KUNHM-3926 | KUI 29671 |
| Upeneus_pfluegeri_santa | USNM | AG9RE99 | |
| Upeneus_pori_stiller | Tel Aviv | (RH1943) | |
| Upeneus_quadrilineatus_santa | CSIRO | IN02060 | MZB 22936 |
| Upeneus_seychellensis_nash | SAIAB | SAIAB-ACEP08-1603 | 84280 |
| Upeneus_spottocaudalis_nash | CSIRO | CSIRO-GT9537 | H6799-02 |
| Upeneus_spottocaudalis_stiller | CSIRO | GT 9535 | H 7462-02 |
| Upeneus_suahelicus_1_nash | SAIAB | SAIAB-MZ/2010-024 | 199990 |
| Upeneus_suahelicus_1_stiller | SMF | KAU11-003 | SMF:33643 |
| Upeneus_suahelicus_2_nash | SAIAB | SAIAB-HM07-371 | 82006 |
| Upeneus_suahelicus_2_stiller | SAIAB | 44 | |
| Upeneus_suahelicus_3_stiller | SAIAB | HM07-372 | |
| Upeneus_subvittatus_arbor2 | LSUMZ | LSUMZ 13 822 | |
| Upeneus_subvittatus_nash | | | |
| Upeneus_subvittatus_stiller | USNM | 435634 | USNM:435634 |
| Upeneus_sulphureus_santa | USNM | AG9RC75 | PI-0093 |
| Upeneus_sulphureus_stiller | Madagascar/L. M. | TUL159 | |
| Upeneus_sundaicus_santa | CSIRO | GT 4266 | CSIRO unreg |
| Upeneus_sundaicus_stiller | USNM | 445541 | USNM:445541 |
| Upeneus_supravittatus_nash | SAIAB | SAIAB-PAK14-013 | 200573 |
| Upeneus_torres_santa | CSIRO | GT 1917 | CSIRO H 6405-05 |
| Upeneus_torres_stiller | CSIRO | GT 1920 | H 6452-04 |
| Upeneus_tragula_1_stiller | NSMT | NSMT-P76727 | |
| Upeneus_tragula_2_stiller | CSIRO | GT 4074 | H 6895-08 |
| Upeneus_tragula_longo | SJ Longo | SJL028 | |
| Upeneus_vittatus_arbor | NSMT | NSMT-P 77815 | |
| Upeneus_vittatus_stiller | ZMUC | 10048 | |
