## Supplementary material for "Hyoid barbels as a key innovation triggering goatfish adaptive radiation": Table S4

| Partition scheme | Number of partitions | Free parameters (k) | lnL | Number of sites (n) | AIC | AICc | BIC |
| --- | --- | --- | --- | --- | --- | --- | --- |
| 2 partitions | 2 | 20 | −2,845,913.290 | 648,994 | 5,691,866.58 | 5,691,866.58 | 5,692,094.24 |
| 3 partitions | 3 | 30 | −2,845,774.406 | 648,994 | 5,691,608.81 | 5,691,608.81 | 5,691,950.31 |

Table S4. Model selection statistics for the two partitioning schemes evaluated in this study. For each scheme, the number of partitions, number of free parameters (k), log-likelihood (lnL), and number of sites (n) are reported alongside the Akaike Information Criterion (AIC), corrected AIC (AICc), and Bayesian Information Criterion (BIC).
