## Supplementary material for "Hyoid barbels as a key innovation triggering goatfish adaptive radiation": Table S5

Table S5. Best-fit partitioning scheme and substitution models for the concatenated UCE and mitochondrial supermatrix, as identified by ModelFinder in IQ-TREE (Nexus format).

#nexus

begin sets;

charset ATP6_1_ATP8_1_ND2_1 = 1-687\3 688-855\3 4588-5634\3;

charset ATP6_2_ATP8_2_ND2_2_ND5_2 = 2-687\3 689-855\3 4589-5634\3 7670-9507\3;

charset ATP6_3_ATP8_3_ND2_3 = 3-687\3 690-855\3 4590-5634\3;

charset COX1_1_COX3_1_CYTB_1 = 856-1509\3 2209-2994\3 2995-3612\3;

charset COX1_2 = 857-1509\3;

charset COX1_3_COX2_3 = 858-1509\3 1512-2208\3;

charset COX2_1_ND4L_1 = 1510-2208\3 7372-7668\3;

charset COX2_2_COX3_2_CYTB_2_ND1_2_ND3_2_ND4_2_ND4L_2 = 1511-2208\3 2210-2994\3 2996-3612\3 3614-4587\3 5636-5985\3 5987-7371\3 7373-7668\3;

charset COX3_3_CYTB_3_ND4L_3 = 2211-2994\3 2997-3612\3 7374-7668\3;

charset ND1_1_ND3_1_ND4_1 = 3613-4587\3 5635-5985\3 5986-7371\3;

charset ND1_3_ND3_3_ND4_3_ND5_3 = 3615-4587\3 5637-5985\3 5988-7371\3 7671-9507\3;

charset ND5_1_rrnL_rrnS = 7669-9507\3 10030-11713 11714-12663;

charset ND6_1 = 9508-10029\3;

charset ND6_2 = 9509-10029\3;

charset ND6_3 = 9510-10029\3;

charset uce_left = 12664-12903 13546-13753 14216-14529 15142-15372 15805-15873 16311-16553 17238-17287 17793-17886 18739-18797 19516-20001 20271-20464 21226-21467 22132-22181 22858-23162 24070-24119 24881-24936 25706-25875 27255-28452 28825-29134 29804-30028 30485-30696 31214-31313 31918-32164 32856-33764 34060-34162 35132-35211 35927-36509 36908-37164 37871-37986 38878-39069 39520-39786 40373-40953 41122-41193 41975-42358 43103-43302 43909-44090 44940-45053 45893-46068 46680-47305 47684-48307 48702-48872 49635-49834 50610-50920 51265-52227 52642-53317 53505-53614 54513-54798 55494-55987 56677-56944 57567-58289 58443-58997 59454-59782 60064-60622 60777-61080 61771-61862 62647-62856 63456-64072 64413-65177 65377-65895 66231-66800 67049-67264 67823-68142 68703-69403 69769-70354 70627-71483 71621-71766 72227-72521 73098-73155 73494-73779 74208-74347 74911-75348 75644-75985 76612-76810 77566-77767 78346-78695 79640-80156 80394-80446 81183-81294 82031-82162 83249-83834 84273-84889 85258-85394 86299-86718 87173-87529 87966-88056 88665-88847 89060-89275 90016-90321 91037-91277 91871-92114 92806-93519 93777-93918 95316-95384 96109-96294 97174-97263 98193-98346 99230-99593 100051-100177 100631-100722 101647-102134 102446-102579 103163-103696 104053-104660 104935-105530 105914-105990 106731-107421 107660-108275 108451-109051 109502-109634 110398-111015 111260-111906 112275-113040 113324-113519 114391-114736 115422-116278 116422-117097 117353-117670 118213-118506 118939-119468 119803-119953 120685-121023 121791-122555 123080-124195 124494-124584 125580-126032 126157-126826 127563-127846 128473-128995 129431-129616 130326-131574 131790-132069 132697-132848 133705-134075 134714-134891 135623-135837 136465-136607 137395-137824 138384-139110 139270-139605 140266-140674 141601-141662 142127-142735 143037-143095 143962-144214 144572-144631 145283-145352 146350-146519 147456-147740 148415-148871 149433-149844 150322-150926 151086-151260 151903-152021 152506-152597 153216-153992 154261-154668 155044-155201 156254-156312 157043-157150 158027-158291 158925-159354 160158-160752 161107-161183 161984-162908 163128-163465 164306-164516 165419-165468 166103-166344 167078-167311 167967-168336 168825-169436 169709-170355 170581-170659 171536-172459 172726-172936 173708-174464 174703-174789 175332-175508 176488-176607 177198-177690 177920-178079 178763-178931 179436-179679 180236-180708 181017-181072 181640-181748 182551-183167 183438-183760 184469-185056 185599-185819 186378-186926 187411-188086 188380-188475 189247-190192 190349-190821 191360-192052 192435-192502 193010-193059 193597-194262 194738-195177 195496-195583 196527-196577 197206-198037 198222-198330 198684-199117 199577-199806 200466-201273 201557-201719 202599-203231 203692-204173 204455-204583 205478-206401 206696-206900 207435-207484 208091-209138 209288-210013 210307-210707 211389-211861 212247-212962 213176-213793 214220-214340 214968-215256 215745-215895 217425-217700 218434-218547 219255-219331 220117-220230 220796-220846 221604-221720 223290-223640 224634-225139 225448-226145 226434-226532 227199-227378 228039-228232 228890-229236 229582-229876 230534-231044 231396-231672 232218-232370 233615-233926 234143-234736 235103-235439 236083-236691 237073-237226 237777-238015 238728-239503 239705-239879 240575-240929 241596-242073 242491-243222 243425-243657 244548-245315 245559-245791 246393-246566 247090-247231 247839-248711 249016-249324 250441-250606 251305-251654 252302-252454 253236-253978 254219-254330 255058-255320 256013-256407 256991-257742 257940-258251 258839-259206 259751-260177 260361-260825 261351-261407 261823-262010 262666-263143 263881-264301 264675-265006 266273-266501 266921-267491 267816-268014 268812-269627 269812-270598 270890-271314 271641-271858 272381-272915 273067-273175 273472-273527 274389-274640 275134-275208 276003-276841 277154-277552 278352-278946 279238-279373 280161-280498 280891-281083 281945-282593 282926-283199 283897-284255 284851-285428 285795-286412 286622-287187 287419-287680 288323-288503 289132-289300 290047-290588 290876-291132 291837-292396 292703-292915 293527-293736 294265-294408 294618-294667 294997-295379 296210-296843 297099-297798 298230-298775 299087-299169 299864-299915 300801-300950 301585-301819 302551-302866 303348-303437 304541-304682 305512-306067 306516-307225 307427-307849 308207-308373 309211-309263 310191-310773 311179-311228 311801-311856 312520-312616 313124-313309 314087-314710 315171-315341 316099-316344 317220-317382 318098-318808 319456-319699 320458-321001 321153-321914 322060-322168 323153-323307 324024-324415 324766-325488 325730-326037 326552-326789 327355-327474 328172-328302 329029-329436 330053-330478 331202-331285 332128-332430 333100-333407 334015-334885 335109-335433 336235-336563 337234-338211 338356-338796 339654-339947 340878-340993 341912-342131 342781-343167 343388-343528 344240-344289 344803-345037 345958-346131 346743-346885 347334-347458 348458-348732 349992-350822 351415-351566 352332-353728 354378-354744 355334-356217 356372-356529 357313-358070 358282-358984 359673-359724 360704-361215 361523-361675 362345-362518 363198-363904 364139-364188 365096-365481 366126-366191 366828-367424 368169-368522 369252-369362 370143-370426 371227-371421 372114-372846 373108-373252 373910-374489 374990-375174 375935-376186 376827-376969 377783-378703 378910-379255 379765-380586 381121-381457 382076-382190 383163-383693 384065-384205 385075-385334 385814-386267 386545-386662 387674-387962 388833-389013 389650-389783 390615-390845 391409-391843 392057-392311 393228-393876 394087-394890 395068-396232 396527-397423 397643-397839 398548-398638 399528-399728 400439-400488 401089-401714 401941-402961 403076-403293 403953-404680 404823-405505 405693-406360 406651-407479 407619-407719 408426-409062 409341-409625 411720-411967 412123-412347 413179-413541 414017-414203 414582-415389 416106-416326 416427-416899 417320-417411 418304-418470 419262-419612 420068-420197 420833-421185 421987-422201 422792-422955 423647-423922 424256-424574 425237-425286 426083-426777 427183-427423 428153-428228 429006-429737 430111-430485 430908-431195 431820-432449 432807-433223 433463-433613 434443-434540 435261-435761 436288-436411 437080-437358 437898-438021 439061-439237 439735-440215 440342-440829 441196-441265 442207-442699 443090-443652 443838-444548 444759-445195 445542-445644 446200-446917 447162-447806 448143-448313 448732-449075 449182-449606 450424-450620 451312-451648 452103-452192 452953-453053 453903-454249 454847-455140 455615-455935 456152-456433 456761-456888 457372-457608 458237-458969 459173-459406 460138-460704 461243-461398 462371-462531 463211-463418 464192-464543 464902-465055 465715-465919 466599-466763 467195-467905 468255-468474 469201-469327 470057-470726 470958-471073 471661-471737 472513-472714 473543-474250 474648-474847 475835-476060 476724-476948 477681-477998 478760-479558 479741-480542 480915-481569 482114-482518 482958-483140 483660-483890 484135-484302 485662-485750 486587-487271 487421-488178 488699-488750 489918-490125 490722-491271 491480-491727 492509-493329 493979-494700 494956-495681 495907-496171 496648-496755 497257-497416 498298-498688 499529-500234 500609-501385 501543-502005 502769-502928 503754-504008 504606-504728 505763-505865 507157-507730 508088-508256 509458-509627 510422-510510 511352-512249 512477-512859 513342-513593 514581-514726 515452-515742 516511-516945 517761-517859 518810-519027 519669-519735 520502-521142 521558-521916 523591-523777 524561-525217 525498-525666 526458-526723 527532-527651 528497-528562 529150-529228 530123-530743 531075-531195 531542-532374 532528-532625 533799-534012 534681-534806 535586-535646 536346-536983 537316-537445 538189-538507 539088-539178 539841-540340 540592-540691 541801-542428 542818-543017 543749-544505 545807-546520 546922-547009 547642-547728 548262-548444 549150-549338 550158-550272 551022-551146 552052-552199 552931-553203 553826-553875 554658-555062 555681-556271 556493-556649 557850-558045 558732-558850 559356-559643 560312-560565 561193-561577 561739-561903 562560-563093 563481-564257 564419-564525 565403-565803 566086-566374 566715-566951 567685-567983 568517-568697 569390-569627 570434-570889 571027-571339 571961-572559 572971-573623 573894-573963 574826-575359 575722-576069 576350-577767 578033-579021 579298-579692 580740-581347 581603-582152 582398-582640 583268-583533 584224-584306 585071-585436 585718-586329 586655-586762 587623-587849 588317-588602 589133-589311 589891-590109 590757-590850 591703-592494 592854-593020 593776-593963 594722-594848 595533-595851 596237-596316 597080-597200 597878-597978 598312-599105 599215-599534 600161-601257 601486-602114 602576-602792 603541-603687 604348-604563 605107-605373 606147-606357 607085-607349 608019-608237 608996-609383 609994-610602 610887-610962 611310-611864 612248-612412 613170-613299 614003-614760 614871-615464 616715-616821 617494-617923 618683-619301 619618-619871 620566-620653 621441-622086 622366-623053 623396-623841 624153-624821 625060-625281 626117-626900 627224-627273 627789-627956 629094-629696 630130-630958 631197-631469 632320-632553 633221-633957 634296-634885 635159-635927 636333-636548 637503-638242 638498-638659 639506-639602 640328-640413 640952-641143 642190-642712 643029-643495 643720-644226 644500-644738 645494-645895 646412-648039 648592-648684 649554-649704 650613-650662 651487-651733 652561-652968 653148-653625 653963-654183 654789-654904 655288-655937 656135-656598 657154-657911 658096-658370 659062-659111 659991-660674 660814-661732 661853-662307 662937-663725 663987-664416 664945-665102 666459-666743 667386-667510 668082-668239 668881-669522 669686-670213 670511-670691 671365-671582 672313-672614 673220-673355 674436-674827 675255-675704 676083-676200 676922-677532 677914-678020 678786-679223 680024-680757 680969-681436 681653-682363 682633-682839 683457-684210 684358-685199 685695-685871 686669-687880 688087-688718 688911-689311 690147-691055 691289-691541 692108-692349 692854-693479 694497-694575 695374-695556 696012-696796 696970-697876 698084-698301 699085-699289 699835-700382 700845-701007 701693-702415 702848-703438 703647-704062 704606-704746 705104-705388 706046-706751 707044-707513 707702-708061 708737-708850 709396-709446 710399-710619 711147-711404 711917-712026 712685-712997 714242-714314 715120-715468;

charset uce_core = 12904-13238 13754-13963 14530-14817 15373-15453 15874-16260 16554-16621 17288-17742 17887-18059 18798-19334 20002-20074 20465-20581 21468-21945 22182-22799 23163-23223 24120-24537 24937-25498 25876-25936 28453-28528 29135-29453 30029-30290 30697-30895 31314-31753 32165-32525 33765-33835 34163-34273 35212-35731 36510-36583 37165-37263 37987-38085 39070-39456 39787-40063 40954-41012 41194-41255 42359-42639 43303-43352 44091-44534 45054-45812 46069-46251 47306-47404 48308-48552 48873-49326 49835-49884 50921-51203 52228-52526 53318-53367 53615-53691 54799-54878 55988-56352 56945-57376 58290-58359 58998-59047 59783-59832 60623-60726 61081-61367 61863-61913 62857-63297 64073-64131 65178-65227 65896-65945 66801-66926 67265-67733 68143-68197 69404-69713 70355-70480 71484-71546 71767-71829 72522-72571 73156-73386 73780-74125 74348-74702 75349-75496 75986-76503 76811-76894 77768-78002 78696-78792 80157-80226 80447-80973 81295-81406 82163-83074 83835-83991 84890-84939 85395-85472 86719-87013 87530-87766 88057-88479 88848-88897 89276-89433 90322-90515 91278-91332 92115-92428 93520-93575 93919-94078 95385-95448 96295-96458 97264-97404 98347-98417 99594-100000 100178-100287 100723-101133 102135-102184 102580-103058 103697-103969 104661-104814 105531-105592 105991-106635 107422-107607 108276-108371 109052-109450 109635-110330 111016-111162 111907-112162 113041-113110 113520-114102 114737-114974 116279-116328 117098-117216 117671-117720 118507-118559 119469-119523 119954-120321 121024-121073 122556-122616 124196-124272 124585-124648 126033-126083 126827-127025 127847-128303 128996-129316 129617-130039 131575-131739 132070-132544 132849-133006 134076-134517 134892-135006 135838-135971 136608-137143 137825-138057 139111-139217 139606-139982 140675-141218 141663-141781 142736-142785 143096-143343 144215-144465 144632-144712 145353-145668 146520-146569 147741-147973 148872-149378 149845-150224 150927-150979 151261-151610 152022-152321 152598-152925 153993-154090 154669-154720 155202-155363 156313-156661 157151-157222 158292-158349 159355-159410 160753-160806 161184-161470 162909-162960 163466-163881 164517-164877 165469-165859 166345-166944 167312-167604 168337-168391 169437-169649 170356-170451 170660-171259 172460-172539 172937-173351 174465-174530 174790-175240 175509-175568 176608-177142 177691-177857 178080-178474 178932-179300 179680-180031 180709-180941 181073-181131 181749-181913 183168-183217 183761-184126 185057-185242 185820-186120 186927-187230 188087-188301 188476-188644 190193-190242 190822-191076 192053-192323 192503-192562 193060-193113 194263-194333 195178-195337 195584-195786 196578-197078 198038-198099 198331-198382 199118-199330 199807-200235 201274-201424 201720-201865 203232-203283 204174-204226 204584-204657 206402-206463 206901-206951 207485-207659 209139-209189 210014-210182 210708-211129 211862-211924 212963-213012 213794-213843 214341-214430 215257-215581 215896-217184 217701-218131 218548-218664 219332-219427 220231-220442 220847-221047 221721-221889 223641-224218 225140-225263 226146-226235 226533-226984 227379-227465 228233-228315 229237-229472 229877-229930 231045-231094 231673-231841 232371-232441 233927-233981 234737-234786 235440-235489 236692-236844 237227-237297 238016-238082 239504-239554 239880-239979 240930-241268 242074-242396 243223-243321 243658-244256 245316-245497 245792-246231 246567-246640 247232-247295 248712-248794 249325-250309 250607-250960 251655-251704 252455-252915 253979-254028 254331-254386 255321-255522 256408-256673 257743-257793 258252-258614 259207-259468 260178-260231 260826-261205 261408-261541 262011-262406 263144-263264 264302-264351 265007-265999 266502-266573 267492-267561 268015-268266 269628-269680 270599-270716 271315-271367 271859-272164 272916-272980 273176-273225 273528-273623 274641-274713 275209-275952 276842-276955 277553-277647 278947-279063 279374-279428 280499-280570 281084-281360 282594-282687 283200-283332 284256-284646 285429-285486 286413-286470 287188-287348 287681-287737 288504-288662 289301-289388 290589-290650 291133-291186 292397-292522 292916-292973 293737-293859 294409-294560 294668-294851 295380-295436 296844-296893 297799-297956 298776-298837 299170-299813 299916-300438 300951-301478 301820-302241 302867-302916 303438-303635 304683-305190 306068-306382 307226-307321 307850-308097 308374-308888 309264-309364 310774-310847 311229-311368 311857-312314 312617-313053 313310-313850 314711-314985 315342-315417 316345-316395 317383-317815 318809-319118 319700-319839 321002-321061 321915-321976 322169-322279 323308-323741 324416-324715 325489-325559 326038-326299 326790-327098 327475-327690 328303-328537 329437-329905 330479-330878 331286-331512 332431-332480 333408-333457 334886-334962 335434-335817 336564-336983 338212-338262 338797-338898 339948-340568 340994-341044 342132-342218 343168-343245 343529-343600 344290-344614 345038-345087 346132-346253 346886-347245 347459-347642 348733-348860 350823-350885 351567-352174 353729-353778 354745-355104 356218-356302 356530-356579 358071-358191 358985-359450 359725-360429 361216-361458 361676-361785 362519-362568 363905-363970 364189-364915 365482-365834 366192-366560 367425-367907 368523-368611 369363-369437 370427-370487 371422-372046 372847-372903 373253-373334 374490-374542 375175-375276 376187-376776 376970-377026 378704-378858 379256-379532 380587-380772 381458-381863 382191-382885 383694-383776 384206-384281 385335-385728 386268-386338 386663-386798 387963-388105 389014-389493 389784-390238 390846-390895 391844-391893 392312-393049 393877-394014 394891-394970 396233-396307 397424-397482 397840-398036 398639-398688 399729-399820 400489-400548 401715-401877 402962-403024 403294-403660 404681-404751 405506-405582 406361-406493 407480-407556 407720-407936 409063-409134 409626-411472 411968-412017 412348-412461 413542-413856 414204-414513 415390-415985 416327-416376 416900-416955 417412-417470 418471-418947 419613-419754 420198-420647 421186-421263 422202-422511 422956-423429 423923-423974 424575-424628 425287-425490 426778-427112 427424-427480 428229-428289 429738-429787 430486-430855 431196-431245 432450-432652 433224-433281 433614-433700 434541-435133 435762-436097 436412-436478 437359-437636 438022-438551 439238-439569 440216-440273 440830-440997 441266-441411 442700-442869 443653-443710 444549-444643 445196-445251 445645-445902 446918-447017 447807-447889 448314-448371 449076-449131 449607-449669 450621-451046 451649-451896 452193-452264 453054-453350 454250-454591 455141-455194 455936-455988 456434-456499 456889-457297 457609-457900 458970-459072 459407-459499 460705-460771 461399-461579 462532-462666 463419-463476 464544-464729 465056-465105 465920-466171 466764-466813 467906-468042 468475-468974 469328-469511 470727-470791 471074-471151 471738-471857 472715-473402 474251-474358 474848-474926 476061-476118 476949-477542 477999-478307 479559-479645 480543-480692 481570-481636 482519-482568 483141-483195 483891-483940 484303-484447 485751-485802 487272-487321 488179-488524 488751-489094 490126-490175 491272-491322 491728-491777 493330-493696 494701-494754 495682-495742 496172-496229 496756-496861 497417-497488 498689-499088 500235-500307 501386-501446 502006-502191 502929-503137 504009-504524 504729-504825 505866-506980 507731-507878 508257-509114 509628-509688 510511-510847 512250-512422 512860-512909 513594-513644 514727-515165 515743-516155 516946-517131 517860-518226 519028-519468 519736-520158 521143-521437 521917-523153 523778-523935 525218-525314 525667-526339 526724-526852 527652-527810 528563-528677 529229-529497 530744-530840 531196-531390 532375-532424 532626-532675 534013-534206 534807-535240 535647-535928 536984-537083 537446-538133 538508-538573 539179-539790 540341-540444 540692-541344 542429-542578 543018-543504 544506-544671 546521-546571 547010-547087 547729-548211 548445-548812 549339-549959 550273-550849 551147-551506 552200-552274 553204-553254 553876-554022 555063-555452 556272-556398 556650-556747 558046-558638 558851-559261 559644-559968 560566-560961 561578-561640 561904-562342 563094-563262 564258-564307 564526-564659 565804-565889 566375-566664 566952-567025 567984-568035 568698-569263 569628-569804 570890-570959 571340-571414 572560-572814 573624-573721 573964-574226 575360-575411 576070-576228 577768-577833 579022-579094 579693-580523 581348-581397 582153-582202 582641-583153 583534-583999 584307-584890 585437-585505 586330-586399 586763-587310 587850-588149 588603-589014 589312-589558 590110-590224 590851-590954 592495-592572 593021-593459 593964-594477 594849-595477 595852-596167 596317-596445 597201-597571 597979-598055 599106-599164 599535-600006 601258-601317 602115-602451 602793-602844 603688-604269 604564-604619 605374-605637 606358-606451 607350-607686 608238-608702 609384-609438 610603-610752 610963-611040 611865-612197 612413-612856 613300-613436 614761-614817 615465-616550 616822-617053 617924-617973 619302-619351 619872-620366 620654-620949 622087-622155 623054-623105 623842-623902 624822-624960 625282-625903 626901-627130 627274-627738 627957-628825 629697-629768 630959-631142 631470-631603 632554-632904 633958-634039 634886-634960 635928-636049 636549-636808 638243-638295 638660-639301 639603-639806 640414-640762 641144-641221 642713-642841 643496-643587 644227-644413 644739-644794 645896-646273 648040-648427 648685-648912 649705-650148 650663-650720 651734-652155 652969-653038 653626-653853 654184-654245 654905-655012 655938-655997 656599-656993 657912-657965 658371-658628 659112-659936 660675-660733 661733-661802 662308-662360 663726-663804 664417-664696 665103-665313 666744-667082 667511-667922 668240-668826 669523-669597 670214-670288 670692-670742 671583-671632 672615-673129 673356-674301 674828-675150 675705-675838 676201-676771 677533-677718 678021-678162 679224-679537 680758-680862 681437-681486 682364-682551 682840-683353 684211-684279 685200-685526 685872-685967 687881-687966 688719-688771 689312-689737 691056-691118 691542-691783 692350-692399 693480-694348 694576-694657 695557-695677 696797-696865 697877-697959 698302-698974 699290-699628 700383-700703 701008-701080 702416-702500 703439-703503 704063-704376 704747-704810 705389-705438 706752-706859 707514-707588 708062-708623 708851-709013 709447-709600 710620-711040 711405-711817 712027-712489 712998-713197 714315-714875 715469-715644;

charset uce_right = 13239-13545 13964-14215 14818-15141 15454-15804 16261-16310 16622-17237 17743-17792 18060-18738 19335-19515 20075-20270 20582-21225 21946-22131 22800-22857 23224-24069 24538-24880 25499-25705 25937-27254 28529-28824 29454-29803 30291-30484 30896-31213 31754-31917 32526-32855 33836-34059 34274-35131 35732-35926 36584-36907 37264-37870 38086-38877 39457-39519 40064-40372 41013-41121 41256-41974 42640-43102 43353-43908 44535-44939 45813-45892 46252-46679 47405-47683 48553-48701 49327-49634 49885-50609 51204-51264 52527-52641 53368-53504 53692-54512 54879-55493 56353-56676 57377-57566 58360-58442 59048-59453 59833-60063 60727-60776 61368-61770 61914-62646 63298-63455 64132-64412 65228-65376 65946-66230 66927-67048 67734-67822 68198-68702 69714-69768 70481-70626 71547-71620 71830-72226 72572-73097 73387-73493 74126-74207 74703-74910 75497-75643 76504-76611 76895-77565 78003-78345 78793-79639 80227-80393 80974-81182 81407-82030 83075-83248 83992-84272 84940-85257 85473-86298 87014-87172 87767-87965 88480-88664 88898-89059 89434-90015 90516-91036 91333-91870 92429-92805 93576-93776 94079-95315 95449-96108 96459-97173 97405-98192 98418-99229 100001-100050 100288-100630 101134-101646 102185-102445 103059-103162 103970-104052 104815-104934 105593-105913 106636-106730 107608-107659 108372-108450 109451-109501 110331-110397 111163-111259 112163-112274 113111-113323 114103-114390 114975-115421 116329-116421 117217-117352 117721-118212 118560-118938 119524-119802 120322-120684 121074-121790 122617-123079 124273-124493 124649-125579 126084-126156 127026-127562 128304-128472 129317-129430 130040-130325 131740-131789 132545-132696 133007-133704 134518-134713 135007-135622 135972-136464 137144-137394 138058-138383 139218-139269 139983-140265 141219-141600 141782-142126 142786-143036 143344-143961 144466-144571 144713-145282 145669-146349 146570-147455 147974-148414 149379-149432 150225-150321 150980-151085 151611-151902 152322-152505 152926-153215 154091-154260 154721-155043 155364-156253 156662-157042 157223-158026 158350-158924 159411-160157 160807-161106 161471-161983 162961-163127 163882-164305 164878-165418 165860-166102 166945-167077 167605-167966 168392-168824 169650-169708 170452-170580 171260-171535 172540-172725 173352-173707 174531-174702 175241-175331 175569-176487 177143-177197 177858-177919 178475-178762 179301-179435 180032-180235 180942-181016 181132-181639 181914-182550 183218-183437 184127-184468 185243-185598 186121-186377 187231-187410 188302-188379 188645-189246 190243-190348 191077-191359 192324-192434 192563-193009 193114-193596 194334-194737 195338-195495 195787-196526 197079-197205 198100-198221 198383-198683 199331-199576 200236-200465 201425-201556 201866-202598 203284-203691 204227-204454 204658-205477 206464-206695 206952-207434 207660-208090 209190-209287 210183-210306 211130-211388 211925-212246 213013-213175 213844-214219 214431-214967 215582-215744 217185-217424 218132-218433 218665-219254 219428-220116 220443-220795 221048-221603 221890-223289 224219-224633 225264-225447 226236-226433 226985-227198 227466-228038 228316-228889 229473-229581 229931-230533 231095-231395 231842-232217 232442-233614 233982-234142 234787-235102 235490-236082 236845-237072 237298-237776 238083-238727 239555-239704 239980-240574 241269-241595 242397-242490 243322-243424 244257-244547 245498-245558 246232-246392 246641-247089 247296-247838 248795-249015 250310-250440 250961-251304 251705-252301 252916-253235 254029-254218 254387-255057 255523-256012 256674-256990 257794-257939 258615-258838 259469-259750 260232-260360 261206-261350 261542-261822 262407-262665 263265-263880 264352-264674 266000-266272 266574-266920 267562-267815 268267-268811 269681-269811 270717-270889 271368-271640 272165-272380 272981-273066 273226-273471 273624-274388 274714-275133 275953-276002 276956-277153 277648-278351 279064-279237 279429-280160 280571-280890 281361-281944 282688-282925 283333-283896 284647-284850 285487-285794 286471-286621 287349-287418 287738-288322 288663-289131 289389-290046 290651-290875 291187-291836 292523-292702 292974-293526 293860-294264 294561-294617 294852-294996 295437-296209 296894-297098 297957-298229 298838-299086 299814-299863 300439-300800 301479-301584 302242-302550 302917-303347 303636-304540 305191-305511 306383-306515 307322-307426 308098-308206 308889-309210 309365-310190 310848-311178 311369-311800 312315-312519 313054-313123 313851-314086 314986-315170 315418-316098 316396-317219 317816-318097 319119-319455 319840-320457 321062-321152 321977-322059 322280-323152 323742-324023 324716-324765 325560-325729 326300-326551 327099-327354 327691-328171 328538-329028 329906-330052 330879-331201 331513-332127 332481-333099 333458-334014 334963-335108 335818-336234 336984-337233 338263-338355 338899-339653 340569-340877 341045-341911 342219-342780 343246-343387 343601-344239 344615-344802 345088-345957 346254-346742 347246-347333 347643-348457 348861-349991 350886-351414 352175-352331 353779-354377 355105-355333 356303-356371 356580-357312 358192-358281 359451-359672 360430-360703 361459-361522 361786-362344 362569-363197 363971-364138 364916-365095 365835-366125 366561-366827 367908-368168 368612-369251 369438-370142 370488-371226 372047-372113 372904-373107 373335-373909 374543-374989 375277-375934 376777-376826 377027-377782 378859-378909 379533-379764 380773-381120 381864-382075 382886-383162 383777-384064 384282-385074 385729-385813 386339-386544 386799-387673 388106-388832 389494-389649 390239-390614 390896-391408 391894-392056 393050-393227 394015-394086 394971-395067 396308-396526 397483-397642 398037-398547 398689-399527 399821-400438 400549-401088 401878-401940 403025-403075 403661-403952 404752-404822 405583-405692 406494-406650 407557-407618 407937-408425 409135-409340 411473-411719 412018-412122 412462-413178 413857-414016 414514-414581 415986-416105 416377-416426 416956-417319 417471-418303 418948-419261 419755-420067 420648-420832 421264-421986 422512-422791 423430-423646 423975-424255 424629-425236 425491-426082 427113-427182 427481-428152 428290-429005 429788-430110 430856-430907 431246-431819 432653-432806 433282-433462 433701-434442 435134-435260 436098-436287 436479-437079 437637-437897 438552-439060 439570-439734 440274-440341 440998-441195 441412-442206 442870-443089 443711-443837 444644-444758 445252-445541 445903-446199 447018-447161 447890-448142 448372-448731 449132-449181 449670-450423 451047-451311 451897-452102 452265-452952 453351-453902 454592-454846 455195-455614 455989-456151 456500-456760 457298-457371 457901-458236 459073-459172 459500-460137 460772-461242 461580-462370 462667-463210 463477-464191 464730-464901 465106-465714 466172-466598 466814-467194 468043-468254 468975-469200 469512-470056 470792-470957 471152-471660 471858-472512 473403-473542 474359-474647 474927-475834 476119-476723 477543-477680 478308-478759 479646-479740 480693-480914 481637-482113 482569-482957 483196-483659 483941-484134 484448-485661 485803-486586 487322-487420 488525-488698 489095-489917 490176-490721 491323-491479 491778-492508 493697-493978 494755-494955 495743-495906 496230-496647 496862-497256 497489-498297 499089-499528 500308-500608 501447-501542 502192-502768 503138-503753 504525-504605 504826-505762 506981-507156 507879-508087 509115-509457 509689-510421 510848-511351 512423-512476 512910-513341 513645-514580 515166-515451 516156-516510 517132-517760 518227-518809 519469-519668 520159-520501 521438-521557 523154-523590 523936-524560 525315-525497 526340-526457 526853-527531 527811-528496 528678-529149 529498-530122 530841-531074 531391-531541 532425-532527 532676-533798 534207-534680 535241-535585 535929-536345 537084-537315 538134-538188 538574-539087 539791-539840 540445-540591 541345-541800 542579-542817 543505-543748 544672-545806 546572-546921 547088-547641 548212-548261 548813-549149 549960-550157 550850-551021 551507-552051 552275-552930 553255-553825 554023-554657 555453-555680 556399-556492 556748-557849 558639-558731 559262-559355 559969-560311 560962-561192 561641-561738 562343-562559 563263-563480 564308-564418 564660-565402 565890-566085 566665-566714 567026-567684 568036-568516 569264-569389 569805-570433 570960-571026 571415-571960 572815-572970 573722-573893 574227-574825 575412-575721 576229-576349 577834-578032 579095-579297 580524-580739 581398-581602 582203-582397 583154-583267 584000-584223 584891-585070 585506-585717 586400-586654 587311-587622 588150-588316 589015-589132 589559-589890 590225-590756 590955-591702 592573-592853 593460-593775 594478-594721 595478-595532 596168-596236 596446-597079 597572-597877 598056-598311 599165-599214 600007-600160 601318-601485 602452-602575 602845-603540 604270-604347 604620-605106 605638-606146 606452-607084 607687-608018 608703-608995 609439-609993 610753-610886 611041-611309 612198-612247 612857-613169 613437-614002 614818-614870 616551-616714 617054-617493 617974-618682 619352-619617 620367-620565 620950-621440 622156-622365 623106-623395 623903-624152 624961-625059 625904-626116 627131-627223 627739-627788 628826-629093 629769-630129 631143-631196 631604-632319 632905-633220 634040-634295 634961-635158 636050-636332 636809-637502 638296-638497 639302-639505 639807-640327 640763-640951 641222-642189 642842-643028 643588-643719 644414-644499 644795-645493 646274-646411 648428-648591 648913-649553 650149-650612 650721-651486 652156-652560 653039-653147 653854-653962 654246-654788 655013-655287 655998-656134 656994-657153 657966-658095 658629-659061 659937-659990 660734-660813 661803-661852 662361-662936 663805-663986 664697-664944 665314-666458 667083-667385 667923-668081 668827-668880 669598-669685 670289-670510 670743-671364 671633-672312 673130-673219 674302-674435 675151-675254 675839-676082 676772-676921 677719-677913 678163-678785 679538-680023 680863-680968 681487-681652 682552-682632 683354-683456 684280-684357 685527-685694 685968-686668 687967-688086 688772-688910 689738-690146 691119-691288 691784-692107 692400-692853 694349-694496 694658-695373 695678-696011 696866-696969 697960-698083 698975-699084 699629-699834 700704-700844 701081-701692 702501-702847 703504-703646 704377-704605 704811-705103 705439-706045 706860-707043 707589-707701 708624-708736 709014-709395 709601-710398 711041-711146 711818-711916 712490-712684 713198-714241 714876-715119 715645-715871;

charpartition mymodels =

TPM3u+F+R3: ATP6_1_ATP8_1_ND2_1,

GTR+F+R3: ATP6_2_ATP8_2_ND2_2_ND5_2,

TN+F+R4: ATP6_3_ATP8_3_ND2_3,

TIM3e+I+G4: COX1_1_COX3_1_CYTB_1,

F81+F: COX1_2,

TN+F+R3: COX1_3_COX2_3,

TIM3e+G4: COX2_1_ND4L_1,

TPM3u+F+R2: COX2_2_COX3_2_CYTB_2_ND1_2_ND3_2_ND4_2_ND4L_2,

TIM3+F+R4: COX3_3_CYTB_3_ND4L_3,

TIM3+F+I+G4: ND1_1_ND3_1_ND4_1,

TN+F+R4: ND1_3_ND3_3_ND4_3_ND5_3,

TIM2+F+R3: ND5_1_rrnL_rrnS,

TIM2+F+I+G4: ND6_1,

TVM+F+G4: ND6_2,

TN+F+ASC+G4: ND6_3,

TVM+F+R3: uce_left,

TVMe+R3: uce_core,

TVM+F+R3: uce_right;

end;
